# α-Synuclein Conformations in Plasma Distinguish Parkinson’s Disease from Dementia with Lewy Bodies

**DOI:** 10.1101/2024.05.07.593056

**Authors:** George T. Kannarkat, Rebecca Zack, R. Tyler Skrinak, James F. Morley, Roseanne Davila-Rivera, Sanaz Arezoumandan, Katherine Dorfmann, Kelvin Luk, David A. Wolk, Daniel Weintraub, Thomas F. Tropea, Edward B. Lee, Sharon X. Xie, Ganesh Chandrasekaran, Virginia M.-Y. Lee, David Irwin, Rizwan S. Akhtar, Alice S. Chen-Plotkin

**Affiliations:** Department of Neurology, Perelman School of Medicine, University of Pennsylvania; Philadelphia, PA, USA, 19104; Parkinson’s Disease Research, Education and Clinical Center, Corporal Michael J. Crescenz VA Medical Center; Philadelphia, PA, USA, 19104; Department of Pathology and Laboratory Medicine, Perelman School of Medicine, University of Pennsylvania; Philadelphia, PA, USA, 19104; Department of Psychiatry and Neurology, Perelman School of Medicine, University of Pennsylvania; Philadelphia, PA, USA, 19104; Department of Biostatistics, Epidemiology and Informatics, Perelman School of Medicine, University of Pennsylvania; Philadelphia, PA, USA, 19104; Ken and Ruth Davee Department of Neurology and Simpson Querrey Center for Neurogenetics, Northwestern University Feinberg School of Medicine, Chicago, IL, USA, 60611

**Author notes:** co-first authors with equal contribution (George T. Kannarkat, Rebecca Zack). **One Sentence Summary:** Antibodies against distinct plasma conformations of α-synuclein distinguish Parkinson’s disease from dementia with Lewy bodies and predict cognitive decline in Parkinson’s disease.

## Abstract

Spread and aggregation of misfolded α-synuclein (aSyn) within the brain is the pathologic hallmark of Lewy body diseases (LBD), including Parkinson’s disease (PD) and dementia with Lewy bodies (DLB). While evidence exists for multiple aSyn protein conformations, often termed “strains” for their distinct biological properties, it is unclear whether PD and DLB result from aSyn strain differences, and biomarkers that differentiate PD and DLB are lacking. Moreover, while pathological forms of aSyn have been detected outside the brain (*e.g.,* in skin, gut, blood), the functional significance of these peripheral aSyn species is unclear. Here, we developed assays using monoclonal antibodies selective for two different aSyn species generated *in vitro* – termed Strain A and Strain B – and used them to evaluate human brain tissue, cerebrospinal fluid (CSF), and plasma, through immunohistochemistry, enzyme-linked immunoassay, and immunoblotting. Surprisingly, we found that plasma aSyn species detected by these antibodies differentiated individuals with PD vs. DLB in a discovery cohort (UPenn, n=235, AUC 0.83) and a multi-site replication cohort (Parkinson’s Disease Biomarker Program, or PDBP, n=200, AUC 0.72). aSyn plasma species detected by the Strain A antibody also predicted rate of cognitive decline in PD. We found no evidence for aSyn strains in CSF, and ability to template aSyn fibrillization differed for species isolated from plasma vs. brain, and in PD vs. DLB. Taken together, our findings suggest that aSyn conformational differences may impact clinical presentation and cortical spread of pathological aSyn. Moreover, the enrichment of these aSyn strains in plasma implicates a non-central nervous system source.

## INTRODUCTION

Lewy body diseases (LBD) are chronic neurodegenerative conditions that are characterized by intraneuronal aggregates of misfolded α-synuclein (aSyn), called Lewy bodies (LBs) and Lewy neurites, collectively Lewy pathology, which affect over 10 million people worldwide with increasing incidence (*1–7*). Two clinical disorders, Parkinson’s disease (PD) and dementia with Lewy bodies (DLB), fall within this pathologic classification (*6, 8*). PD is diagnosed by its motor symptoms of bradykinesia, tremor, and rigidity (*9, 10*). PD has numerous non-motor symptoms including sleep changes, constipation, hyposmia, depression, and anxiety. However, cognitive impairment, characterized as mild (PD-MCI) or progressing to dementia (PDD) and occurring in over 70% of PD individuals throughout the disease course, may be the most detrimental to quality of life (*11–15*).

DLB is distinguished clinically from PD based on the emergence of dementia prior to or within one year of motor symptoms (*16*). At autopsy, PDD and DLB are virtually indistinguishable, with diffuse Lewy pathology in limbic and cortical areas, along with comorbid Alzheimer’s disease (AD) pathology in up to half of individuals (*6, 8, 11, 17, 18*). Why such similar pathology manifests so differently during life is not understood, and there are no diagnostic biomarkers that distinguish PD from DLB (*11, 19, 20*). Because disease progression, and specifically cognitive impairment, correlates with spread of Lewy pathology from brainstem to neocortical areas, understanding the molecular features of aSyn that impact its rate of misfolding and spread within the brain is of great importance (*18, 21, 22*). Moreover, this understanding may be key to developing novel therapies for LBD (*6, 17, 18*).

The biochemical context, such as increased aSyn concentration or alterations in its lipid binding, may promote misfolding of monomeric aSyn, leading to the formation of oligomers and fibrils (*6, 7, 23, 24*). Certain aggregated forms of aSyn cause cellular toxicity, trigger inflammatory responses, and further promote aggregation of aSyn monomer in a prion-like manner (*5–7, 24, 25*). Increasing evidence demonstrates that specific aSyn conformations may act as “strains”, with distinct and reproducible biological behaviors, impacting clinical phenotype. For example, structural, *in vitro*, and *in vivo* studies suggest that aSyn “strains” differ for LBD vs. multiple systems atrophy (MSA), a degenerative disorder characterized by glial cytoplasmic aSyn inclusions (*25–34*). While different aSyn strains have been generated and extensively studied *in vitro*, the development of conformation-specific aSyn antibodies allows for the investigation of distinct *in vivo* aSyn species (*35–37*). Given the potential for different aSyn strains to associate with disease-specific characteristics, we sought to determine whether antibodies generated against *in vitro* aSyn strains can differentially recognize aSyn species in human biofluids. We further asked whether aSyn species detected by these strain-selective antibodies may act as diagnostic and prognostic biomarkers in LBD.

## RESULTS

### Antibodies selective for aSyn strains detect differing brain pathology

As previously described (*35, 36*), the anti-aSyn monoclonal antibodies, 7015 and 9027, were isolated via immunization of mice with two artificially-generated aSyn strains, termed Strain A (7015) and Strain B (9027), respectively (Fig. 1A). These two strains were created through serial passaging of recombinant forms of pathological aSyn in cells, with Strain B demonstrating an emergent ability to induce phosphorylated tau (p-tau) pathology in primary mouse hippocampal neurons (*35–37*). The 7015 antibody epitope is discontinuous, including a region around residue 50 and at the extreme C-terminus of aSyn as previously described (*36*), while the 9027 epitope is continuous between residues 120-140 (Fig. S1).

**Figure 1.**
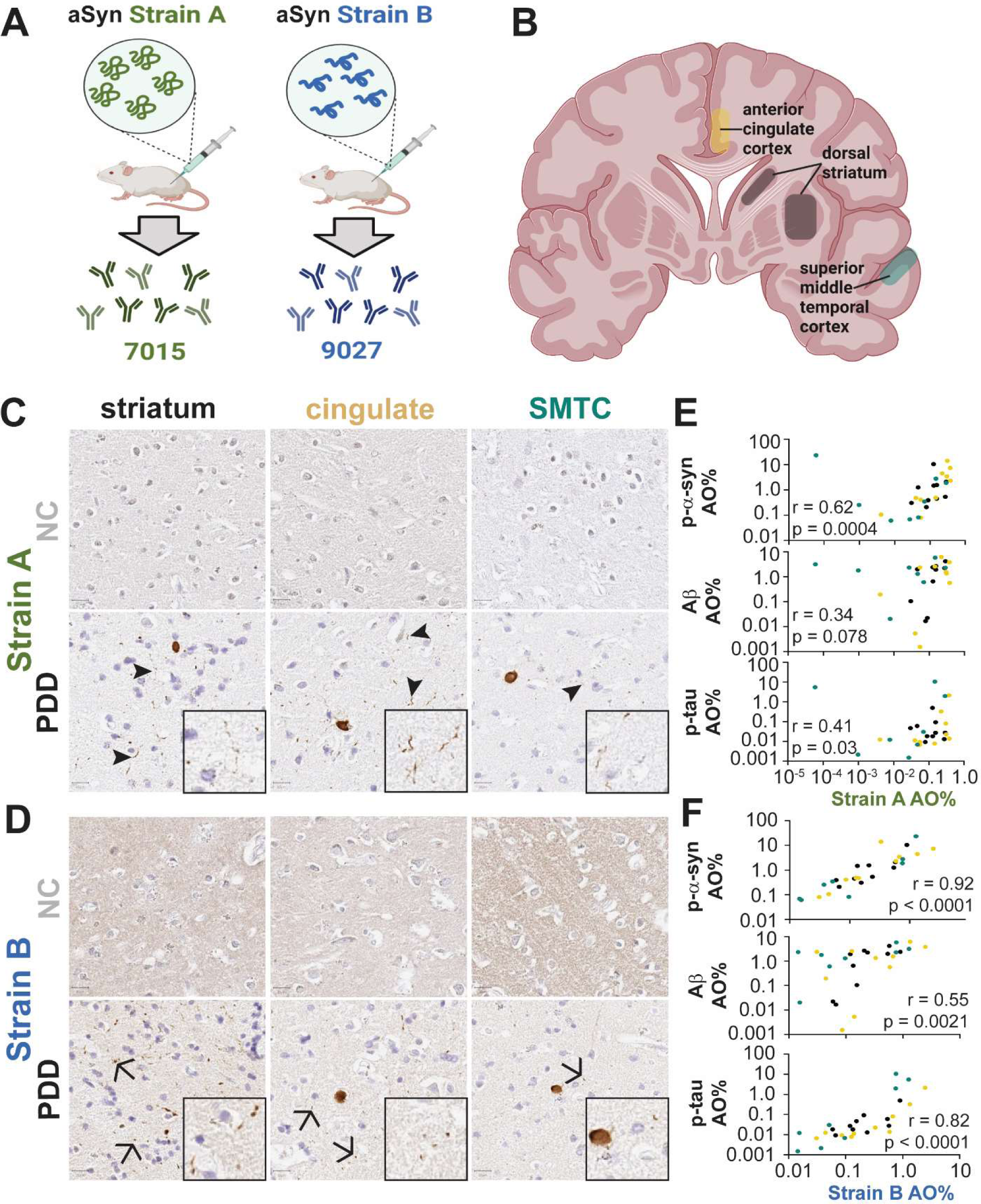
Strain A (7015) and Strain B (9027) antibodies selectively and differentially recognize aSyn pathology in human brain tissue. Mice were immunized with two different *in vitro* generated strains of aSyn to produce 7015 (Strain A) and 9027 (Strain B) monoclonal antibodies (**A**). Brain regions (**B**) known to associate with diffuse spread of Lewy pathology from ten NC and ten individuals with LBD were immunostained with Strain A (**C**) and Strain B (**D**) antibodies with representative images showing preferential recognition of thread-like, “neuritic”, (arrowheads) vs. dot-like, “aggregate”, pathology (arrows), respectively. Compared to Strain A-associated pathology (**E**), Strain B-associated aSyn pathology (**F**) more strongly correlated with p-aSyn, amyloid, and p-tau staining across all three examined brain regions. P-value and r statistic for Pearson correlation reported in **E-F**. AO = area occupied, PDD = Parkinson’s disease dementia, NC = normal control, p-aSyn = phosphorylated aSyn, p-tau = phosphorylated tau, SMTC = superior middle temporal cortex.

We first confirmed that these antibodies recognize aSyn species in human brain by performing immunohistochemistry of striatum, temporal cortex, and cingulate cortex from ten individuals with LBD (Fig. 1B, S2A-B; Table S1). Brain regions were selected based on the pattern of spread of Lewy pathology proposed in the latter stages of the Braak hypothesis (striatum to temporal cortex to cingulate cortex) which parallels disease progression and cognitive decline (*38*). In all three brain regions, Strain A antibody preferentially recognized forms of aSyn associated with neuronal processes (“neuritic” pattern), while Strain B antibody recognized aSyn in LBs as well as a more dot-like pattern (Fig. 1C-D). The staining pattern for both strain antibodies reflected a subset of total aSyn pathology in both PD and DLB (Fig. S3A). Furthermore, the aSyn species detected by the Strain B antibody was more strongly correlated with phosphorylated aSyn (p-aSyn r = 0.92 vs. r = 0.62 for Strain A), amyloid β (Aβ; r = 0.55 vs. r = 0.34 for Strain A), and p-tau (r = 0.82 vs. r = 0.41 for Strain A) staining across all evaluated brain regions (Fig. 1E-F, Fig. S3B). Both antibodies also detected aSyn pathology in multiple system atrophy (MSA) cases, with Strain A recognizing smaller glial cytoplasmic inclusions relative to Strain B (Fig. S2C, Table S2). These data confirm prior reports that Strain A and Strain B antibodies recognize different aSyn species in human brain. They also demonstrate that the Strain B antibody recognizes a form of aSyn that is more correlated with AD co-pathology in human brain (*35, 36*).

### Plasma aSyn strain levels do not reflect CSF or brain levels

Having confirmed that Strain A and Strain B antibodies detect aSyn species in brain tissue with aSyn pathology, we next tested whether we could use these strain-selective antibodies to detect aSyn conformations in biofluids and tissue lysates by ELISA. We established two sandwich ELISAs using an antibody recognizing total aSyn (MJFR1 antibody) for capture and biotinylated Strain A or Strain B antibodies for detection. Differences in these strain ELISAs were reflected in their differential detection of even artificial substrates: the Strain A assay preferentially detected recombinant aSyn pre-formed fibrils (PFFs) over aSyn monomer, but this was not true of the Strain B assay (Fig. 2A). Moreover, in human brain lysates from normal controls (NC), individuals with PD (iwPD), and individuals with DLB (iwDLB, Table S3), both the Strain A and Strain B antibodies detected high-molecular weight (HMW) aSyn species in iwPD and iwDLB by immunoblotting (Fig. 2B-C); these HMW species were not detected in NC brain lysates, or using secondary antibody alone (Fig. S4A). At an individual patient level, Strain A ELISA measures from caudate (Fig. 2E) corresponded well with quantities of HMW species detected by immunoblotting (Fig. 2B), with higher Strain A levels found in iwPD and iwDLB. Finally, Strain A ELISA measures from cerebellum (Fig. S4B), a brain region with little aSyn pathology even in iwPD or iwDLB, differed minimally comparing NC, iwPD, and iwDLB. These studies further confirm that our strain-selective antibodies detect pathological species of aSyn, while also validating them for use in biofluids and tissue lysates.

**Figure 2.**
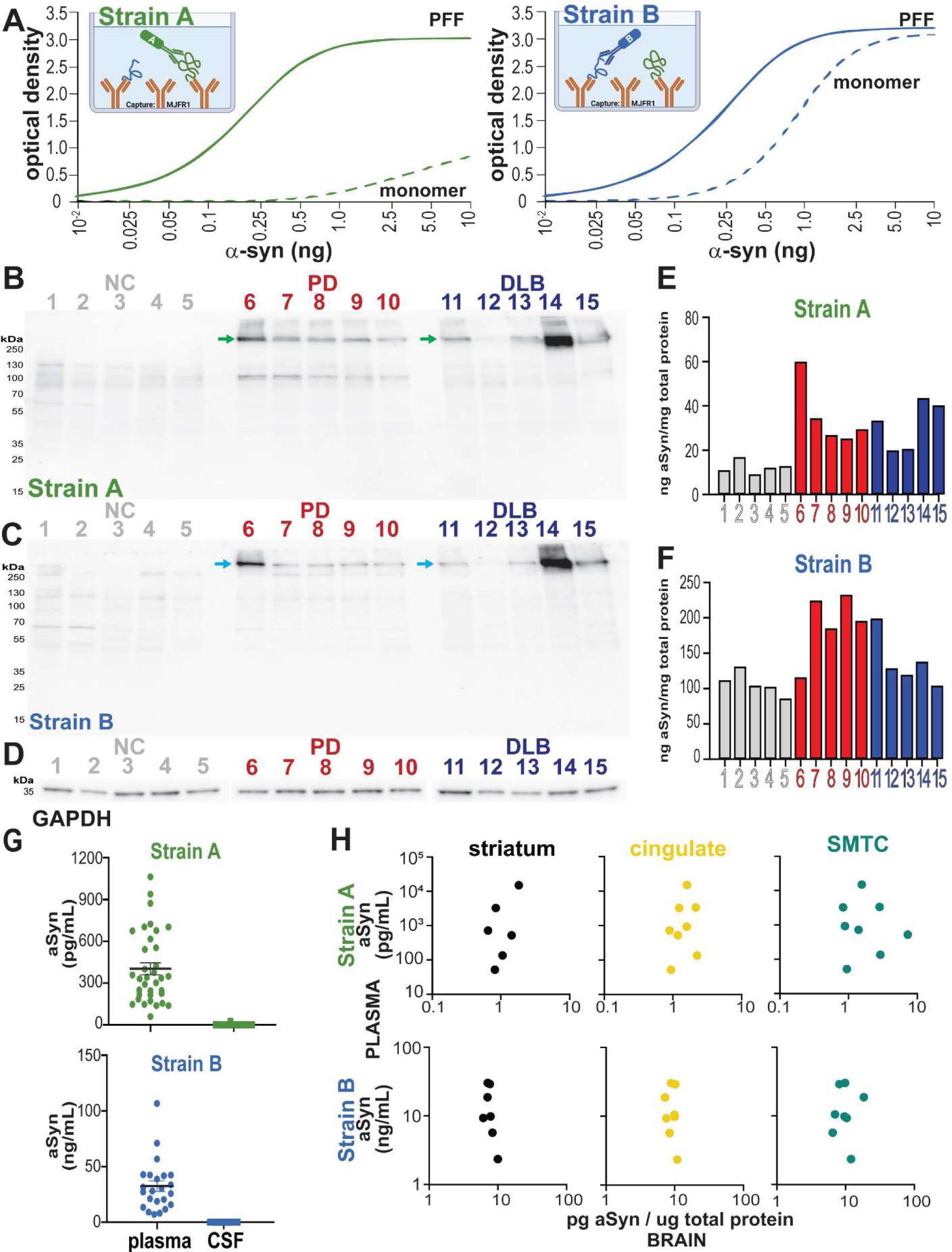
aSyn strains can be detected by ELISA and are uniquely enriched in plasma over cerebrospinal fluid. Sandwich ELISAs using MJFR1 (total aSyn) antibody for capture and Strain A or Strain B biotinylated antibodies for detection show differential selectivity for recombinant a-syn preformed fibrils (PFF) vs. monomer (**A**). Western blot (WB) with Strain A (**B**) and with Strain B (**C)** antibodies detect high molecular weight (HMW, arrows) aSyn species only in individuals with PD or DLB. GAPDH is displayed as a loading control (**D**). Strain A aSyn levels measured by ELISA (**E**) correspond at the individual level with levels of HMW aSyn species detected by immunoblot with Strain A antibody (**B**), while Strain B aSyn levels measured by ELISA (**F**) show less individual-level correspondence to quantities detected by immunoblot (**C**). Strain A and B aSyn species are not readily detectable in CSF (n = 44 individuals tested), but robustly detected in plasma (**G**). For matched brain and plasma samples (plasma collected within two years of autopsy for n=10 individuals), plasma aSyn Strain levels do not correlate with brain aSyn levels (RIPA extracts shown, **H**). NC = normal controls, PD = Parkinson’s disease, RIPA = radioimmunoprecipitation assay, SMTC = superior middle temporal cortex.

We then used our aSyn strain ELISAs to measure levels of these aSyn species in CSF from LBD individuals (n = 44, Table S4). Strain A aSyn species were detectable in only 3 individuals at very low levels, and Strain B species were not detectable in any individual in the CSF (Fig. 2G). These findings were also replicated in pooled CSF from LBD individuals, with serial dilution in assay buffer to exclude matrix effect (data not shown). Surprisingly, however, the same 44 LBD individuals from whom we had tested CSF samples had robustly detectable levels of both Strain A and Strain B aSyn species in the plasma (Fig. 2G).

To understand whether plasma aSyn species reflected brain aSyn species, we measured blood and brain levels using each aSyn strain ELISA in individuals with LBD (n = 10) who had plasma collected within 2 years of death (Table S1), with an average plasma collection-autopsy interval of 13.3 [2-23] months. We found no significant correlation between plasma aSyn strain levels and aSyn strain levels measured from brain lysates extracted in RIPA buffer (Fig. 2H) or 2% sodium dodecyl sulfate (SDS) buffer (Fig. S5A). Total aSyn levels also did not correlate in plasma and brain fractions (Fig. S5B). Neither strain nor total aSyn plasma levels correlated with quantifications obtained from brain IHC staining (Fig. S6). These data indicate that plasma levels of aSyn detected by each strain ELISA do not reflect brain aSyn levels in individuals with LBD.

### Strain-selective aSyn antibodies distinguish plasma from individuals with PD versus DLB

Given the mounting evidence for the presence of pathological forms of aSyn outside the central nervous system (CNS), we asked whether the aSyn species detected by our strain ELISAs in blood plasma might reflect clinical disease states. We measured Strain A and Strain B aSyn species in the plasma from NC and individuals with AD, PD, and DLB. In 235 samples from UPenn (Table S5), PD plasma contained significantly higher levels of aSyn detected by both strain-selective ELISAs compared to all other groups (Fig. 3A). Relative to DLB, PD plasma contained seven-fold more Strain A species and 2.5-fold more Strain B aSyn species.

**Figure 3.**
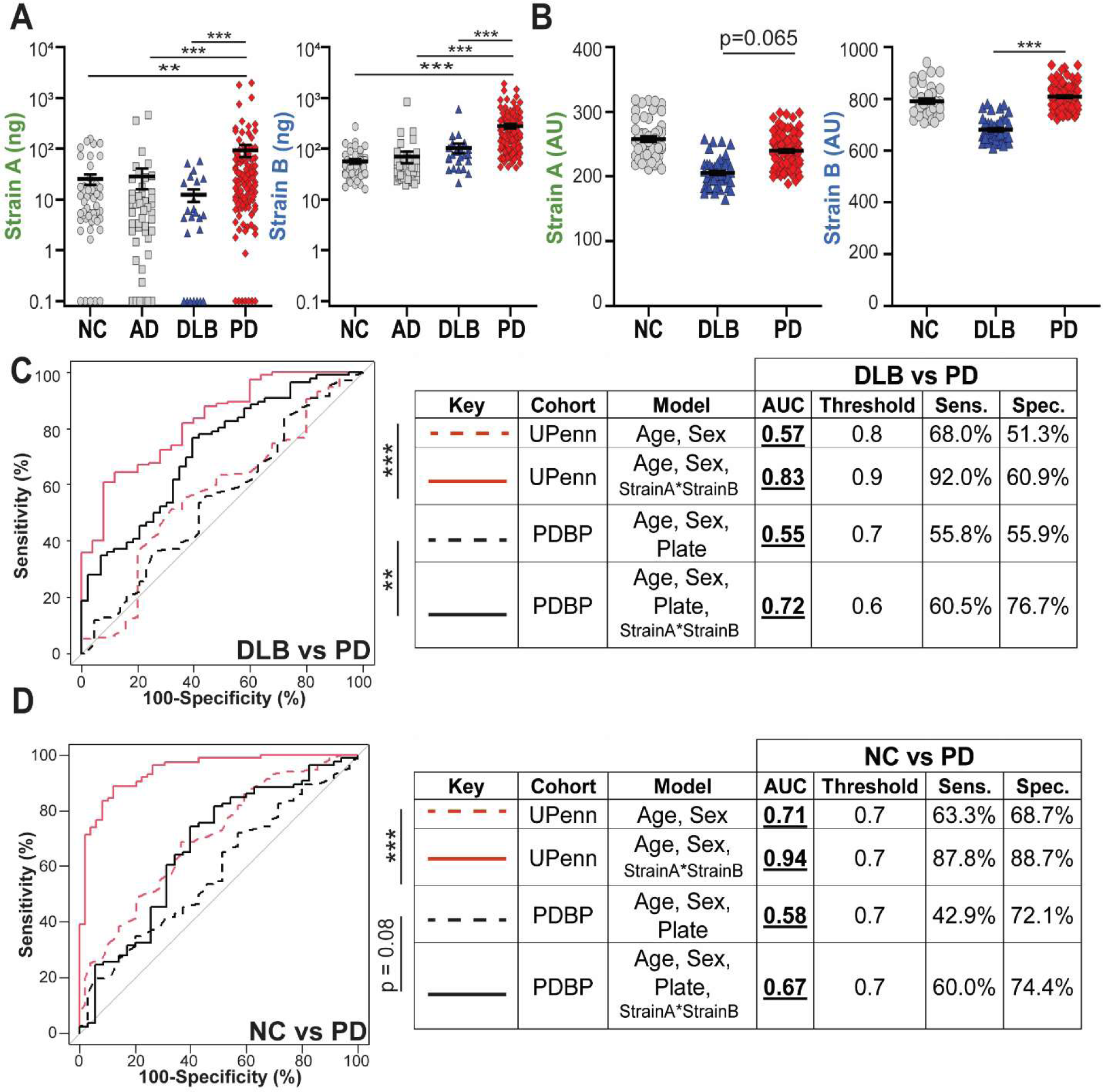
Plasma levels of aSyn strains differentiate individuals with DLB from iwPD. Both aSyn Strain A and Strain B levels are significantly elevated in PD plasma (red diamonds) compared to DLB (blue triangles), NC (grey circles), and AD (grey squares) in the UPenn cohort, n = 235 (**A**), while only Strain B levels are significantly elevated in PD relative to DLB in the PDBP cohort, n = 200. (**B**). Differentiation of PD vs. DLB (**C**) but not PD vs. NC (**D**) in both cohorts is significantly improved (as captured in the Area Under the ROC Curve, AUC) when plasma aSyn strain levels are incorporated into predictors, compared to predictors using only age, sex, and plate to predict disease class. Thresholds, sensitivity, specificity, are established using Youden’s method. Error bars represent SEM. For differences among groups, p-values (corrected for multiple comparisons for Kruskal-Wallis post-hoc tests) for one-way ANOVA and for linear mixed-effects model are shown in **A** and **B**, respectively. Significance testing for differences between ROC curves was performed using the DeLong method. **p < 0.01, ***p < 0.001. AD = Alzheimer’s disease, AU = arbitrary units for plate and batch normalized measures, AUC = area under the curve, DLB = dementia with Lewy bodies, NC = normal control, PD = Parkinson’s disease, ROC = receiver-operator characteristic, Sens. = sensitivity, Spec. = specificity.

We sought to replicate our findings in an additional multi-site cohort, from the NIH-NINDS Parkinson’s Disease Biomarker Program (PDBP). In 200 PDBP samples (Table S5), we again found that PD plasma had higher Strain B aSyn measures than DLB, with a non-significant elevation (p=0.065) in Strain A measures (Fig. 3B). Moreover, in receiver operating characteristic (ROC) curve analyses, aSyn plasma strain values significantly improved discrimination of DLB vs. PD samples over models based only on age and sex in both the UPenn cohort (AUC of 0.83 vs. AUC 0.57, Fig. 3C) and the PDBP cohort (AUC of 0.72 vs. AUC 0.55, Fig. 3C). For discrimination of PD vs. NC samples, aSyn plasma strain values significantly improved discrimination for the UPenn cohort (AUC of 0.94) but not the PDBP cohort (AUC of 0.67, Fig. 3D).

Given the evidence for unique strains in MSA vs LBD, we measured aSyn strain levels in plasma from 18 individuals with MSA (iwMSA)(*26, 32*). While the limited sample size (reflecting the rare occurrence of MSA) did not allow us to detect significant differences in iwMSA compared to other groups, Strain A measures from iwMSA were closer to iwPD than iwDLB (Fig. S7A-B, Table S6).

To determine whether these findings were unique to aSyn species detected by Strain A and Strain B ELISAs, we also evaluated total aSyn levels in plasma by ELISA in UPenn (n = 178, Table S7) and PDBP (n = 200, Table S5) samples (Fig. S7C-D). While addition of plasma total aSyn levels did improve discrimination of DLB vs. PD slightly, improvement in AUC was much more modest over models based on clinical variables alone (Fig. S7E-F) than what we observed using Strain A and Strain B ELISA measures. Taken together, our data suggest that aSyn species, detected by our strain ELISAs in plasma but not in CSF, differentiate DLB from other aSyn pathology states.

### Elevated plasma Strain A levels predict slower subsequent cognitive decline in PD

DLB and PDD differ clinically in the timing of cognitive decline, with the former showing early cognitive impairment and the latter showing cognitive decline after at least one year of motor symptoms (*8, 12, 16, 18*). aSyn strains may have differential ability to spread from cell to cell, and the spread of aSyn pathology from brainstem to cortical areas is a major factor in the development of cognitive decline in the LBD (*18, 22*). Given the finding of lower plasma aSyn strain levels in iwDLB, who have a higher rate of clinical cognitive decline compared to iwPD, we asked whether plasma aSyn strain measures might predict cognitive course in PD.

In longitudinally-followed iwPD (n=95), we asked whether baseline plasma strain aSyn levels associated with subsequent rates of cognitive decline (Table S8). Higher plasma Strain A levels predicted less rapid cognitive decline in linear mixed-effects models adjusting for age, sex, disease duration, baseline cognitive status, and follow-up time. Specifically, individuals with plasma Strain A measures within the top tertile at baseline were predicted, based on model fit to cognitive scores collected over ten years, to preserve their cognition, whereas those in the middle and lowest tertile were predicted to fall into the cognitively impaired range (Fig. 4A, β=0.013, p=0.004, for effect on change in cognitive test score per year). In contrast, plasma Strain B levels did not associate with subsequent rates of cognitive decline. Neither plasma strain predicted rate of motor decline in linear mixed-effects models adjusting for age, sex, disease duration, baseline motor function, dopaminergic medication dose and follow-up time (Fig. 4B) (*39*).

**Figure 4.**
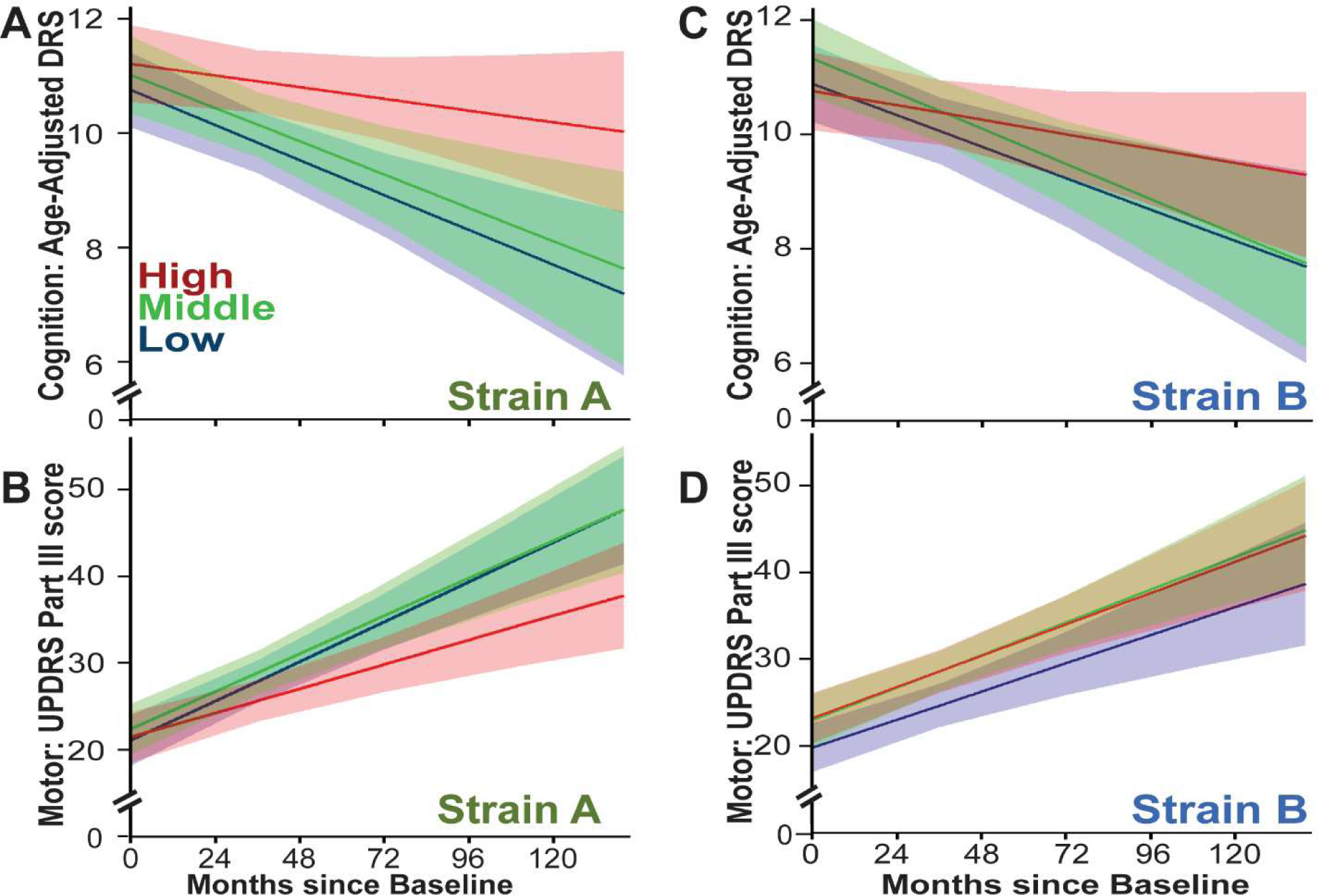
Strain A levels predict the rate of cognitive decline in Parkinson’s disease (PD). Plasma measures of Strain A and Strain B aSyn species were used to predict subsequent cognitive and motor course over 10 years for 95 longitudinally-followed iwPD in linear mixed-effects models. Disease trajectory for iwPD with different levels of Strain A levels at baseline are shown in Panels **A** (cognitive disease course) and **B** (motor disease course). Disease trajectory for iwPD with different levels of Strain B levels at baseline are shown in Panels **C** (cognitive disease course) and **D** (motor disease course). For all panels, colors indicate the modeled disease course for each tertile of aSyn strain measure, from highest (red), to middle (green), to lowest (blue), with bands representing 95% confidence intervals. Cognitive performance is reflected in the age-adjusted DRS score, where higher values represent better performance, and motor impairment is reflected in the Unified Parkinson’s Disease Rating Scale (UDPRS) Part III score, where higher values represent greater impairment.

Because baseline Strain A levels in plasma significantly correlated with subsequent rates of cognitive decline, we asked whether aSyn strain levels changed over time and whether these changes mirrored changes in cognitive status. Using a cohort of iwPD (n = 22) who had at least four serial plasma collections (Table S9), we compared changes in aSyn strain measures over time for those iwPD who had stable cognition versus those iwPD who experienced cognitive decline. We found no significant change in plasma strain values over time, with no significant differences between individuals who did (n=11) vs. those who did not (n=11) develop cognitive decline (Fig. S8).

### Plasma aSyn species induce aSyn fibrillization in seed amplification assay (SAA)

Having discovered plasma aSyn species that discriminate PD vs. DLB individuals and predict subsequent cognitive course in PD, we sought to determine whether these plasma aSyn species had pathogenic potential. To do this, we immunoprecipitated (IP) aSyn species from the plasma and from brain lysates, using Strain A and Strain B antibodies.

We first confirmed that material IP’d from each compartment by our Strain A and Strain B antibodies was enriched for aSyn. To do this, we assayed the IP’d material with (1) our Strain A and Strain B ELISAs, (2) immunoblots detected by multiple anti-aSyn antibodies, and (3) by mass spectrometry (MS) peptide identification. As shown in Table S12, material IP’d with our Strain A and Strain B antibodies from the plasma was enriched by 2.2-to 227-fold, while material IP’d with our Strain A and Strain B antibodies from the brain was enriched by 3.6-to 9803-fold, when quantified with our Strain A and Strain B ELISAs. Immunoblotting our Strain B IP eluates from the plasma with two different anti-aSyn antibodies (HuA and MJFR14) revealed enrichment of a 17kD species, consistent with monomeric aSyn, while immunoblotting our Strain A IP eluates from the plasma with these same anti-aSyn antibodies revealed enrichment of a higher-molecular weight species between 150kD and 250kD (Fig. S9A-D). Finally, MS confirmed aSyn in Strain B antibody IPs from the plasma and the brain, and in Strain A antibody IPs from the brain as well (Figure S9E-F).

Having enriched for Strain A and Strain B aSyn species from each compartment, we asked whether these species could seed fibrillization of monomeric aSyn substrates via seed amplification assays (SAA) based on the real-time quaking induced conversion (RT-QuIC) method (Fig. 5A).

**Figure 5.**
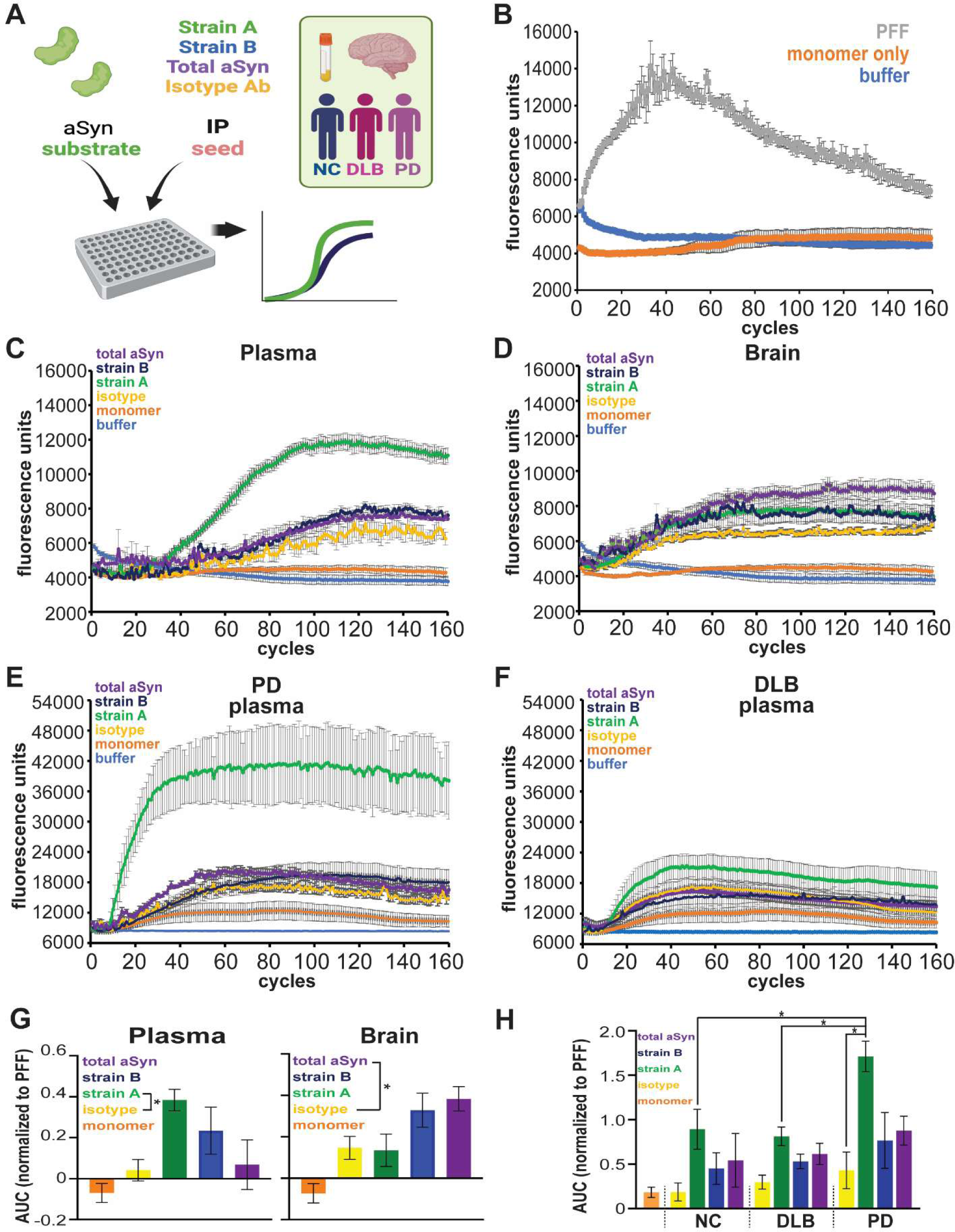
aSyn species isolated from plasma by strain-selective antibodies can seed aSyn fibrillization by seed amplification assay (SAA). **A.** 0.05 ug/mL of protein isolated by immunoprecipitation (IP) using Strain A, Strain B, or total aSyn (syn211) antibodies, or negative control isotype antibody, from plasma or brain lysates (caudate) was tested for ability to seed fibrillization of aSyn monomer substrate in SAA. **B.** In control conditions, addition of aSyn pre-formed fibrils (PFFs) readily induced aSyn fibrillization, while addition of monomeric aSyn did not. **C and D.** Representative SAA curves from plasma (**C**) and brain (**D**) IPs in material from PD individuals. **E and F.** Representative SAA curves from plasma IPs in material from PD (**E**), compared to DLB (**F**) individuals. **G.** Quantification of area under the curve (AUC), normalized to PFF-seeded condition, for n=4 replicate SAAs seeded with 2 different plasma and 3 different brain lysate pools from PD individuals. For plasma, Strain A aSyn (green) was significantly more potent in inducing aSyn fibrillization, compared to the isotype control IP condition (yellow). For brain, total aSyn IP (purple) was significantly more potent in inducing aSyn fibrillization, compared to the isotype control IP condition (yellow). One-way ANOVA (factor: IP seed) with post-hoc Tukey test was used to assess whether AUC was significantly different from isotype versus aSyn seed IPs. **H.** Quantification of AUC, normalized to PFF-seeded condition, for n=3 replicate SAAs seeded with 3 PD, 3 NC, and 3 DLB plasma pools. Plasma aSyn IP’d with Strain A antibody (green) from PD was significantly more potent in inducing aSyn fibrillization, compared to plasma aSyn IP’d with Strain A antibody from DLB or NC. Two-way ANOVA (factors: IP seed and disease group) with post-hoc Tukey test was used to assess for significant differences between SAA conditions. *p<0.05, error bars represent SEM.

As expected, addition of aSyn PFFs induced fibrillization of aSyn monomer in SAA, while monomeric aSyn shaken alone did not (Fig. 5B). Strikingly, aSyn species IP’d from plasma with Strain A and Strain B antibodies induced significant fibrillization in SAA, with a greater effect for Strain A species (Fig. 5C and 5G). In contrast, aSyn species IP’d from plasma with total aSyn antibody did not induce fibrillization over IgG-IP’d controls, and aSyn species IP’d with Strain A antibodies from brain also did not induce fibrillization (Fig. 5C,5D, and 5G; Fig. S10).

We compared SAA behavior for plasma aSyn strain species isolated from iwPD, iwDLB, and NC. Strain A species from iwPD induced more aSyn fibrillization than other groups (Fig. 5E, 5F, and 5H; Fig. S11). These results demonstrate that aSyn species isolated by Strain A and B antibodies from plasma have the capacity to seed aSyn aggregation, while in brain, only species isolated by Strain B and total aSyn antibodies have this capacity. Moreover, Strain A species in plasma and specifically those from individuals with PD relative to DLB have the greatest capacity to seed aSyn amplification.

## DISCUSSION

Using antibodies generated against forms of aSyn with distinct biological properties in cultured neuronal systems, we asked whether distinct aSyn “strains” can be detected in human brain, CSF, and plasma. Remarkably, we found that strain-selective antibodies detect aSyn species in both the plasma and the brain, but not in the CSF. Moreover, plasma aSyn species discriminate PD from DLB samples, and Strain A aSyn measures in plasma at baseline predict subsequent cognitive decline in iwPD. Plasma aSyn species detected by Strain A vs. Strain B antibodies differ from one another and across LBD states in multiple, complementary assessments of their conformation and seeding potential. Together, our data provides human evidence for a strain model of LBD, where distinct conformations of aSyn confer different biological properties that result in different clinical manifestations. Moreover, they suggest that aSyn species outside the CNS have pathological properties, with implications for models of PD pathogenesis that are currently CNS-centric.

Our finding that aSyn Strain A and Strain B antibodies detect aSyn species in the plasma but not in the CSF is notable for several reasons. First, while aSyn is widely expressed across many cell and tissue types in the body, prior characterization of the Strain A and Strain B antibodies used here (*35, 36*), confirmed in our current study, demonstrates that they are selective for overlapping but distinct pathological forms of aSyn. Specifically, in IHC of postmortem brain samples, these antibodies detect abundant aSyn LBs, dot-like inclusions, and Lewy neurites (shown in Figure 1) in LBD brain, absent in brain samples from NC individuals. Moreover, as shown in Figure 2, Strain A and Strain B antibodies detect HMW species of aSyn in LBD brain lysates, not detected in NC brain lysates. Finally, with respect to recombinant aSyn species, the Strain A antibody preferentially detects fibrillar forms of aSyn used in our ELISAs over monomeric forms (shown in Figure 2A). Thus, the detection of aSyn species with these antibodies in the plasma, as well as data demonstrating that species from plasma, particularly in individuals with PD, can template monomeric aSyn in SAA, suggests that pathological forms of aSyn exist in this peripheral compartment. To our knowledge, there is only one other publication to date that demonstrates the potential of aSyn from PD plasma to induce *in vitro* aggregation (*40*). Moreover, the fact that these aSyn species are not readily detected in the CSF and that plasma and brain aSyn species detected by these antibodies (1) do not correlate within an individual and (2) have different *in vitro* capacities to induce fibrillization of monomeric aSyn suggest a non-CNS source for these plasma aSyn species.

In this context, we note that numerous studies have reported aSyn to be enriched in L1CAM-associated extracellular vesicles (EVs) with varying levels of ability to discriminate PD from control samples (*41–44*). Despite L1CAM’s expression in various other tissues including endothelium, epithelium, and immune cells, and our prior report that L1CAM antibodies may not be specific for transmembrane (and thus EV-anchored) isoforms, these studies have assumed a neuronal origin for these aSyn species (*45–47*). At the same time, blood-derived EV-associated aSyn, including phosphorylated and oligomeric forms, have been shown to have prion-like properties, with ability to induce *in* vitro aSyn aggregation and to accelerate propagation of *in vivo* aSyn pathology in mouse models (*48–51*). Our study raises the question of whether L1CAM-EV-associated aSyn in these prior studies is indeed CNS-derived. Future work in this area, along with mechanistic studies assessing the pathogenicity of plasma-derived aSyn strains, would inform our understanding of LBD pathogenesis.

While it remains to be seen whether plasma aSyn strains are *pathogenic*, our current finding that plasma aSyn detected by our strain A and strain B ELISAs discriminate PD from DLB in ELISA and SAA argues that these plasma species reflect pathology, as opposed to normal aSyn function. Three aspects of our plasma aSyn strain studies warrant emphasis. First, to date, there are no reliable biofluid-based biomarkers that differentiate PD from DLB (*20*). As such, our work is important to researchers seeking neurodegenerative disease biomarkers. Moreover, an ongoing debate in the field concerns whether there are true biological differences between PD and DLB, since neuropathological and genetic evidence shows that PD/PDD and DLB are highly similar (*4, 8, 20*). Our study suggests that PD and DLB do differ biologically, that these two clinical entities may stem from underlying differences in aSyn conformation. Second, our findings are reflected in samples drawn from 12 clinical sites in a discovery-replication design. Thus, the observed differences between PD and DLB in plasma aSyn strain values are robust and unlikely to reflect site-based confounders. Indeed, as observed in many multi-site clinical studies, we find evidence of pre-analytical variability in our PDBP cohort studies, with batch and site differences in measures of aSyn strains and total aSyn (Fig. S12). These sources of noise may have contributed to our failure to replicate differences in aSyn strain measures comparing PD and NC. Alternately, the substantial differences between PD and NC groups in plasma aSyn strain measures seen in UPenn samples could reflect an unknown confounder, despite harmonized collection protocols and uniform banking at one site for PD, NC, AD, and DLB samples from UPenn. In either case, our current findings underline the value of multi-site replication in adding confidence to biomarker observations. Third, within the PD group, plasma Strain A values predict subsequent cognitive course, with individuals with higher plasma Strain A measures preserving cognition over time. The fact that within-group analyses demonstrate that plasma Strain A measures are informative also increases confidence that our findings reflect true biological processes. Indeed, a model in which conformation influences aSyn’s propensity to spread to neocortical areas is consistent with both our finding that aSyn strain measures discriminate PD from DLB and our finding that aSyn strain measures predict cognitive decline.

Limitations of our study should be considered alongside its strengths. For example, the site-to-site differences in total aSyn and aSyn strain measures observed in the PDBP cohort decrease our signal-to-noise ratio. Moreover, because PDBP already uses standardized procedures for sample collection and storage, site-to-site variability suggests that plasma-based aSyn measures will inevitably suffer from some degree of noise, limiting potential as a clinically-useful biomarker. That said, the biological insight gained from the current study is still considerable. Second, as previously mentioned, while our study provides clinical evidence for aSyn strains in the LBD and demonstrates that plasma aSyn species isolated with Strain A and Strain B antibodies can induce aSyn fibrillization by SAA, the *in vivo* pathogenicity of plasma-derived aSyn strains remains to be determined. Third, our study is based on two strain-selective antibodies and thus can only characterize a subset of conformations of aSyn. There may be other aSyn conformations that delineate other features of LBD clinical heterogeneity and have the ability to distinguish LBD from MSA (which the antibodies used in this study do not, see Fig. S7A). Fourth, our study leaves open the question of where plasma aSyn species detected by our Strain A and Strain B antibodies originate. In this context, we note that that plasma aSyn strain levels did not correlate with aSyn strain levels in red or white blood cell lysates from matched samples (Fig. S13A, Table S13), or with hemolysis measures (Fig. S13B, Table S5), making aSyn from cellular contamination of plasma unlikely to explain our findings. Fifth, while we confirmed by IP-MS that our Strain B antibody IP’d aSyn from the plasma, we were unable to detect aSyn in our Strain A plasma IP material by MS. We believe that this reflects a known limitation of plasma IP-MS, which suffers from poor sensitivity due to the presence of high-abundance plasma proteins such as immunoglobulins and complement components that can “overwhelm” the MS signal. Because the Strain A antibody preferentially detects a non-monomeric species of aSyn, we could not use the same approach we took to IP-MS of Strain B IP’d proteins, where we isolated the 15kD-to 20kD portion of the gel for MS-peptide identification. While this limitation leaves open the possibility that our Strain A antibody may detect something other than an aSyn species in plasma, multiple lines of evidence suggest otherwise. Specifically, by ELISA and immunoblot with multiple aSyn antibodies, we find evidence for enrichment of aSyn species in Strain A antibody IPs from plasma. More important, plasma species IP’d with Strain A antibody consistently induce aSyn aggregation in SAA, with PD plasma Strain A species proving most potent. It is still possible that our Strain A antibody preferentially detects aSyn species in complex with an as-yet-unidentified cofactor (and indeed previously reported structural studies predict the presence of aSyn cofactors (*27*)) that accelerates fibrillization, but this does not detract from the value of the work shown.

Limitations notwithstanding, our study provides evidence for differences in aSyn conformation between PD and DLB, paving the way for aSyn-strain-related biomarker development. Moreover, our work raises the provocative possibility that these aSyn strains may originate from non-CNS sources, opening new avenues for research into LBD pathogenesis. Indeed, should future work support a non-CNS origin for pathogenic aSyn strains, ramifications for therapeutic development would be substantial.

## MATERIALS AND METHODS

### Generation of strain-selective antibodies

Strain-selective antibodies were generated as previously described (*35, 36*). Briefly, aSyn strains were generated through serial passage of 10% aSyn fibrils with 90% monomer and then immunized into mice to generate hybridomas. Monoclonal antibodies were isolated from hybridoma supernatant and validated for specificity with direct ELISA. For this study, monoclonal antibodies 7015 (recognizing Strain A, the original aSyn strain) and 9027 (recognizing Strain B, a new strain of aSyn that co-recruits tau protein into pathological aggregates) aSyn were used. These antibodies have been previously characterized (*35–37, 52*).

### Brain Collection and Characterization

Brain tissue was derived from the University of Pennsylvania (UPenn) autopsy program as previously described (*53*). Briefly, patients are recruited into the autopsy program via informed consent. Upon death, brains are removed by pathologists or pathology assistants. One hemisphere and brain stem are fixed for 2 weeks in 10% neutral buffered formalin, while the other hemisphere is sliced coronally and frozen. Demographic, clinical, diagnostic, and pathologic data are recorded for each individual as determined by monitoring clinician and post-mortem examination (macroscopic, microscopic, and immunohistochemical) by expert pathologists using established diagnostic criteria (*16, 54–59*).

### Brain immunohistochemistry

Autopsy-derived brain tissue from the Center for Neurodegenerative Disease Research (CNDR) at UPenn was fixed overnight in 10% ethanol with 150 mm NaCl, frozen, and then embedded in paraffin for the preparation of 6µm sections for evaluation as previously reported (*53, 60*). Regions analyzed in this study included superior middle temporal cortex (SMTC), anterior cingulate (CING), cerebellum, and striatum. Semi-adjacent sections were immunohistochemically stained for phosphorylated-tau (AT8, ThermoFisher, 0.2 µg/mL, no antigen retrieval), amyloid beta (Aβ; NAB228, CNDR 1:40,000, no antigen retrieval), phosphorylated alpha-synuclein (MJF-R13, Abcam, 0.17 ug/mL, proteinase K antigen retrieval, 4.229 mg/mL), syn9027 (0.026 ug/mL, proteinase K antigen retrieval), and syn7015 (0.026 ug/mL, proteinase K antigen retrieval). Immunostained sections were imaged at 20x magnification on a digital slide scanner. Images were analyzed in QuPath software (version 0.2.0-m5), using pixel classifier tool with empirically derived positive pixel classifiers for each immunostaining to calculate the percentage of DAB-positive pixels as area occupied (%AO) in representative grey matter segmentations as previously validated (*60*).

### Plasma, Blood Cell, and Cerebrospinal fluid collection

Plasma, blood cells, and cerebrospinal fluid (CSF) samples were obtained from patients enrolled in IRB-approved research studies from the UPenn Parkinson’s Disease and Movement Disorder Center (PDMDC), Frontotemporal Dementia Center (FTDC), or Alzheimer’s Disease Center (ADC) as previously described (*53, 61*). Blood samples were drawn into EDTA vacutainers. CSF was collected using plastic round-bottom tubes in the respective clinic. All samples were kept cool using ice packs and insulated bags following collection, and delivered to the CNDR Biobank on the same day.

For plasma collection, EDTA tubes were spun at 2000g for 15 minutes at 4°C. Plasma was aspirated and stored in 500uL aliquots. CSF samples were aliquoted directly into 500uL cryogenic tubes. All samples were stored at -80°C at the UPenn CNDR Biobank until analysis.

For matched blood cell and plasma collection, blood was layered onto an equivalent volume of Polymorphoprep (CosmoBioUSA) and spun at 500 x g for 30 minutes without brake. Plasma and red blood cell fractions were collected and frozen at -80C. White blood cell fraction was incubated in red blood cell lysis buffer (Sigma) per manufacturer protocol and then frozen at - 80C.

### Hemoglobin Absorbance Measurement

Hemolysis was measured in plasma via absorbance to estimate the hemoglobin levels. Absorbance was measured at multiple wavelengths (380, 415, 450) and hemoglobin calculated in g/L based on this formula: (167.2 * A415 – 83.6 * A380 – 83.6 * A450) * 0.001 (*62*). Plasma samples with hemoglobin levels above 30 mg/dL were excluded from analyses.

### Subject Recruitment and Characterization

#### U19 Cohort

Individuals were recruited at UPenn, and plasma and cerebrospinal fluid collected after informed consent. These procedures were approved by the UPenn IRB. Individuals with known genetic AD- or PD-causing mutations were excluded from this and all other cohorts in this manuscript. Individual diagnoses were determined by clinically established diagnostic criteria by the treating physician (*9, 16, 57, 58*). Cognitive and motor testing were performed as previously described (*53*).

To obtain samples for brain immunohistochemistry and matched plasma and brain samples (Table S1), records from the CNDR Integrated Neurodegenerative Database (INDD) were cross-referenced to identify patients with a plasma sample collected within 24 months of autopsy. Brain tissue was, then, surveyed to identify patients with available tissue from the superior middle temporal cortex, anterior cingulate, and striatum. A subset of these potential samples were used to compare detection of aSyn strains by western blot vs. strain and total aSyn ELISA (Table S2). For studies comparing matched CSF and plasma samples, 44 individuals with PD or DLB from the CNDR database were identified who had less than a two-year interval between collection of these biofluids (Table S3).

To obtain samples from individuals with longitudinally-collected plasma, the CNDR database was searched for PD patients with multiple, serial plasma collection dates. Patients with a clinical diagnosis of PD, four or more plasma sample timepoints spanning 4 or more years, and a normal cognitive consensus diagnosis at baseline visit were nominated for analysis of aSyn strain measures over time. 22 iwPD with longitudinal plasma sampling were ultimately selected (Table S7), and their samples were pulled from the CNDR Biobank: 11 individuals experienced cognitive decline over time, and 11 did not. Plasma samples were analyzed using ELISAs for Strain A, Strain B, and total aSyn.

#### PDBP Cohort

The NIH Parkinson’s Disease Biomarker Project (PDBP) maintains a number of recruitment sites across the continental United States whose practices are previously published (*63*). Eligible patients are recruited and enrolled at these locations, and resulting samples are delivered to a central location for processing and deposition into the biobank. Age and sex-matched plasma samples from NC, iwPD, and individuals with DLB were requested and approved by the NIH PDBP steering committee (Table S4).

### Strain-Specific aSyn ELISA

#### Standards

Monomeric aSyn was generated through recombinant expression in E. coli followed by dialysis-based purification as previously described (*64*). Recombinant wild-type human aSyn PFFs were generated as previously described through shaking of monomeric aSyn at 37°C for 7 days (*35, 64*). aSyn monomer and PFFs were used to generate standard curves for each ELISA plate. PFFs were sonicated on high power (10 cycles of 30 sec on and 30 sec off) at 10°C with the Diagenode Bioruptor Plus (Diagenode B01020001, Denville, NJ, USA) prior to use.

#### ELISA

Flat, clear-bottom 384 well plastic plates (NUNC Maxisorp) were coated overnight at 4°C with MJFR-1 (Abcam ab138501) at 0.4 or 0.2 ng/uL as the capture antibody for the strain A or B ELISAs, respectively, then thrice washed in PBS with 0.05% Tween-20 and blocked with Block ACE buffer overnight at 4°C. Samples were applied to ELISA diluted in PBS with 0.05% Tween-20 and incubated for four hours at 37°C. Plasma was diluted 1:1 for 9027 (Strain B) and 2:1 for 7015 (Strain A) ELISAs. Brain lysate and CSF were diluted at 1:20-1:50 and 1:1, respectively. After washing in PBS with 0.05% Tween-20, biotinylated 9027 (Strain B) or 7015 (Strain A) antibodies were added overnight at 4°C at 0.166 ng/uL and 1.33 ng/uL, respectively. Plates were then incubated, after washing with PBS with 0.05% Tween-20, in Streptavidin-HRP at 1 ug/mL for 1 hour at room temperature while being protected from light. A final wash was completed before ELISA plates were developed with TMB substrate solution for 10 minutes. The reaction was stopped with 10% phosphoric acid and read immediately on a FluoStar Omega (BMG LabTech, Cary, NC, USA) plate reader at an absorbance of 450 nm. Plates were centrifuged at 1000 x g for 1 minute at room temperature before each incubation step except for blocking and application of TMB or phosphoric acid. Sample concentrations were determined using a five-parameter logistic curve based on aSyn PFF standard curves run on the same plate.

### Total aSyn ELISA

Total aSyn was measured using the chemiluminescence-based (Cat. No. 844101) or colorimetric-based (Cat. No. 448607) LEGEND MAX ™ Human α-Synuclein ELISA Kit (following the commercial protocol. Dilutions for plasma and brain lysate were 1:200 and 1:10,000-1:50,000, respectively.

### Brain Tissue Lysis and Sequential Extraction

To prepare brain lysates, 50mg of Diagenode Protein Extraction Beads (Diagenode C20000021, Denville, NJ, USA) were loaded into 0.5 mL Axygen Maxymum Recovery tubes (Axygen #PCR-05-L-C, Tewksbury, MA, USA) and placed onto dry ice. Brain tissue was sectioned into 35 – 55 mg fragments and added to the extraction beads with 5 uL of 1x RIPA buffer (50 mM Tris Base, 150 mM NaCl, 1% NP-40, 5mM EDTA, 0.5% sodium deoxycholate, and 0.1% SDS) per mg tissue and placed on ice. Protease inhibitor cocktail (25mg of Pepstatin, 25mg of Leupeptin, 25mg of TPCK, 25mg of TLCK, 25mg of Trypsin Inhibitor in 5ml of 0.5M EDTA and 3ml of 100% EtOH), and PMSF (Thermofisher, Waltham, MA, USA) were added to both the 1x RIPA and the 2% SDS solutions (1:1000 and 1:500 dilutions, respectively).

Tubes were gently vortexed and placed in the Diagenode Bioruptor Plus (Diagenode B01020001, Denville, NJ, USA) and sonicated on the high-power setting at 4°C for 5 cycles of 30 seconds of sonication, followed by 30 seconds of rest. Samples were subsequently vortexed and put through two additional rounds of sonication. Tubes containing tissue that was incompletely lysed were sonicated for up to 2 additional rounds. The tubes were then spun down at 15,000xg for 30 minutes at 4°C. The supernatant was extracted and stored at -80°C. Resulting pellets were placed on ice, resuspended in 2% SDS, and subjected to the same sonication protocol as above.

### Gel Electrophoresis and Western Blotting

Samples for 2D gel electrophoresis were diluted in 5x sample buffer. Brain lysates but not IP eluates were boiled for 5 minutes at 95°C whereas other samples were not. Samples were loaded and run on 4-20% Criterion TGX Midi Protean gels (Bio-Rad #5671094, Hercules, CA, USA) and transferred to nitrocellulose 0.45 µm membrane (Bio-Rad #1620113, Hercules, CA, USA) via semi-dry TransBlot Turbo (Bio-Rad 1703848, Hercules, CA, USA) transfer apparatus. Blots were blocked in 5% milk with Tris-buffered saline (TBS) with 0.1% Tween-20 (TBS-T) for 1 hour. Blots were incubated in primary antibody (7015 1.29 µg/mL, 9027 2.57 µg/mL, syn211 1:200, GAPDH 0.33 ug/mL) overnight in 5% milk in TBS-T, washed with TBS-T twice and once in TBS, then incubated in secondary antibody (goat anti-mouse 2.67 µg/mL). Blots were washed again and then developed with Westernbright ECL (K-12045) or Sirius (K-12043) reagent (Advansta, San Jose, CA, USA) before imaging on the Chemidoc system (Bio-Rad #12003154, Hercules, CA, USA).

### Epitope Mapping

Epitope mapping was performed as previously described (*36*). Briefly, full-length or fragments of human aSyn were recombinantly expressed in *E. coli* and purified, run on a SDS-PAGE 5-20% gel, and probed by Western blot with Syn9027.

### Immunoprecipitation

Antibody was immobilized on columns using an Aminolink kit (Thermofisher #44890, Waltham, MA, USA) per manufacturer’s protocol. For each column, 1 mg of antibody was covalently immobilized. At least 2 mL of pooled plasma or 6.5 mg of pooled brain lysate, which contain approximately equal amounts of aSyn strains, (Table S8-S9) were applied to each column, as indicated in the text, and allowed to bind for 1.5 hours at room temperature. aSyn was eluted from columns using 3 mL of 1M glycine pH 2.5 buffer per column and immediately neutralized using 1M Tris pH 9.0 to pH 7 before storage at -80°C.

### Seed amplification Assay (SAA)

SAA protocols were adapted from previously published protocols (*65*). Briefly, a 96-well clear bottom black-well plate (Thermofisher #3165305, Waltham, MA, USA) was loaded with 6 silica beads per well (OPS Diagnostics PFMB 800-100, Lebanon, NJ, USA). Reaction mixture was made to a final volume of 100 uL (40 mM phosphate buffer, 170 mM NaC1, 10 uM thioflavin T, 0.1 mg/mL monomeric aSyn, 0.0015% SDS, pH 8.0). Eluates were loaded at a total protein concentration of 0.05 ug/mL and aSyn PFFs to a final concentration of 0.01 mg/mL. All samples were run in quadruplicate at 42°C on a FluoStar Omega, with plates read up to 160 cycles (1 min shaking and 1 min rest) at 300 rpm double orbital shaking. Fluorescence readings were taken every 45 minutes at excitation of 448 nm and emission of 482 nm with gain set to 1375. Fluorescence calculations were performed after subtraction of background based on buffer alone condition and normalized as indicated in the text. Area under the curve was calculated using the trapezoidal method in R. C50 was calculated based on the cycle at which half of maximum normalized fluorescence for each seed was reached.

### Mass spectrometry

After 2D gel electrophoresis, gel was stained with Coomassie blue and indicated range of molecular weight bands were cut out of gel for IP samples. Gel was digested and isolated protein subjected to tryptic digestion before undergoing mass spectrometry. Peptide identification and quantification was performed using Scaffold.

### Statistical Analyses

For correlation testing, Pearson coefficients, and p-values for association were calculated. For demographic data, data are reported as mean ± standard error of the mean (SEM) or frequency (%) as appropriate. To test for differences between groups, non-parametric t-tests/ANOVA and chi-square tests were used for numerical and categorical data, respectively. Otherwise, statistical tests for comparisons between groups are indicated in the text or figure legend. ROC curves were generated, with cutoffs selected using the Youden method, and the DeLong method was used to determine statistical significance for differences between ROC curves. Linear mixed-effects models were used throughout the manuscript to (1) normalize PDBP data to account for batch effects using plasma strain or total aSyn level as the dependent variable, with age, sex, loading group, plate, and diagnosis as fixed effects, sample number with random intercept and random slope for plate, (2) predict cognitive trajectory based on baseline plasma strain level using age-adjusted DRS as the dependent variable, with age, sex, disease duration, baseline age-adjusted DRS, and an interaction term between log10 of strain aSyn levels and time as fixed effects, with a random intercept included, (3) predict motor symptom trajectory based on baseline plasma strain level using Unified Parkinson’s Disease Rating Scale (UPDRS) part III as the dependent variable, with age, sex, disease duration, baseline age-adjusted UPDRS part III, and an interaction term between log10 of strain aSyn levels and time as fixed effects, with a random intercept included, and (4) determine whether plasma aSyn strain values change over time using strain plasma concentration as the dependent variable, with baseline age-adjusted DRS, sex, disease duration, age, and an interaction of cognitive group with time as fixed effects, and sample number as random intercept. Correction for multiple comparisons was performed using the Benjamini, Kreuger, and Yekutieli method (*66*). Statistical analyses performed in R and R-scripts are included as supplementary files. All statistical tests are two-sided unless indicated.

## List of Supplementary Materials

Supplementary figures S1-S13

Supplementary tables S1-S13 S14

Table of reagents

R Markdown Files

## Acknowledgements

Data and biospecimens used in preparation of this manuscript were obtained from the Parkinson’s Disease Biomarkers Program (PDBP) Consortium, supported by the National Institute of Neurological Disorders and Stroke at the National Institutes of Health. Investigators include: Roger Albin, Roy Alcalay, Alberto Ascherio, Thomas Beach, Sarah Berman, Bradley Boeve, F. DuBois Bowman, Shu Chen, **Alice Chen-Plotkin**, William Dauer, Ted Dawson, Paula Desplats, Richard Dewey, Ray Dorsey, Jori Fleisher, Kirk Frey, Douglas Galasko, James Galvin, Dwight German, Steven Gunzler, Lawrence Honig, Xuemei Huang, David Irwin, Kejal Kantarci, Anumantha Kanthasamy, Daniel Kaufer, Qingzhong Kong, James Leverenz, Carol Lippa, Irene Litvan, Oscar Lopez, Jian Ma, Lara Mangravite, Karen Marder, Nandakumar Narayanan, Laurie Orzelius, Vladislav Petyuk, Judith Potashkin, Liana Rosenthal, Rachel Saunders-Pullman, Clemens Scherzer, Michael Schwarzschild, Tanya Simuni, Andrew Singleton, David Standaert, Debby Tsuang, David Vaillancourt, Jerrold Vitek, David Walt, Andrew West, Cyrus Zabetian, and Jing Zhang. The PDBP Investigators have not participated in reviewing the data analysis or content of the manuscript.

We thank the Children’s Hospital of Philadelphia (CHOP) Proteomics Core and its director, Dr. Lynn Spruce, for assistance with mass spectrometry experiments. We thank the many patients and their families who contributed samples for this work.

## Author Contributions

Conceptualization: RZ, GTK, RTS, JFM, RSA, DI, ACP

Data curation: RZ, GTK, RTS, RDR, SA

Formal Analysis: RZ, GTK, RTS, RDR, SA, DI, RSA, SX, ACP

Funding acquisition: ACP, TT, DAW, DW

Investigation: GTK, RZ, RTS, RDR, RSA, SA, KD

Methodology: GTK, RZ, RTS, RDR, SA, KD, KL, DW, TT, EL, VL, DI, RSA, ACP

Project administration: GTK, RTS, RDR, ACP

Resources: DW, DAW, TT, EBL, VL, DI, ACP

Supervision: JFM, RSA, ACP Validation: GTK, RZ, RTS, RDR

Visualization: RZ, GTK, RTS, RDR, SA, ACP

Writing – original draft: GK, RZ, ACP

Writing – review & editing: all authors

## Data and materials availability

R scripts for analysis are included in the supplementary files and all methods or reagents used are included in the text or supplementary files. Reagents produced at UPenn are available upon request pending material transfer agreements. All relevant data are included in the manuscript or in supplementary files, but raw data can be provided upon request.

## SUPPLEMENTARY FIGURES AND TABLES

**Supplementary Figure 1.**
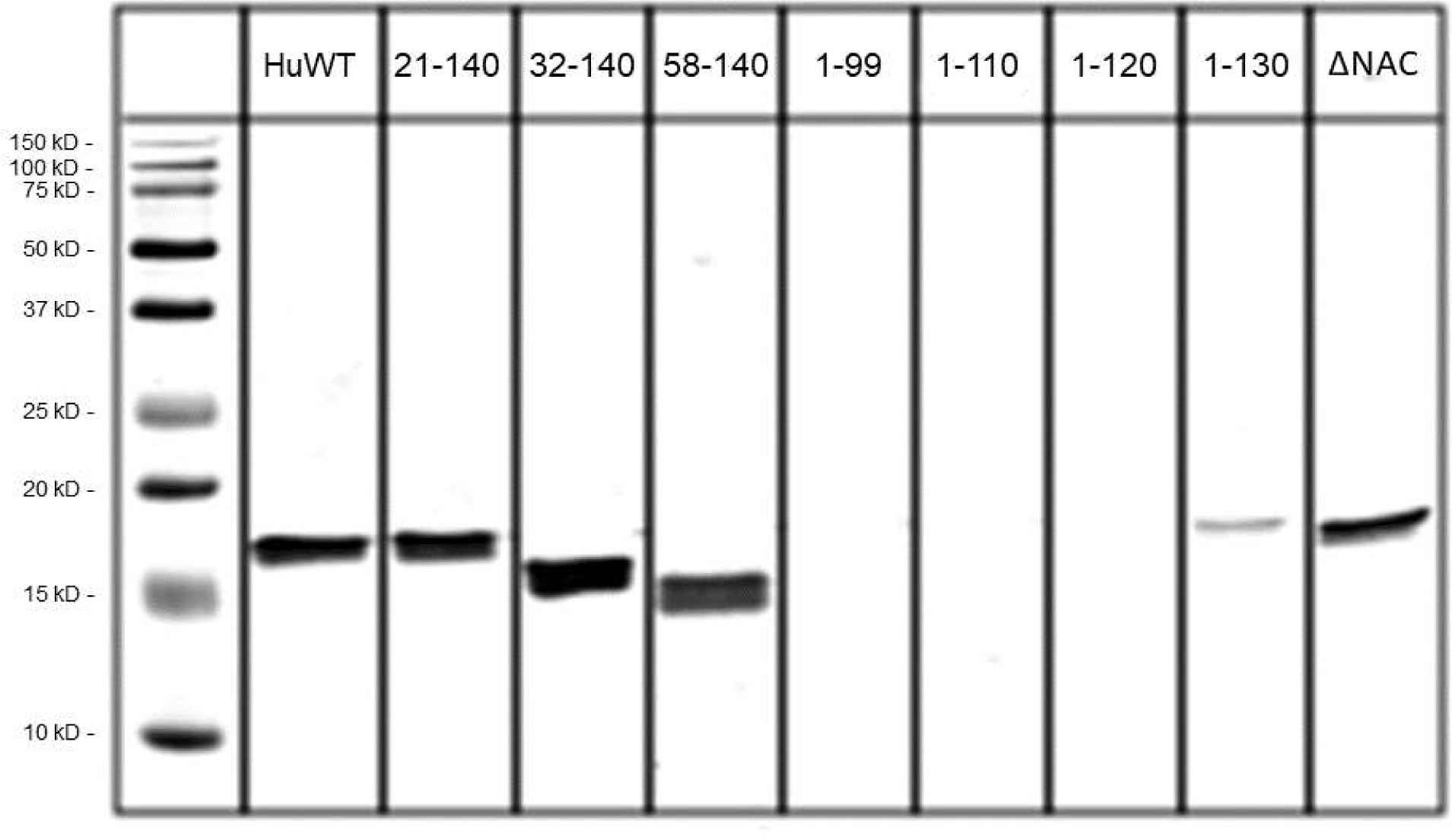
Epitope map for monoclonal 9027 antibody generated against strain B. Immunoblotting against full-length human wildtype (HuWT) aSyn and indicated peptide fragments demonstrate that the 9027 antibody detects a continuous region on aSyn between residues 120-140. ΔNAC = NAC domain deletion mutant.

**Supplementary Figure 2.**
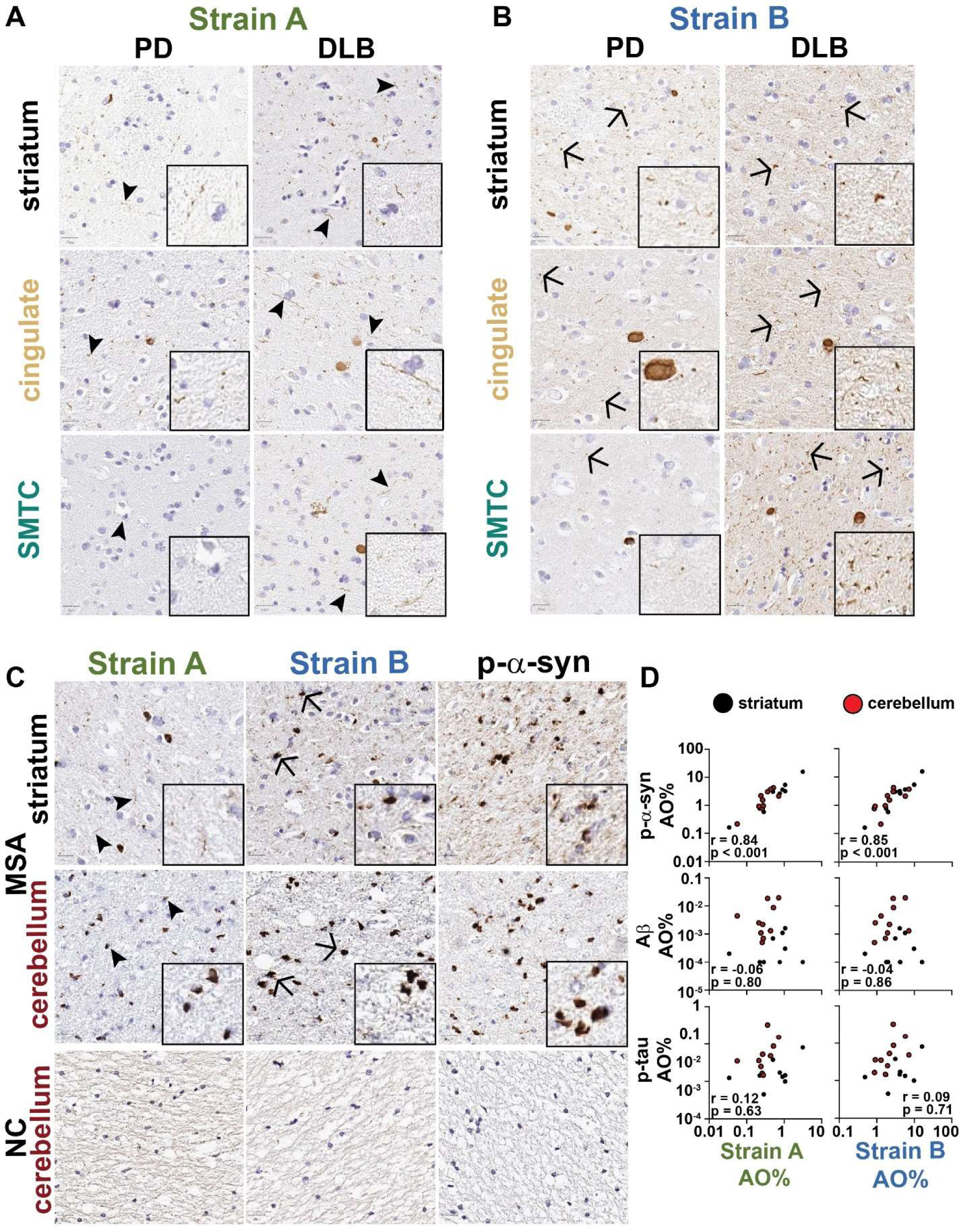
Strain-selective antibodies detect overlapping but distinct. (**A**) and strain B (**B**) immunohistochemical staining are shown for PD (A) and DLB from the set of 10 individuals with LBD in Fig 1. Representative images of strain A, strain B, and p-aSyn immunohistochemical staining are shown from a set of 10 individuals with MSA (**C**). aSyn strain staining correlates strongly with p-aSyn in MSA brains but not with Ab or p-tau staining in both striatum (black) and cerebellum (red, **D**). AO = area occupied, DLB = dementia with Lewy bodies, PD = Parkinson’s disease, PDD = PD dementia, SMTC = superior middle temporal cortex.

**Supplementary Figure 3.**
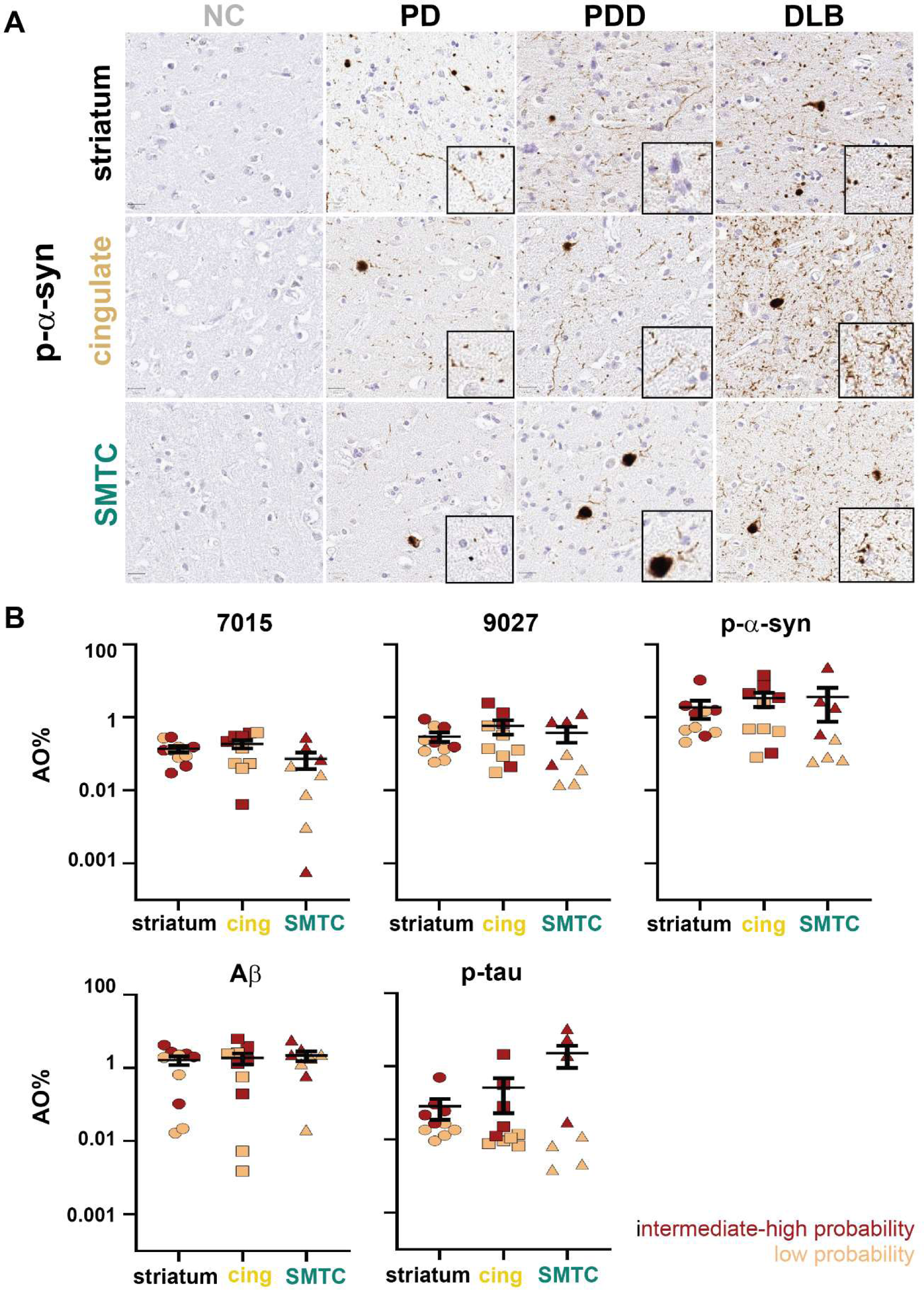
Strain B immunostaining is higher in individuals with higher. immunohistochemical staining are shown (**A**) from a set of three NC and ten individuals with LBD. The levels of strain A, strain B, p-aSyn, Aβ, and p-tau stratified by intermediate/high (red) or low (tan) probability of secondary AD neuropathologic diagnosis across striatum (circles), anterior cingulate cortex (squares), and SMTC (triangles) are shown (**B**), indicating higher AD co-pathology in areas with higher levels of Strain B antibody staining. Line represents mean and error bars represent SEM. AO = area occupied, cing = anterior cingulate cortex, SMTC = superior middle temporal cortex.

**Supplementary Figure 4.**
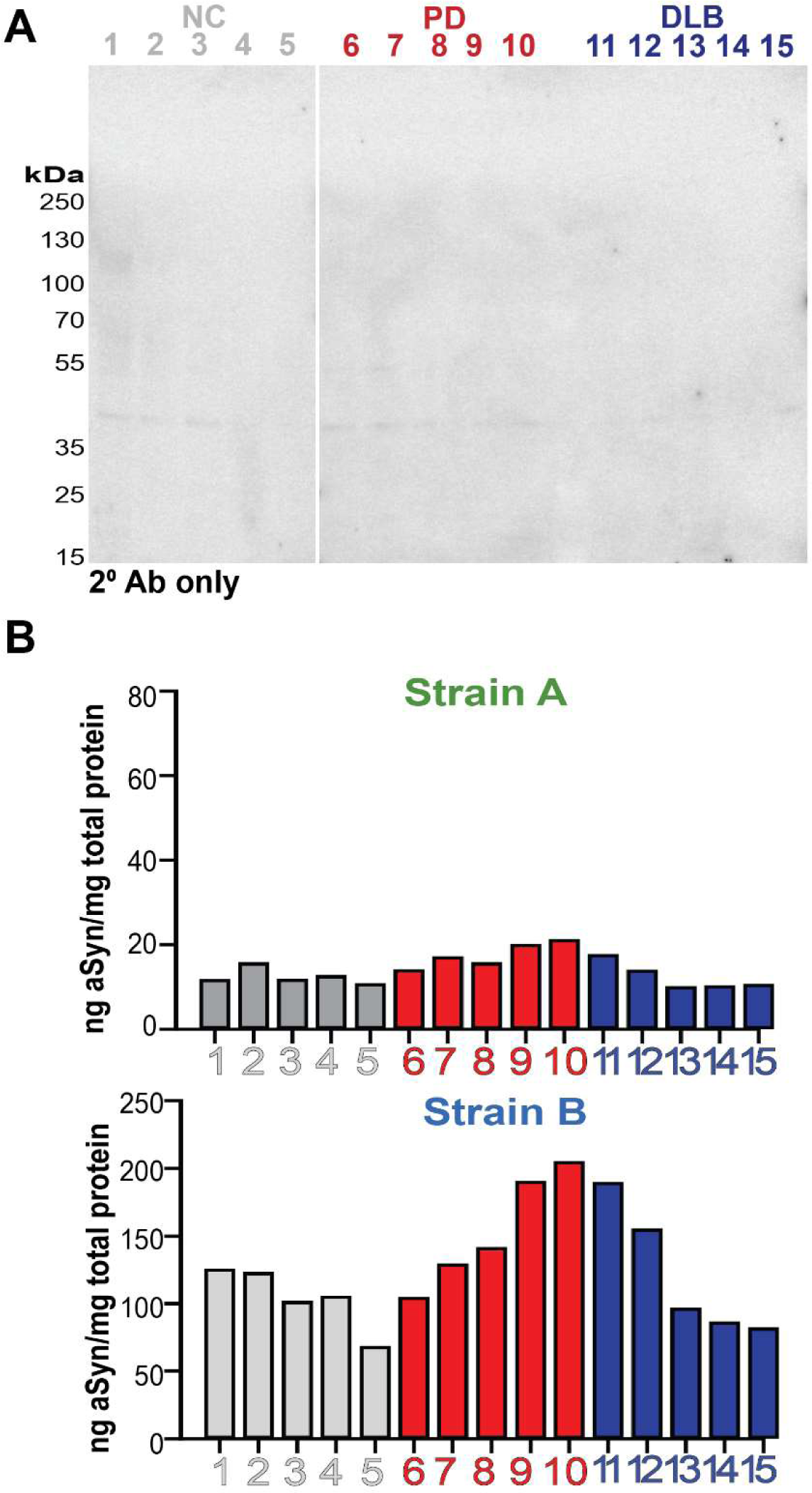
aSyn strain ELISAs detect pathologic forms of aSyn. Western blot of caudate brain lysates from healthy controls (NC), individuals with Parkinson’s disease (PD), and individuals with dementia with Lewy bodies (DLB) with secondary antibody only does not show significant off-target binding (**A**). Levels of aSyn strain levels in cerebellum brain lysates in individuals with PD or DLB were lower than detected in caudate (Fig. 2E-F), a region relatively enriched for aSyn pathology.

**Supplementary Figure 5.**
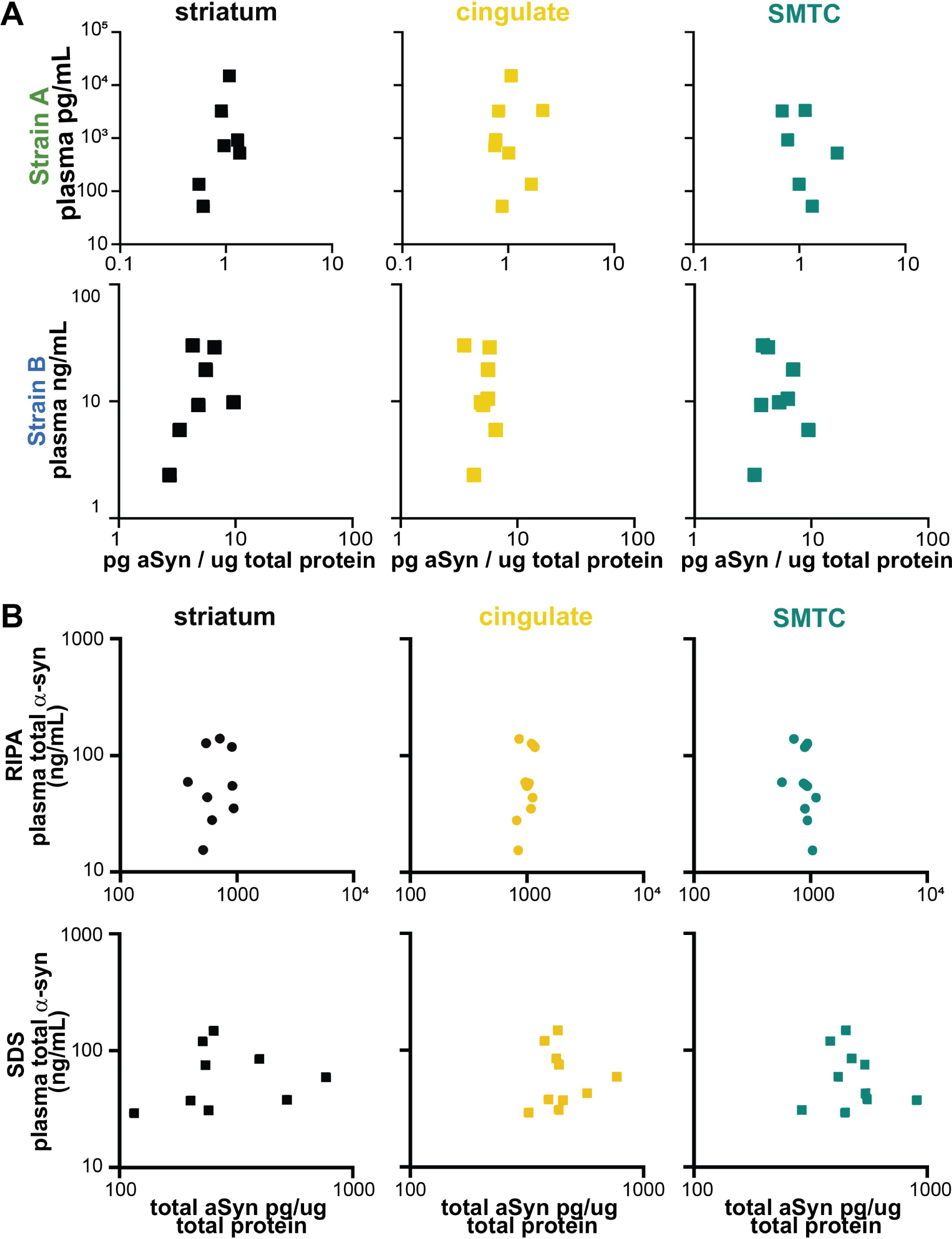
Strain and total aSyn plasma levels do not correlate with brain levels. For individuals with plasma collected within two years of autopsy (n = 10), plasma strain aSyn levels do not correlate with levels in brain caudate lysates extracted in 2% SDS buffer in regions associated with diffuse cortical spread of Lewy pathology (**A**). Total plasma aSyn levels also do not correlate with brain lysate levels in RIPA or SDS fractions (**B**). SDS = sodium dodecyl sulfate, SMTC = superior middle temporal cortex.

**Supplementary Figure 6.**
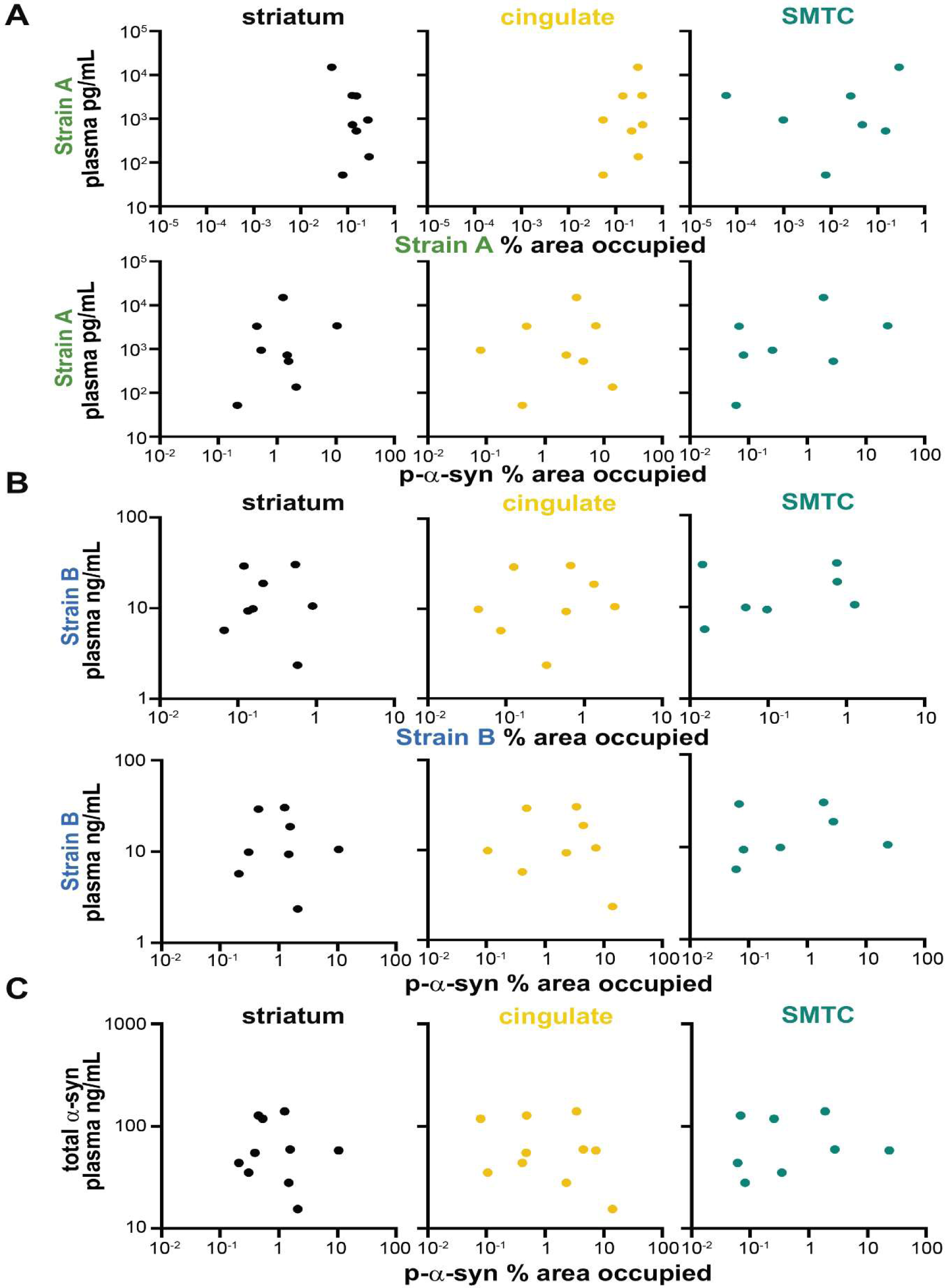
Plasma levels of total and strain aSyn measured by ELISA do not correlate with brain immunohistochemistry. For plasma samples collected within two years of autopsy (n = 10), plasma strain (**A, B**) or total (**C**) aSyn levels do not correlate with levels of immunohistochemical staining in regions associated with diffuse cortical spread of Lewy pathology. SMTC = superior middle temporal cortex.

**Supplementary Figure 7.**
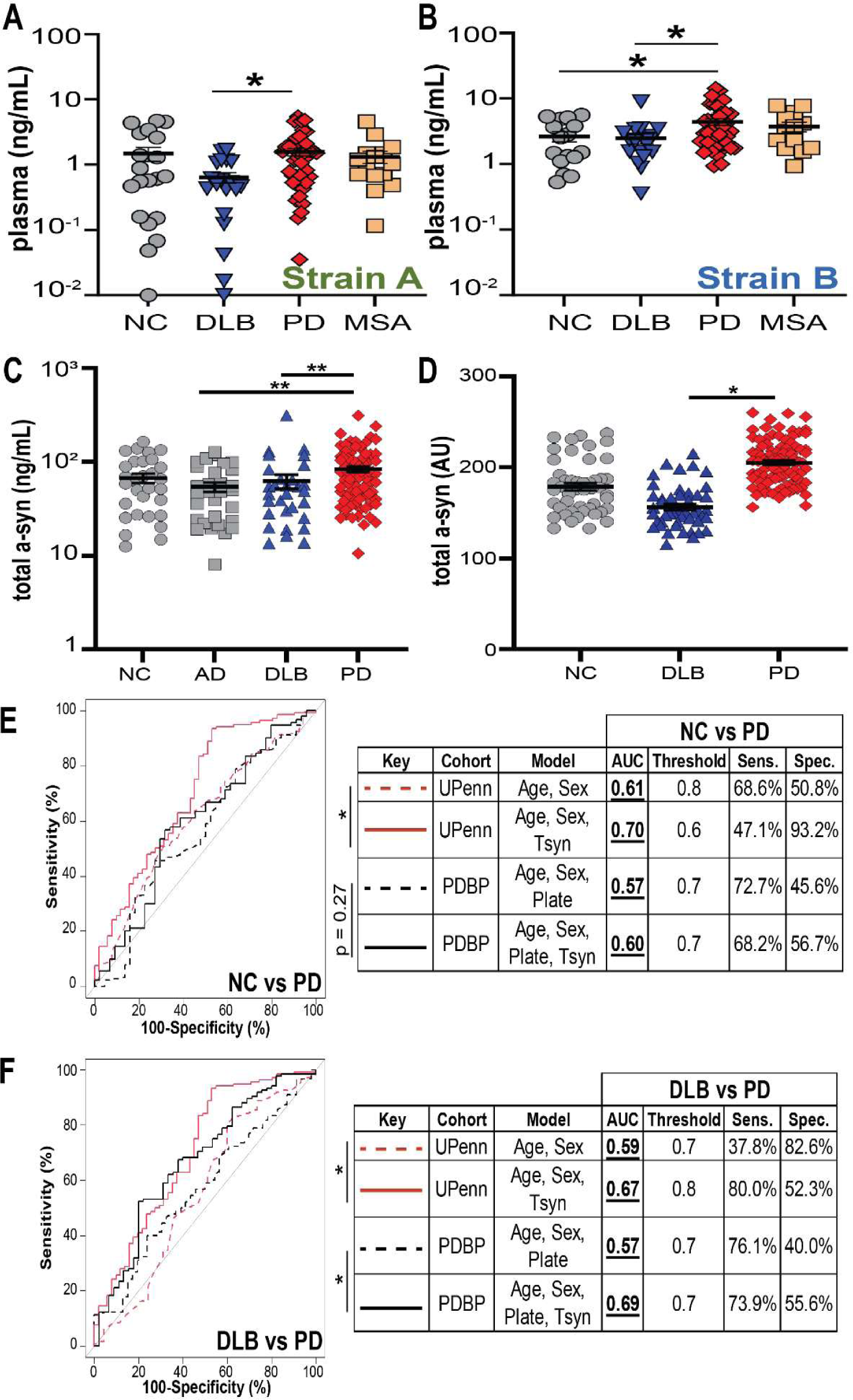
Plasma aSyn strain levels do not differ between individuals with Parkinson’s disease (PD) and multiple systems atrophy (MSA) and total plasma aSyn levels do not robustly differentiate PD and dementia with Lewy bodies (DLB). Measurement of aSyn strain levels (n = 101) by ELISA reproduced the decrease previously observed in plasma from dementia with Lewy bodies relative to PD but levels were similar between PD and MSA plasma (**A, B**). Total plasma aSyn levels were measured by a commercially available ELISA in the UPenn (**C**) and PDBP cohorts (**D**). ROC curves were used to differentiate PD vs. NC (**E**) or PD vs. DLB (**F**) incorporating age, sex, and plate with or without plasma total aSyn strain levels, as predictors. In all cases, incorporation of total aSyn plasma measures minimally improved discrimination. Thresholds generated by the Youden method. Error bars represent SEM. For differences among groups, p-values (corrected for multiple comparisons for Kruskal-Wallis post-hoc test) for one-way ANOVA are reported in **A** and **B.** Significance testing for differences between ROC curves was performed using the DeLong method. AU = arbitrary units, NC (grey circles) = normal controls, DLB (blue triangles) = dementia with Lewy bodies, PD (red diamonds) = Parkinson’s disease.

**Supplementary Figure 8.**
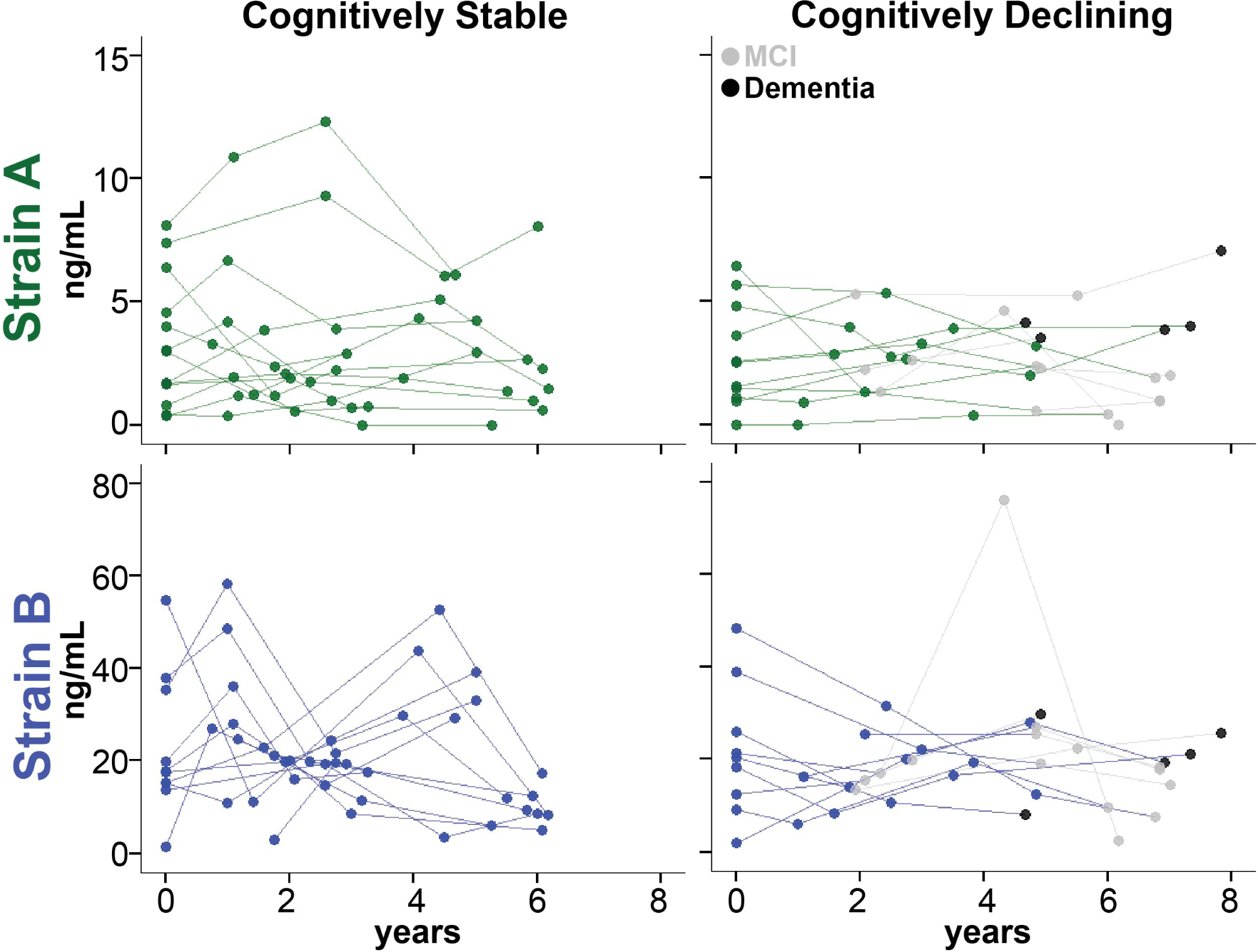
aSyn strain plasma levels do not significantly change over time and do not mirror changes in cognition at the individual level. Strain A and Strain B plasma levels were measured in 22 individuals with longitudinal plasma sampling. No differences were observed between individuals who did (n=11) vs. did not (n=11) develop cognitive decline. Groups were compared using a linear mixed-effects model adjusting for disease duration, baseline age, baseline cognition, and sex.

**Supplementary Figure 9.**
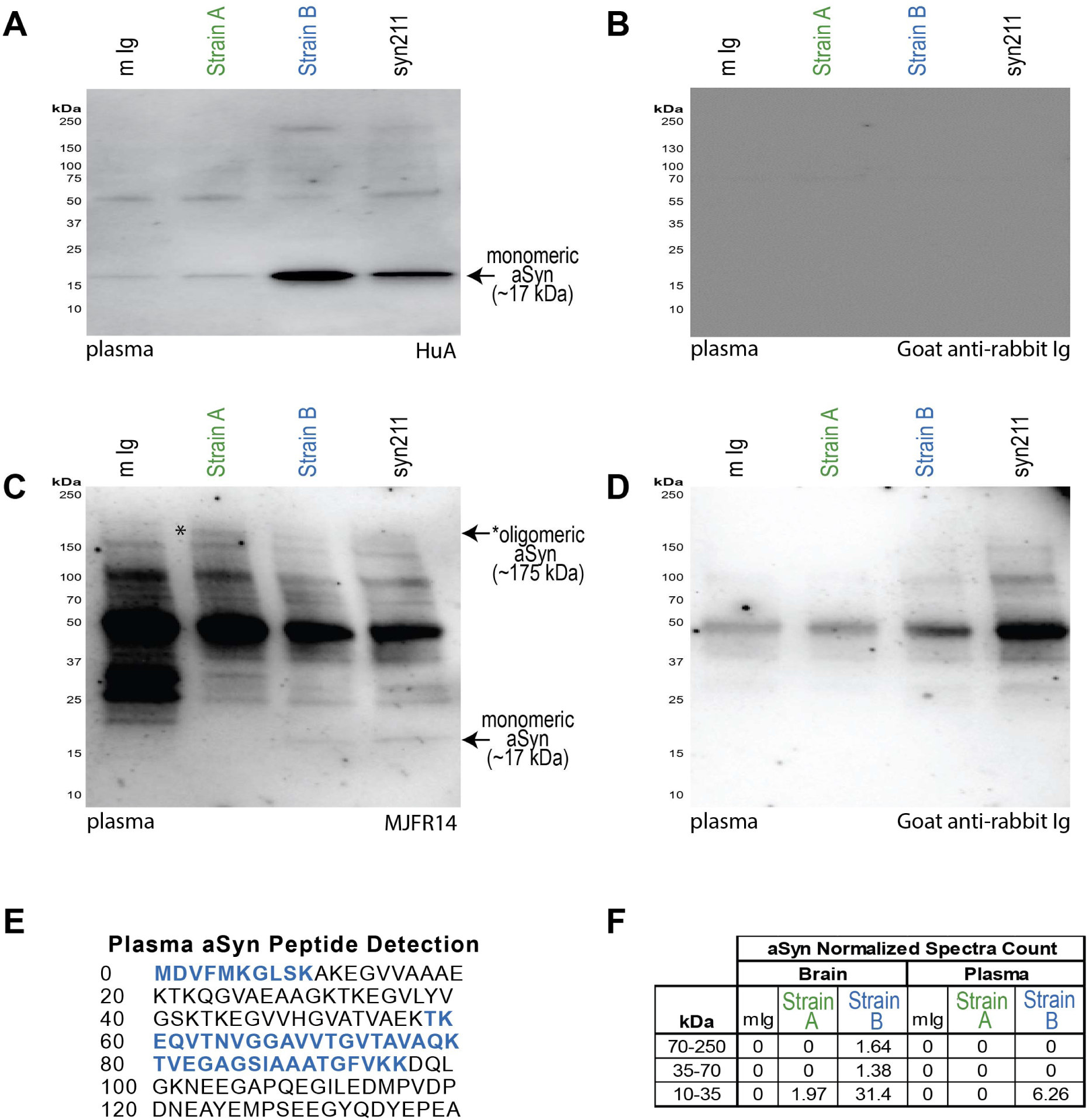
aSyn is detectabIe after immunoprecipitation with strain-selective antibodies. Strain A, Strain B, or total aSyn (syn211) species were isolated by IP from pooled PD plasma or pooled PD brain caudate lysates. Western blot (WB) with HuA, a non-selective anti-human aSyn antibody, demonstrates robust signal at expected molecular weight of monomeric aSyn (∼17 kDa) in Strain B and syn211 plasma IP with weak detection in Strain A plasma IP (A). WB with MJFR14, an anti-human aSyn antibody that prefers oligomeric species, demonstrates a ∼175 kDa band (asterisk) that is enriched in the strain A IP (C) and not seen when probing with secondary antibody alone (B, D). These blots are representative of 5 and 3 independent plasma pools for HuA and MJFR14, respectively. aSyn was detected by mass spectrometry in both strain A and B IPs from brain but only in Strain B IP from plasma. Peptide sequences detected in Strain B plasma IP (E) and normalized spectra counts for 2D gel regions isolated and sent for proteomics identification are indicated (F).

**Supplementary Figure 10.**
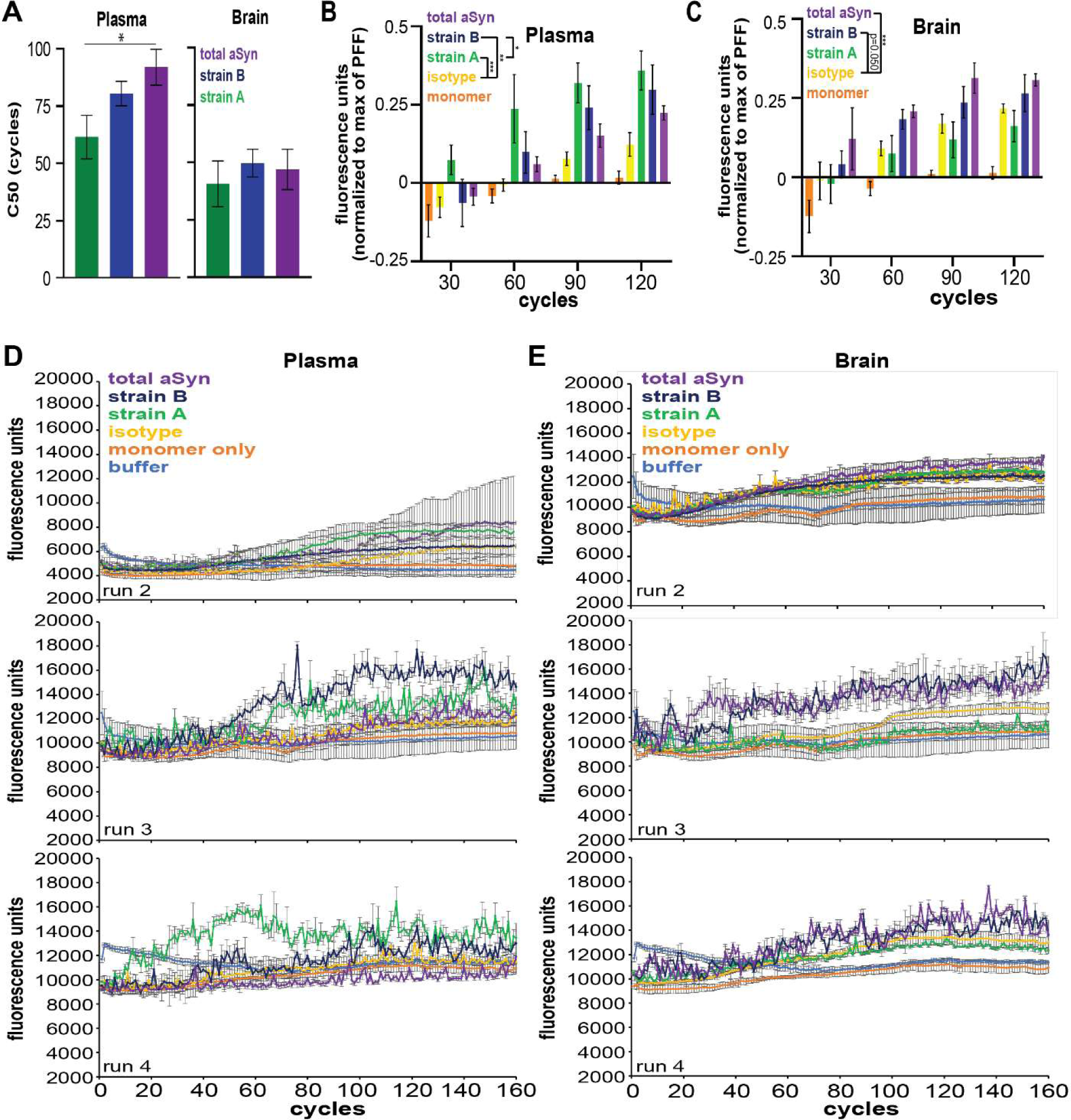
aSyn amplification curves from seed aggregation assays (SAA) using aSyn plasma and brain species in Parkinson’s disease from all performed runs. A total of 4 SAA replicates with aSyn species immunoprecipitated with strain-selective and total aSyn (syn211) antibodies from plasma (n = 2 pools) and caudate brain lysates (n = 3 pools) with PFF, monomer, and buffer only controls were performed. C50 (cycle number at which half of maximum relative fluorescence is reached) was significantly lower with strain A seed compared to total aSyn only in plasma (**A**). In plasma, quantification of fluorescence by normalization to PFF maximum fluorescence demonstrated significantly increased fibrillization of aSyn with plasma aSyn species enriched by both Strain A and B antibodies (**B**). In brain, aSyn species enriched by syn211 (total) aSyn antibody significantly increased fibrillization and species enriched by Strain B antibody trended to significantly increased fibrillization (p = 0.052, **C**). SAA curves for additional replicates are also displayed (**D, E**). Background fluorescence based on buffer only condition was subtracted from all values. Error bars represent SEM. PFF = pre-formed fibrils. * p < 0.05, ** p < 0.01, *** p < 0.001.

**Supplementary Figure 11.**
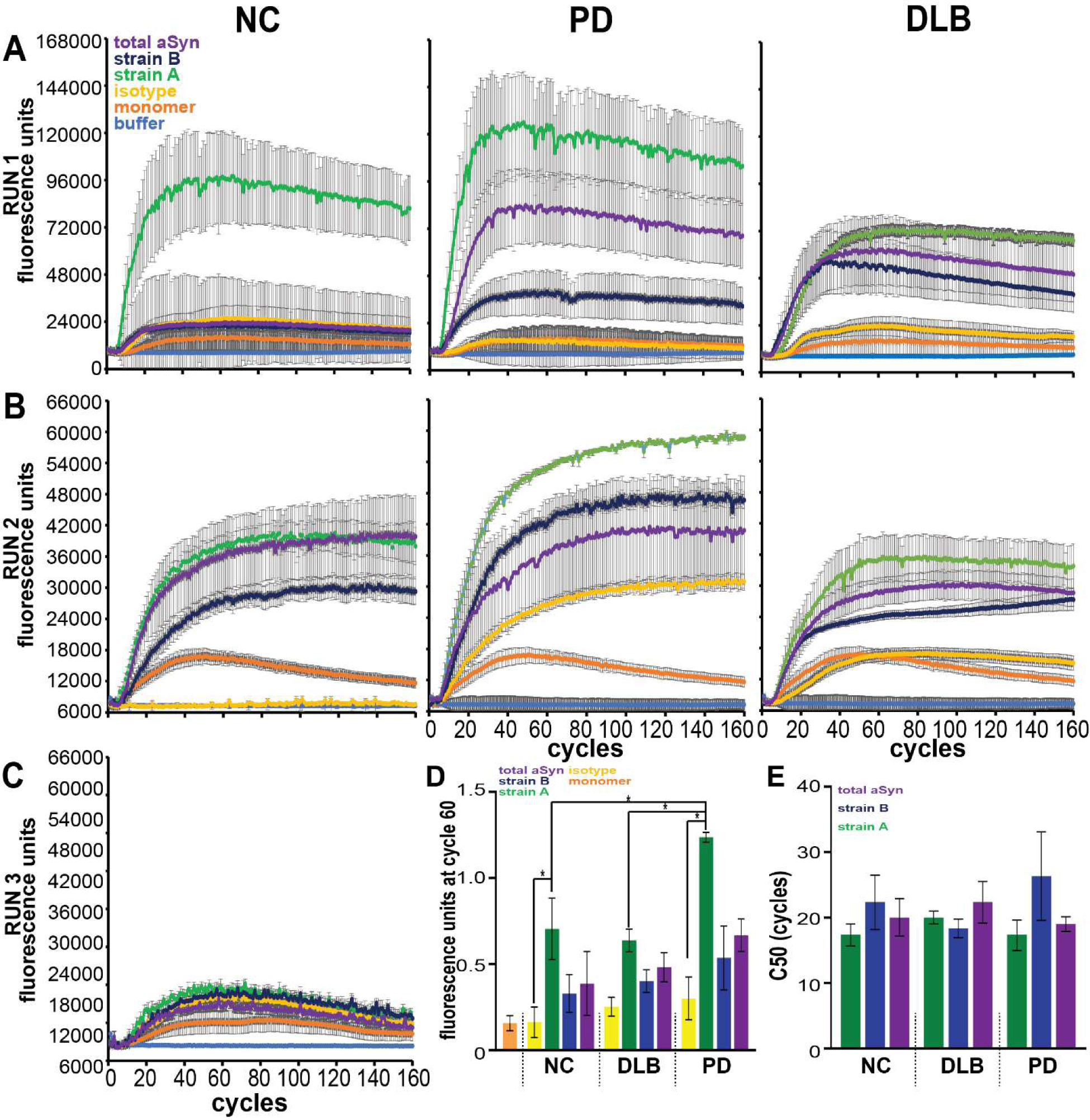
aSyn amplification curves from seed aggregation assays (SAA) using aSyn plasma species in normal controls (NC), dementia with Lewy bodies (DLB), and Parkinson’s disease (PD) from all performed runs. A total of 3 SAA replicates with aSyn species immunoprecipitated with strain-selective and total aSyn (syn211) antibodies from plasma (n = 3 pools per disease group) with preformed fibril, monomer, and buffer only controls were performed. SAA curves for additional replicates are displayed (**A-C**). Strain A aSyn species significantly induced aggregation in SAA relative to Ig control when quantified by normalizing fluorescence at cycle 60 to maximum PFF fluorescence (**D**). Strain A aSyn species from PD plasma induced more significant aggregation relative to species from NC or DLB plasma. C50 (cycle number at which half of maximum relative fluorescence is reached) was unchanged across strain or disease group (**E**). Background fluorescence based on buffer only condition was subtracted from all values. Error bars represent SEM. * p < 0.05.

**Supplementary Figure 12.**
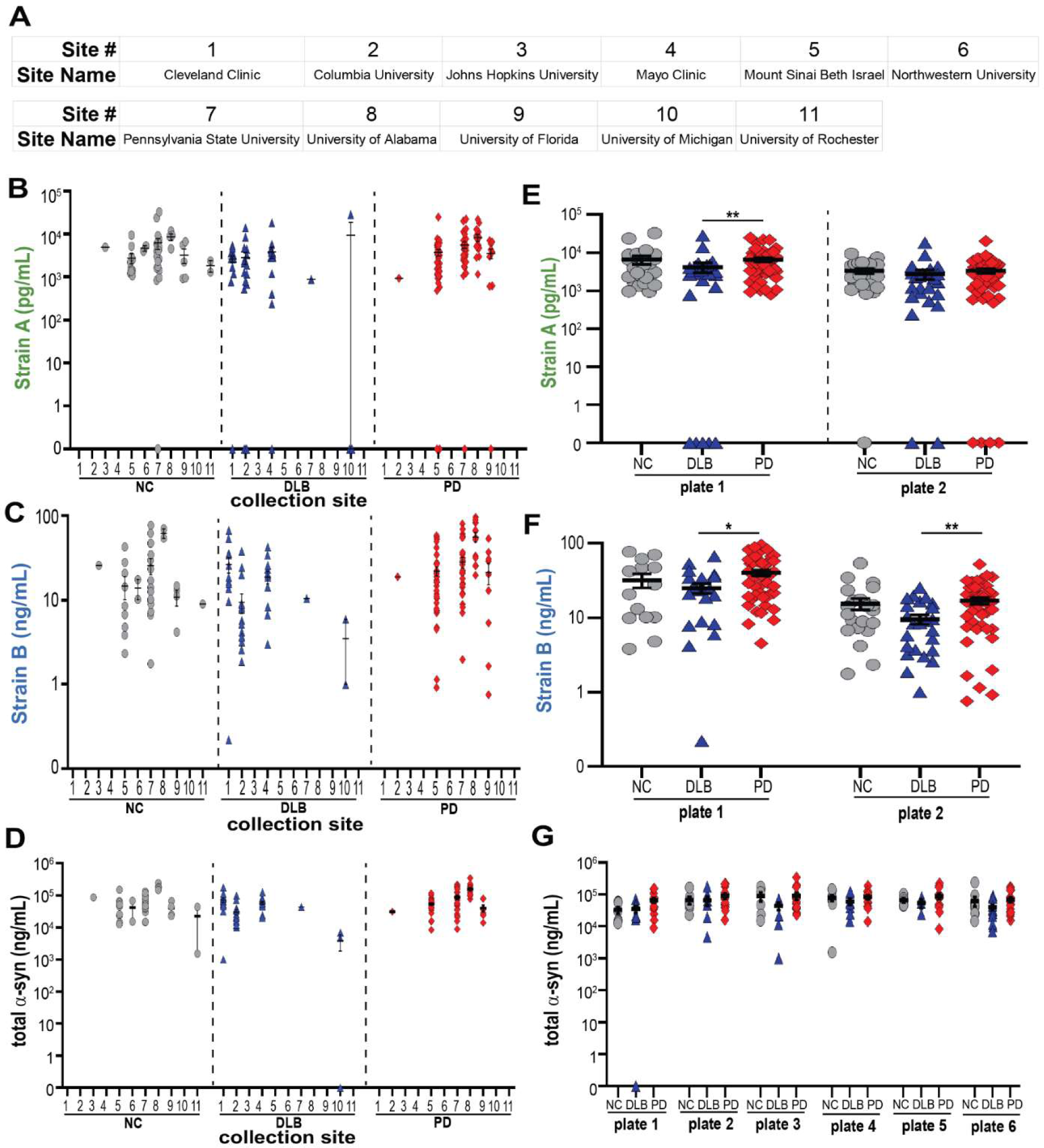
PDBP cohort measures show site-to-site and plate-to-plate variability, demonstrating need for normalization. The Parkinson’s Disease Biomarker Project (PDBP) cohort (n = 200) had plasma collected at 11 sites (**A**). Among sites, there was considerable site-to-site variability in plasma strain and total aSyn measurements (**B-D**). While each plate showed the same trends for PD vs. DLB, plate-to-plate variability was noted (**E-G**). For differences among groups, p-values (corrected for multiple comparisons for Kruskal-Wallis post-hoc test) for one-way ANOVA are reported in **E** and **F.** Error bars represent SEM. *p < 0.05, **p < 0.01. NC (grey circles) = normal controls, DLB (blue triangles) = dementia with Lewy bodies, PD (red diamonds) = Parkinson’s disease.

**Supplementary Figure 13.**
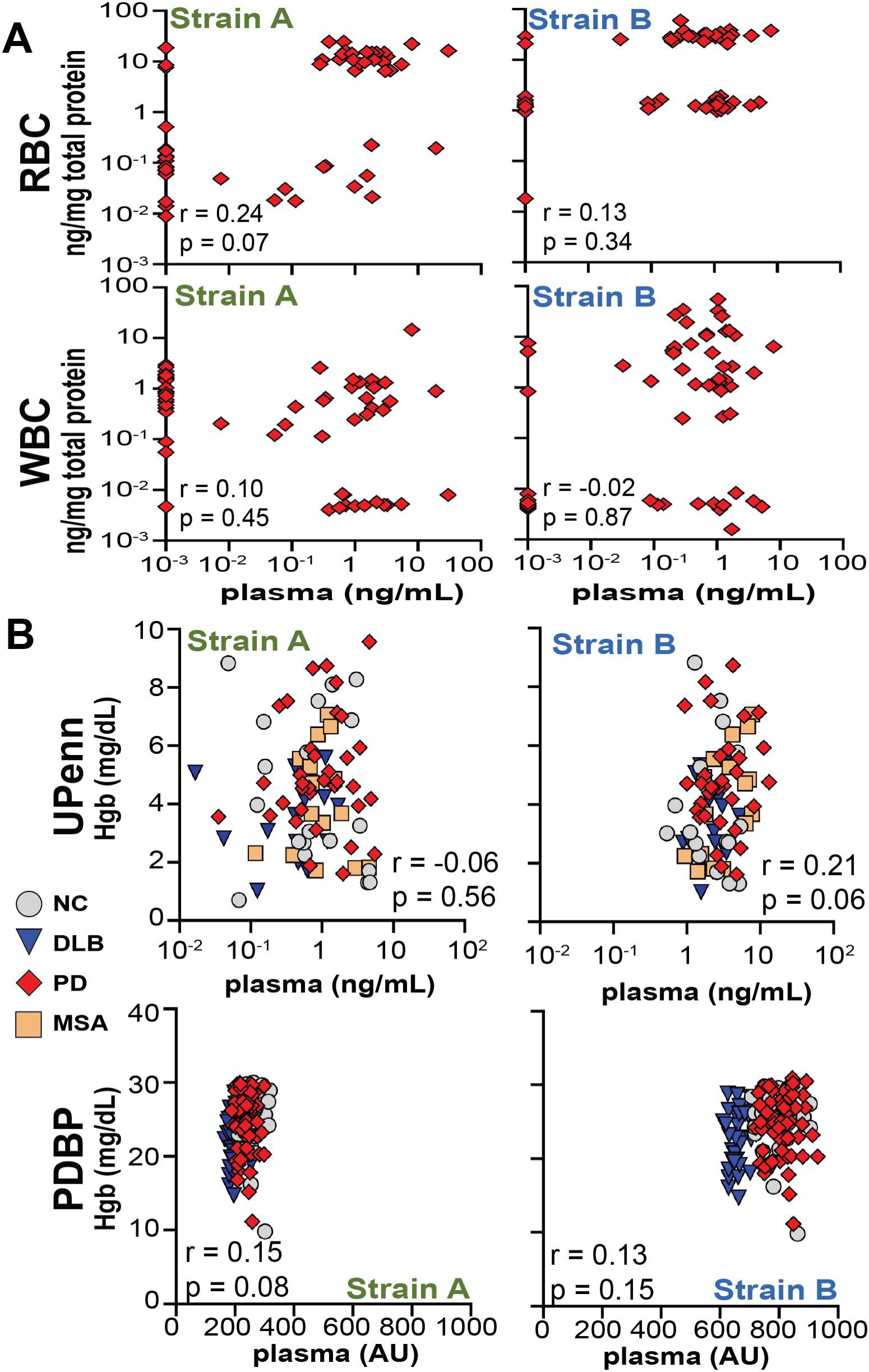
Plasma aSyn strain levels do not correlate with blood cell levels or hemolysis. Levels of aSyn strains in plasma do not correlate (Pearson) with levels in matched white or red blood cell fractions (n = 65) from individuals with PD as measured by ELISA (**A**). In both the University of Pennsylvania (n = 235) and the PD Biomarker Project (n = 200) cohorts, plasma aSyn strain levels measured by ELISA did not correlate with hemoglobin levels (Hgb) measured by absorbance. Between group differences were assessed by one-way ANOVA with Kruskal-Wallis post-hoc testing (**B**). Pearson correlation r and p-values are displayed (**A-B**). * p < 0.05. AU = arbitrary units, NC = normal control, RBC = red blood cells, WBC = white blood cells.

**Supplementary Table 1.**
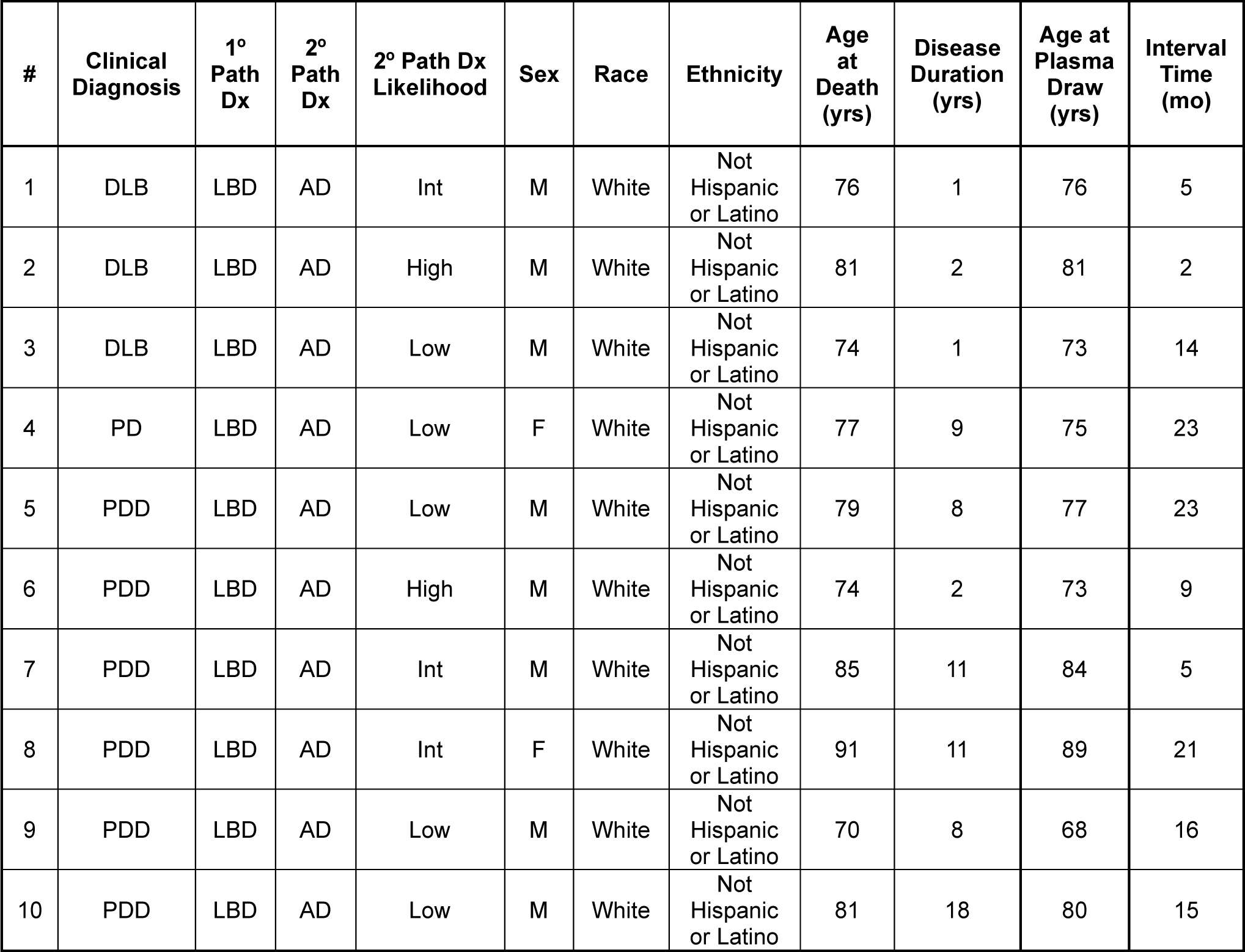
Samples for brain and matched plasma in Lewy boy disease (LBD). Summary of characteristics of individuals selected for the brain immunohistochemistry and comparison of aSyn strain quantification in brain lysate and plasma by ELISA. Interval time refers to the number of months between the plasma sample collection date and the date of death. DLB = dementia with Lewy bodies, PD = Parkinson’s disease, PDD = PD dementia, Int = Intermediate, Path Dx = Pathological Diagnosis.

**Supplementary Table 2.**
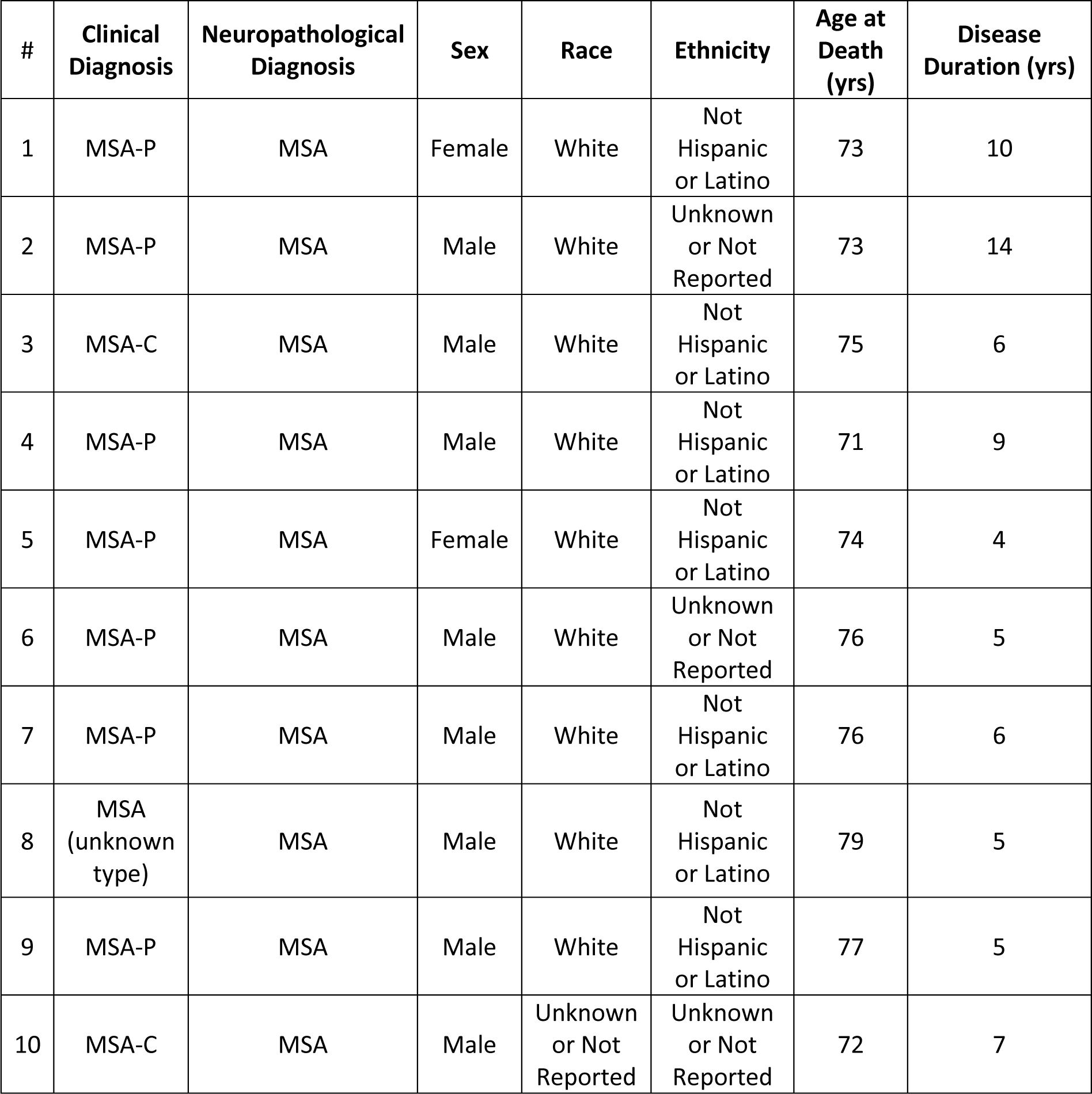
Samples for MSA brain immunohistochemistry. Summary of characteristics of individuals with MSA selected for the brain immunohistochemistry. MSA = multiple systems atrophy, MSA-P = MSA parkinsonian type, MSA-C = MSA cerebellar type

**Supplementary Table 3.**
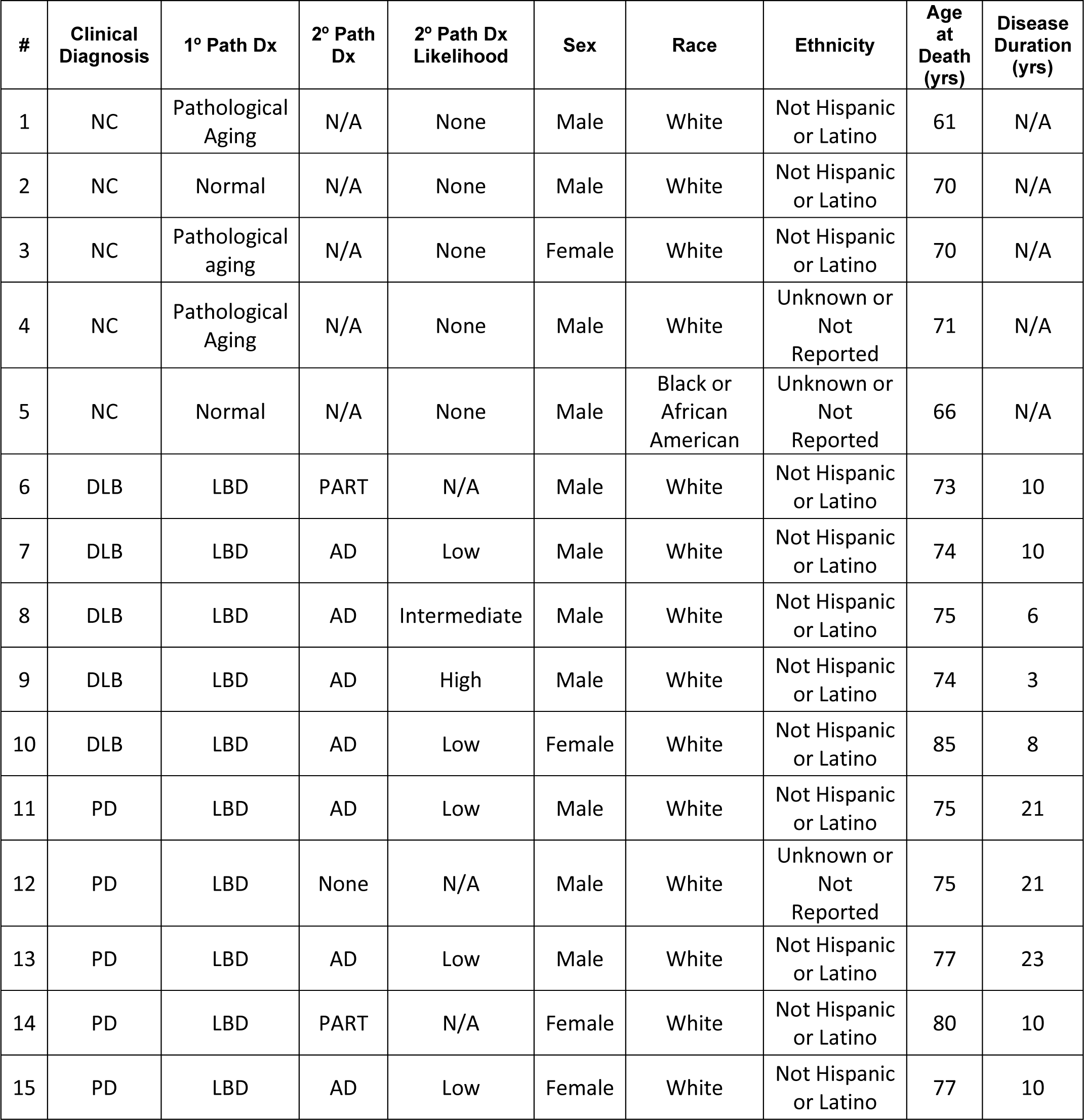
ELISA and Western Blot Comparison Samples. Characteristics of individuals selected for the comparison of aSyn strains by ELISA and Western Blot. Path Dx = Pathological Diagnosis, NC = normal control, DLB = dementia with Lewy bodies, LBD = Lewy Body Disease, PD = Parkinson’s disease, AD = Alzheimer’s disease, N/A = not applicable.

**Supplemental Table 4.**
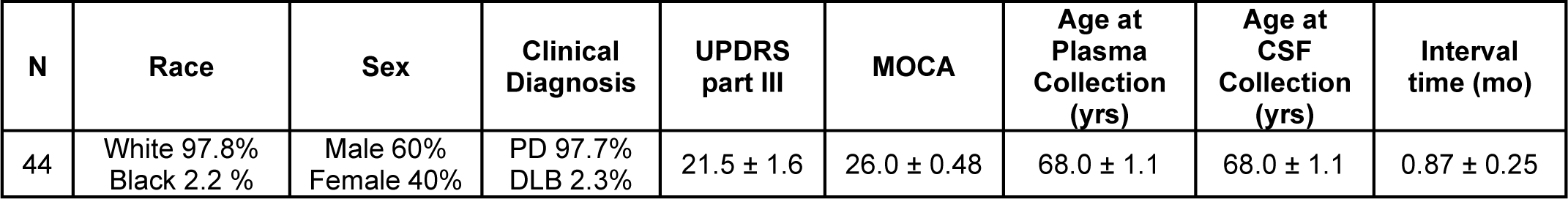
Matched CSF and Plasma Samples. Summary of characteristics of individuals selected for the comparison of CSF and Plasma values on ELISA listed as mean ± standard error of the mean or frequency (%). Interval time refers to the number of months between the plasma sample date and the CSF sample date.

**Supplementary Table 5.**
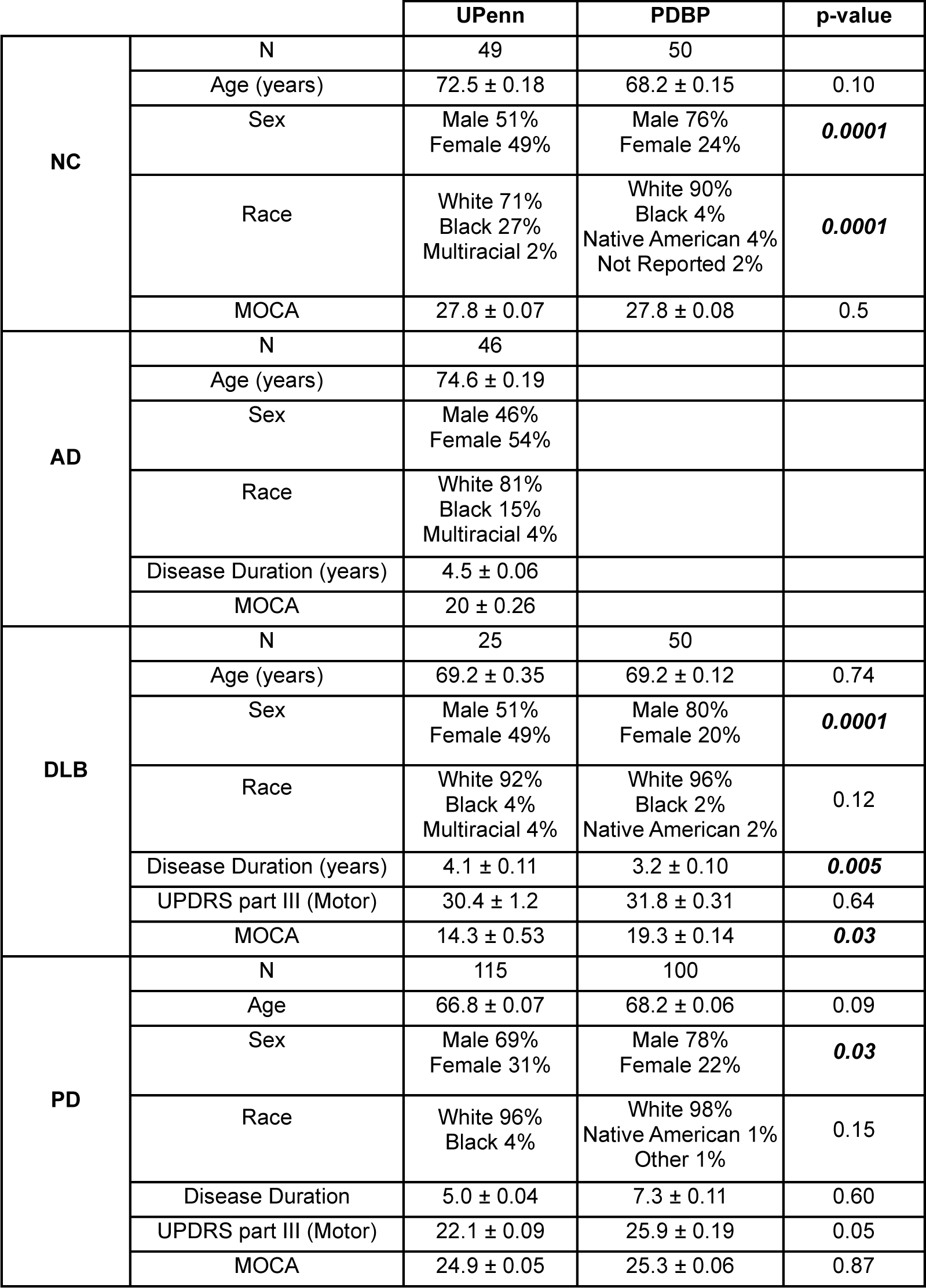
UPenn and PDBP Plasma Cohorts. Characteristics of UPenn and PDBP cohorts used for plasma and clinical analyses listed as mean ± standard error of the mean or frequency (%). P-values represent Mann-Whitney tests and chi-square tests for continuous variables or proportions, respectively.

**Supplementary Table 6.**
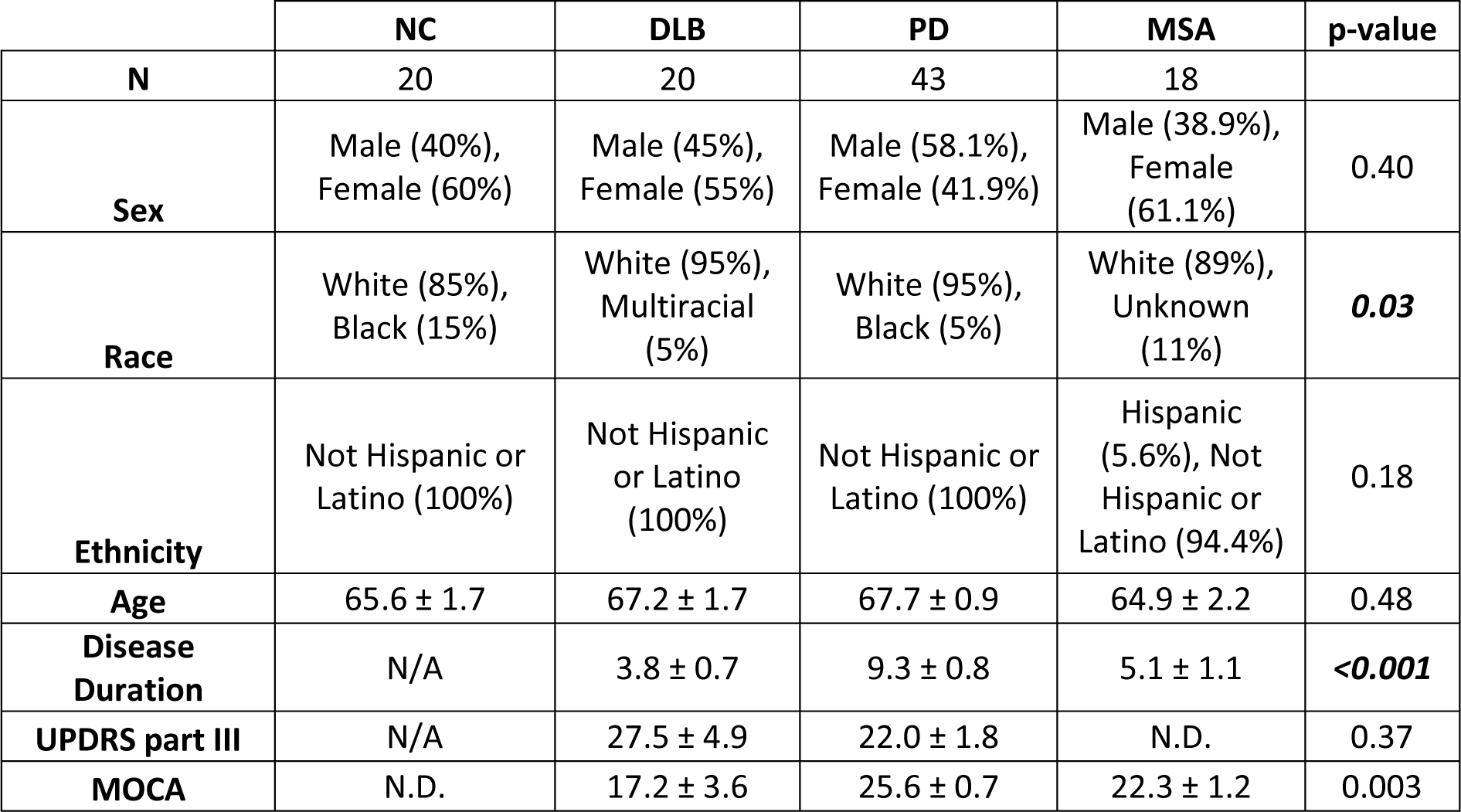
Characteristics of Plasma Samples for comparison of aSyn strains in plasma from normal controls (NC), dementia with Lewy bodies (DLB), Parkinson’s disease, and Multiple Systems Atrophy (MSA). Characteristics of plasma samples listed as mean ± standard error of the mean or frequency (%). P-values represent Kruskal-Wallis tests and Fisher exact tests for continuous variables or proportions, respectively. N/A = not applicable, N.D. = no data.

**Supplementary Table 7.**
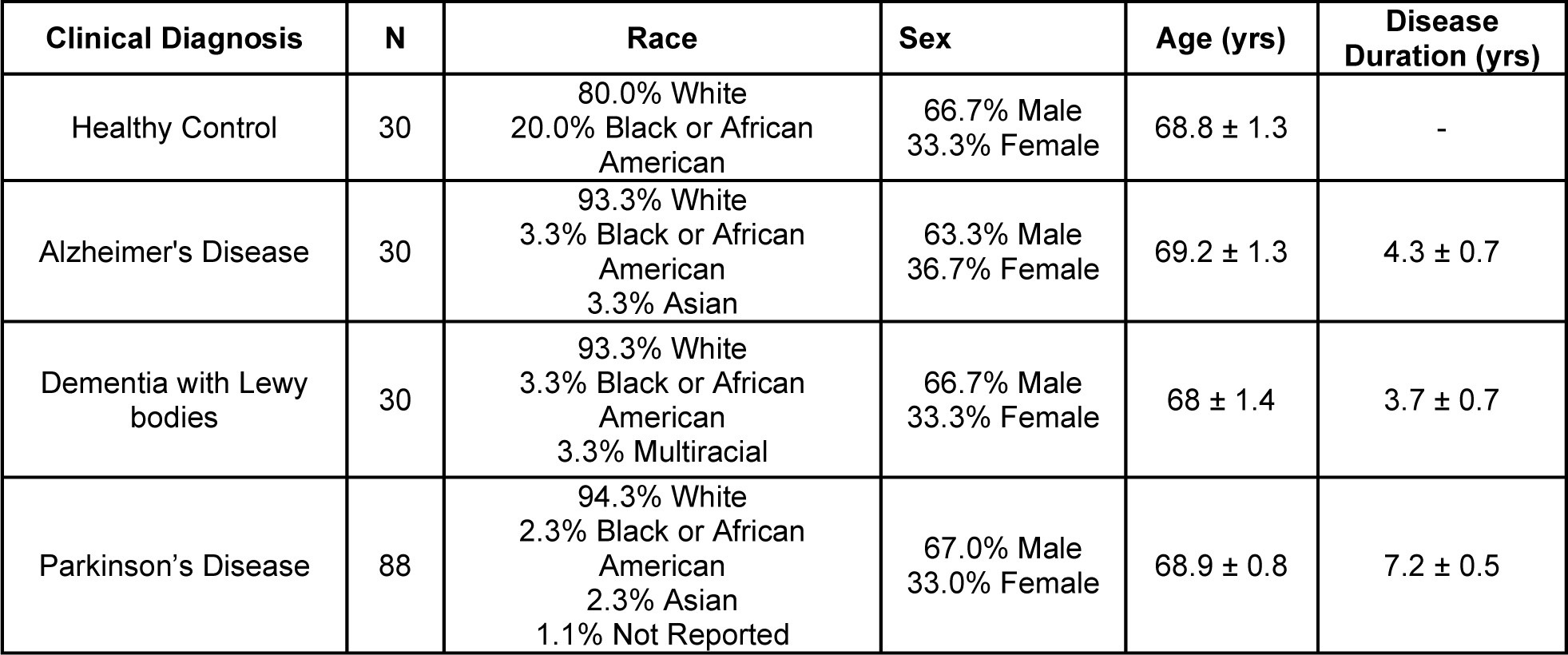
Total α-synuclein ELISA Samples. Characteristics of individuals selected for analysis on total α-synuclein ELISA, listed as mean ± standard error of the mean or frequency (%).

**Supplementary Table 8.**
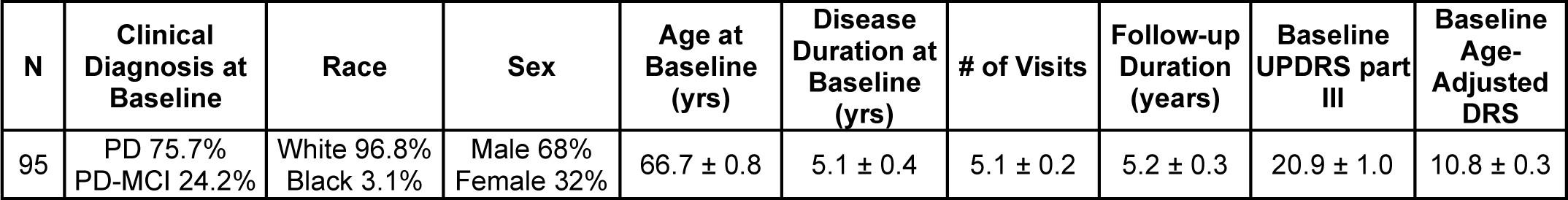
Samples for Motor and Cognitive Linear Mixed-Effects Model. Characteristics of individuals selected for motor and cognitive linear mixed-effects model listed as mean ± standard error of the mean or frequency (%). Cognitive status established by consensus diagnosis.

**Supplemental Table 9.**
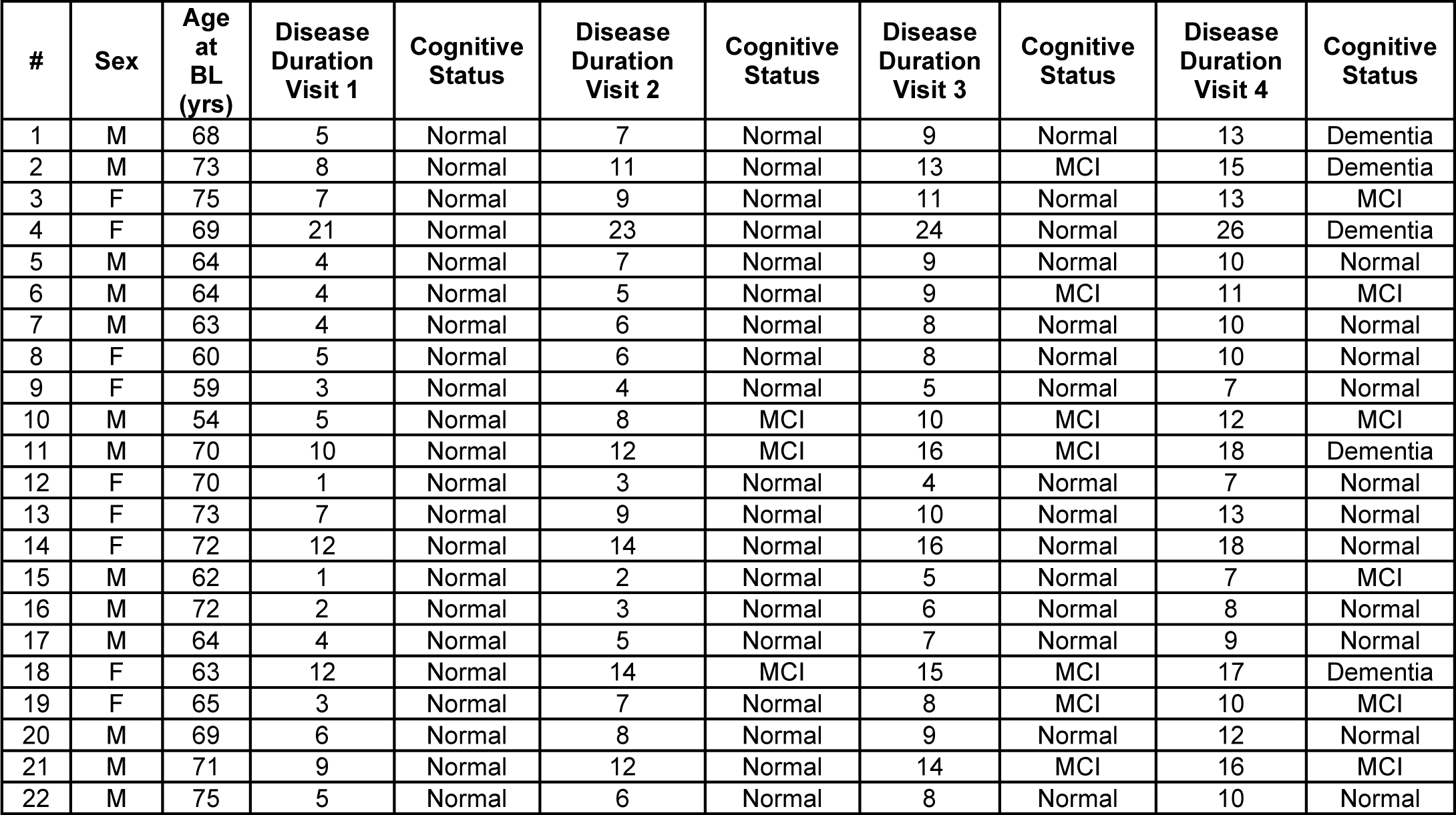
Longitudinal Plasma ELISA Samples. Characteristics of individuals selected for the longitudinal plasma cohort whose aSyn strain levels were measured on the ELISA. All individuals were of white race. Disease duration reported in years. Cognitive status established by consensus diagnosis.

**Supplementary Table 10.**
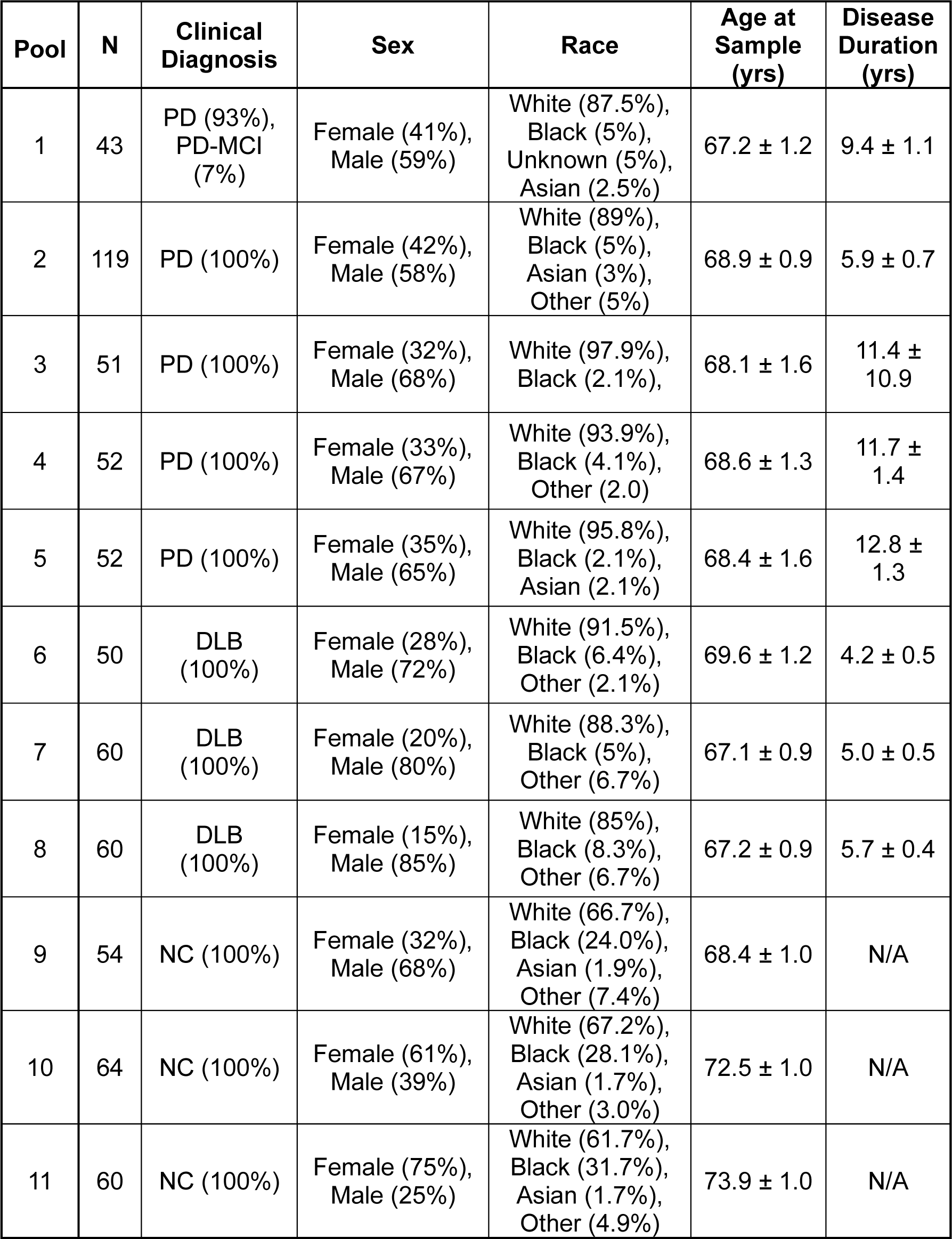
Pooled Plasma Immunoprecipitation Samples. Characteristics of individuals selected for pooled plasma for immunoprecipitation listed as mean ± standard error of the mean or frequency (%).

**Supplementary Table 11.**
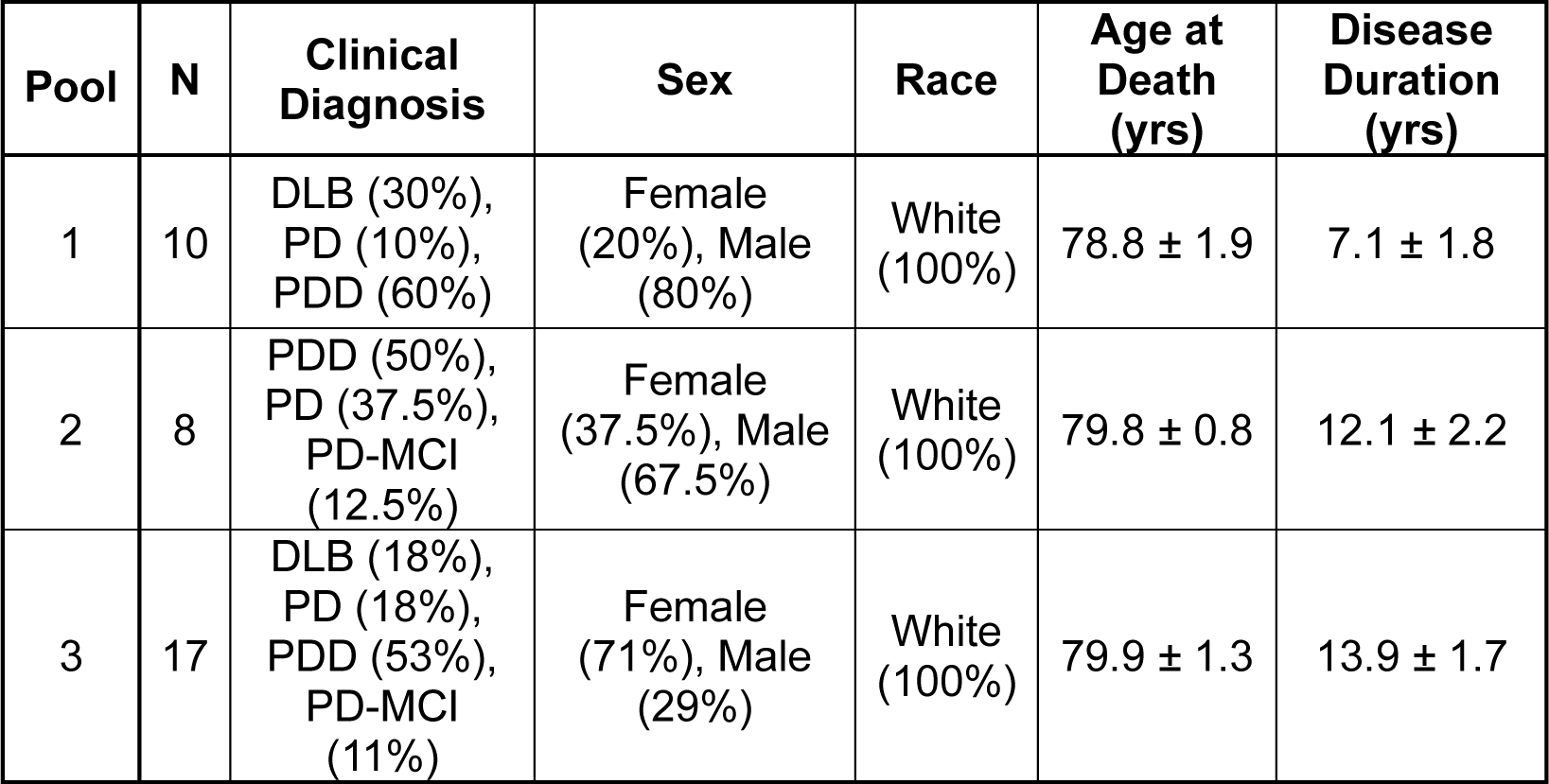
Pooled Brain Lysate Immunoprecipitation Samples. Characteristics of individuals selected for pooled brain caudate lysate for immunoprecipitation listed as mean ± standard error of the mean or frequency (%).

**Supplementary Table 12.**
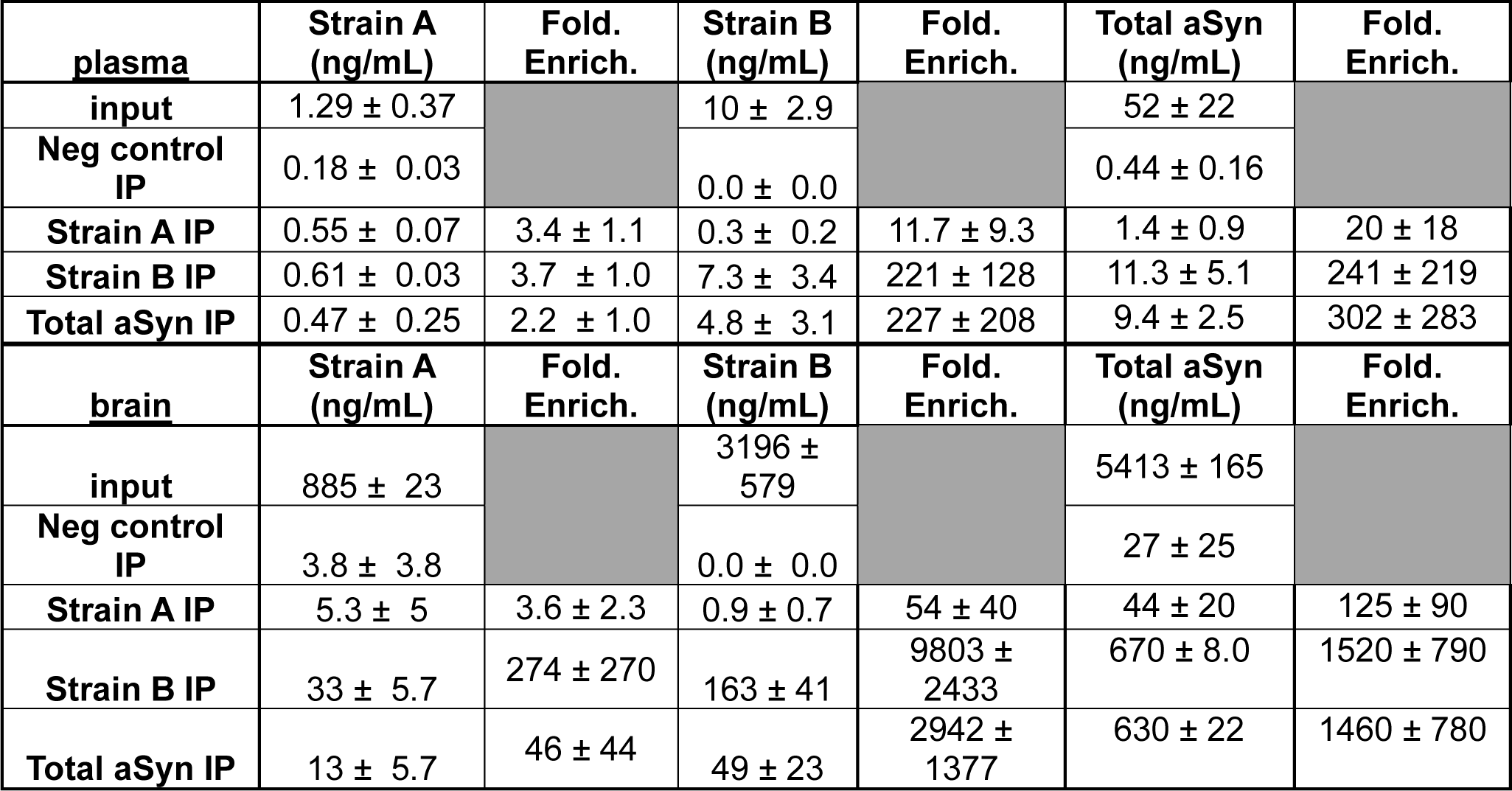
Concentration of aSyn Isolated by IP from Plasma and Brain with Strain-selective Antibodies. Concentration of aSyn in eluates isolated by strain-selective or total aSyn antibody IP are represented by the mean ± standard error of the mean of three brain IPs and six plasma IPs. The negative control IP was performed with isotype-matched IgG antibody. Fold enrichment calculated based on half of lower limit of detection if negative (neg) control IP ELISA concentration was below limit of detection. IP = immunoprecipitation.

**Supplementary Table 13.**
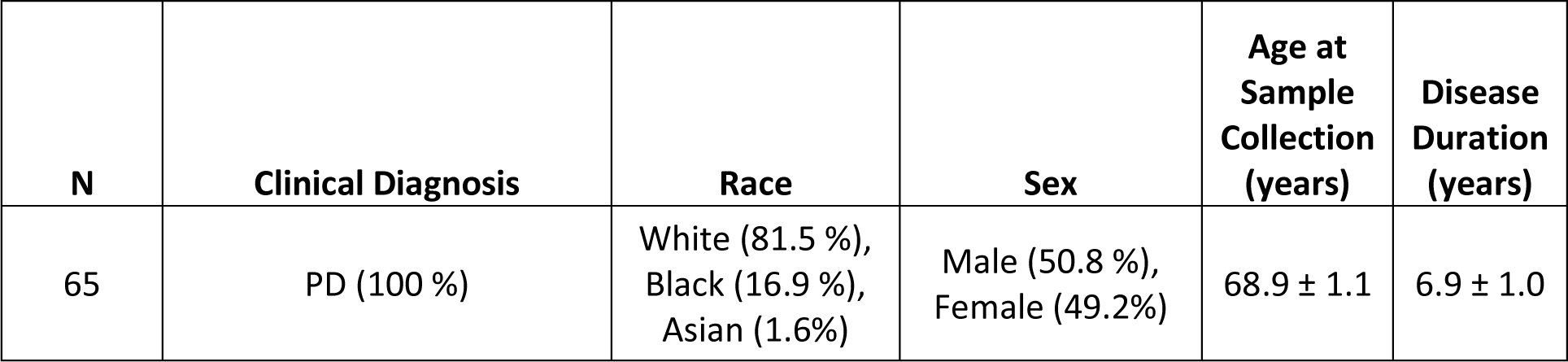
Parkinson’s disease (PD) cohort for matched blood cell and plasma samples. Characteristics of PD cohorts used for measurement of aSyn strain levels in matched blood cell and plasma samples listed as mean ± standard error of the mean or frequency (%).

**Supplementary Table 14.**
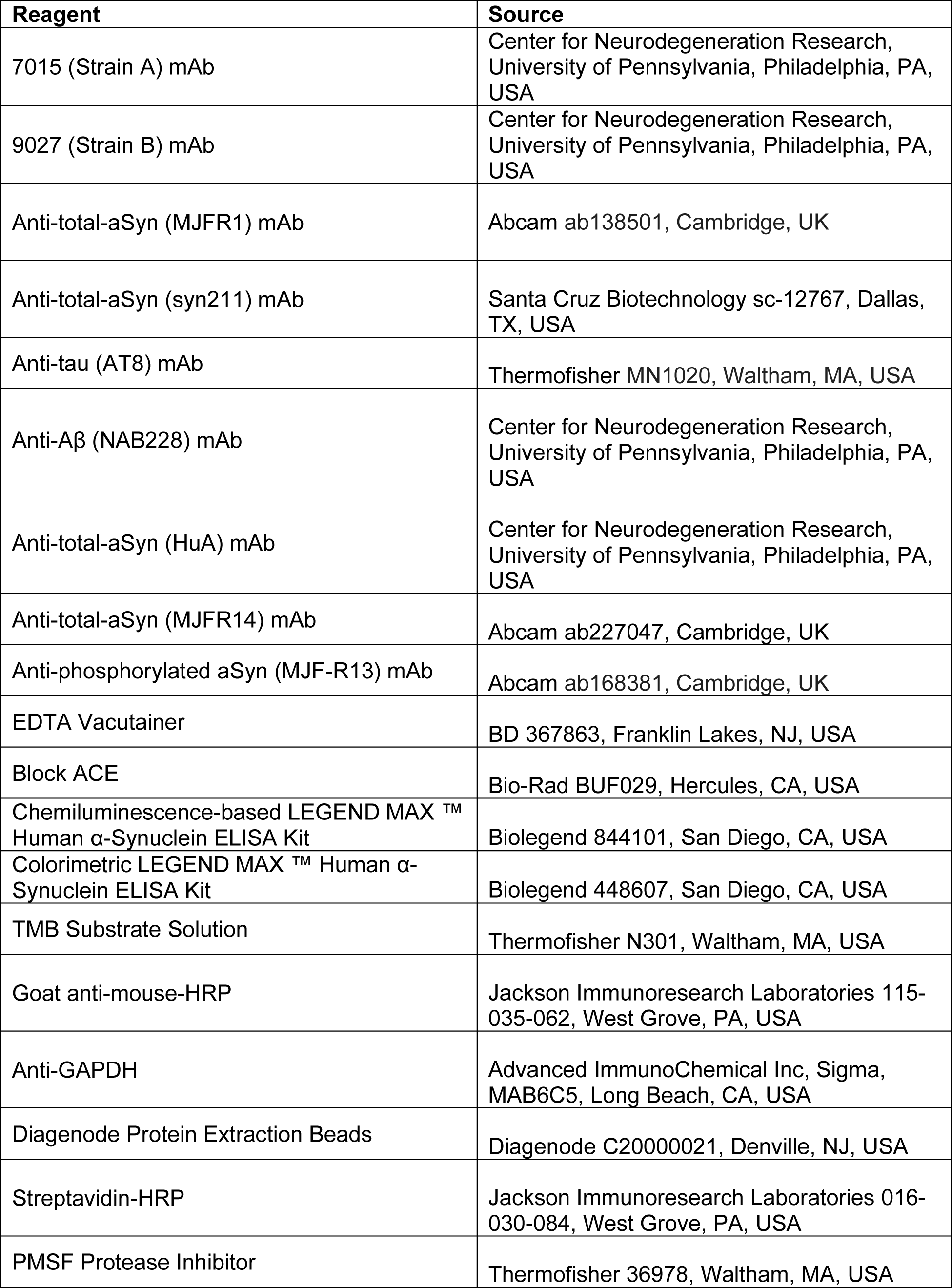
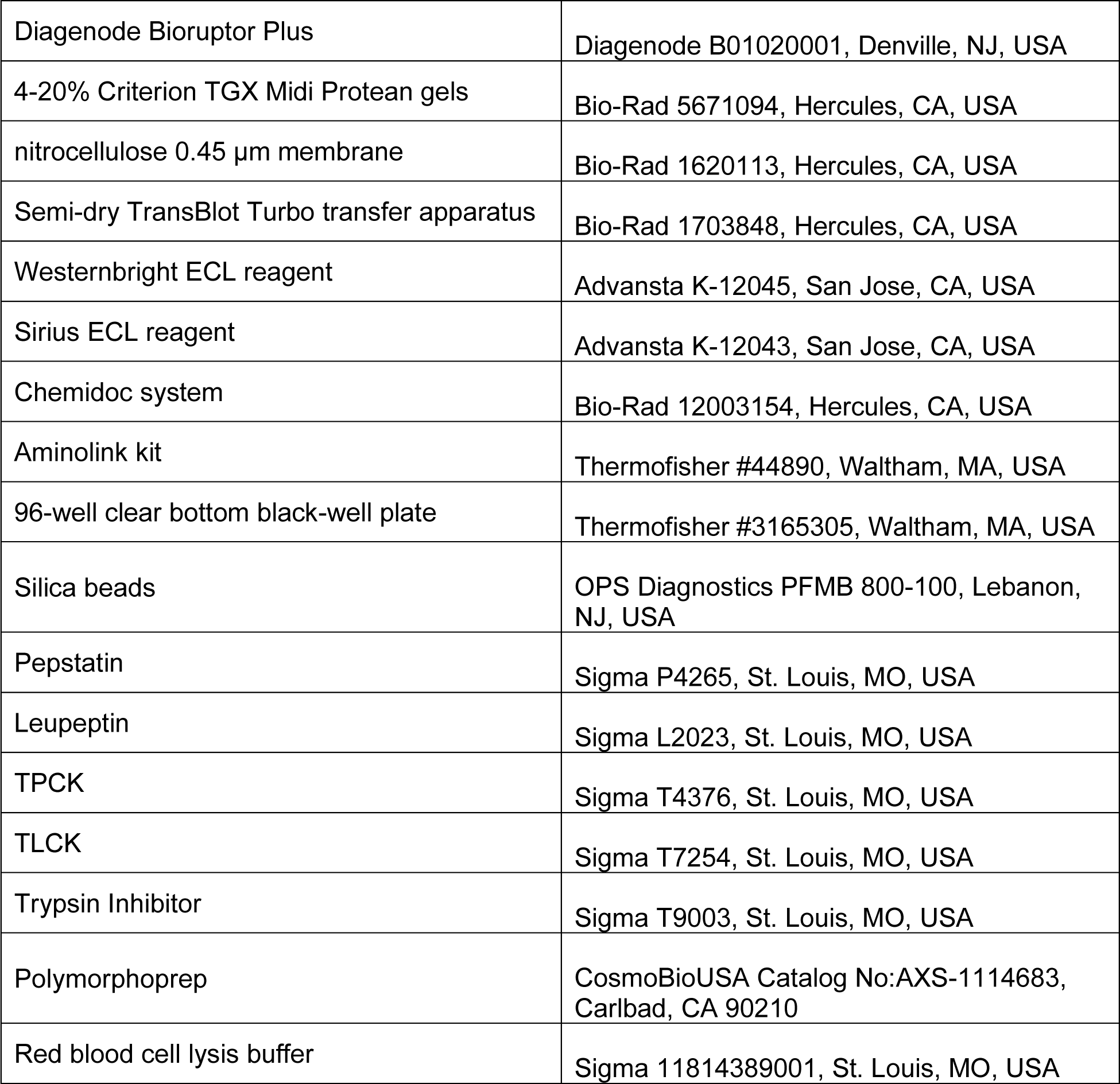
Antibodies and reagents used in this manuscript.

