## Supplementary Figures and Tables for "α-Synuclein Conformations in Plasma Distinguish Parkinson’s Disease from Dementia with Lewy Bodies"

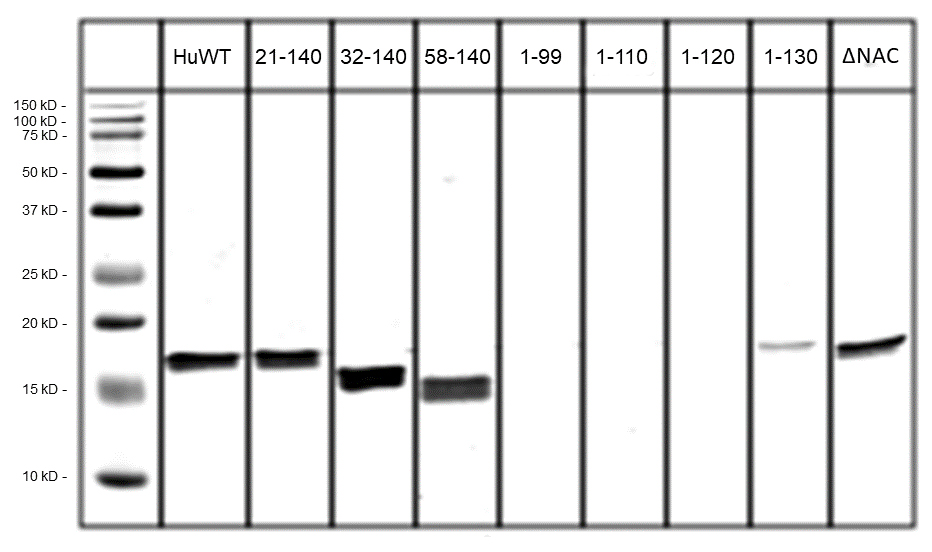

**Supplementary Figure 1. Epitope map for monoclonal 9027 antibody generated against strain B.** Immunoblotting against full-length human wildtype (HuWT) aSyn and indicated peptide fragments demonstrate that the 9027 antibody detects a continuous region on aSyn between residues 120-140. ΔNAC = NAC domain deletion mutant.

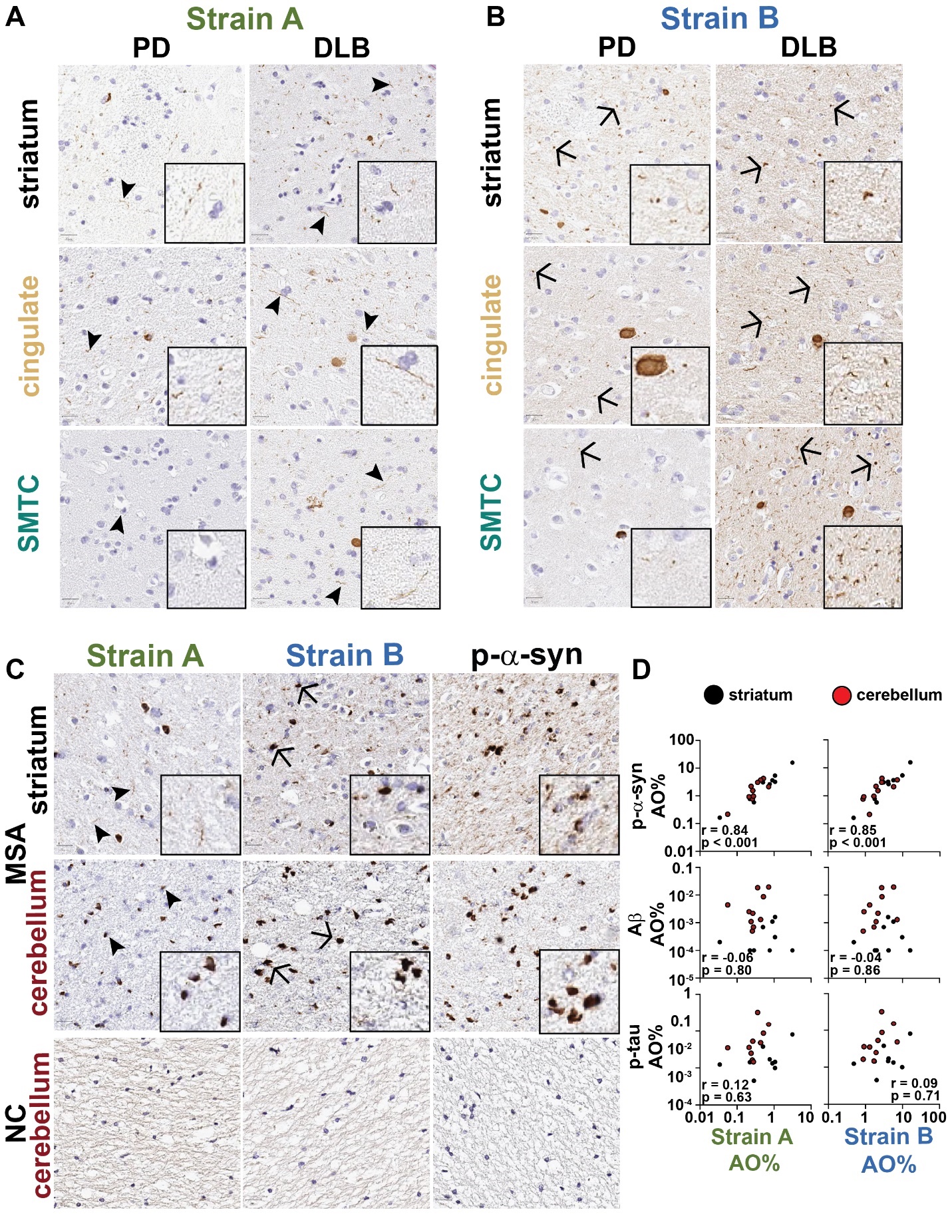

**Supplementary Figure 2. Strain-selective antibodies detect overlapping but distinct subsets of pathology across PD, DLB, and MSA brains.** Representative images of strain A (**A**) and strain B (**B**) immunohistochemical staining are shown for PD (A) and DLB from the set of 10 individuals with LBD in Fig 1. Representative images of strain A, strain B, and p-aSyn immunohistochemical staining are shown from a set of 10 individuals with MSA (**C**). aSyn strain staining correlates strongly with p-aSyn in MSA brains but not with Ab or p-tau staining in both striatum (black) and cerebellum (red, **D**). AO = area occupied, DLB = dementia with Lewy bodies, PD = Parkinson’s disease, PDD = PD dementia, SMTC = superior middle temporal cortex.

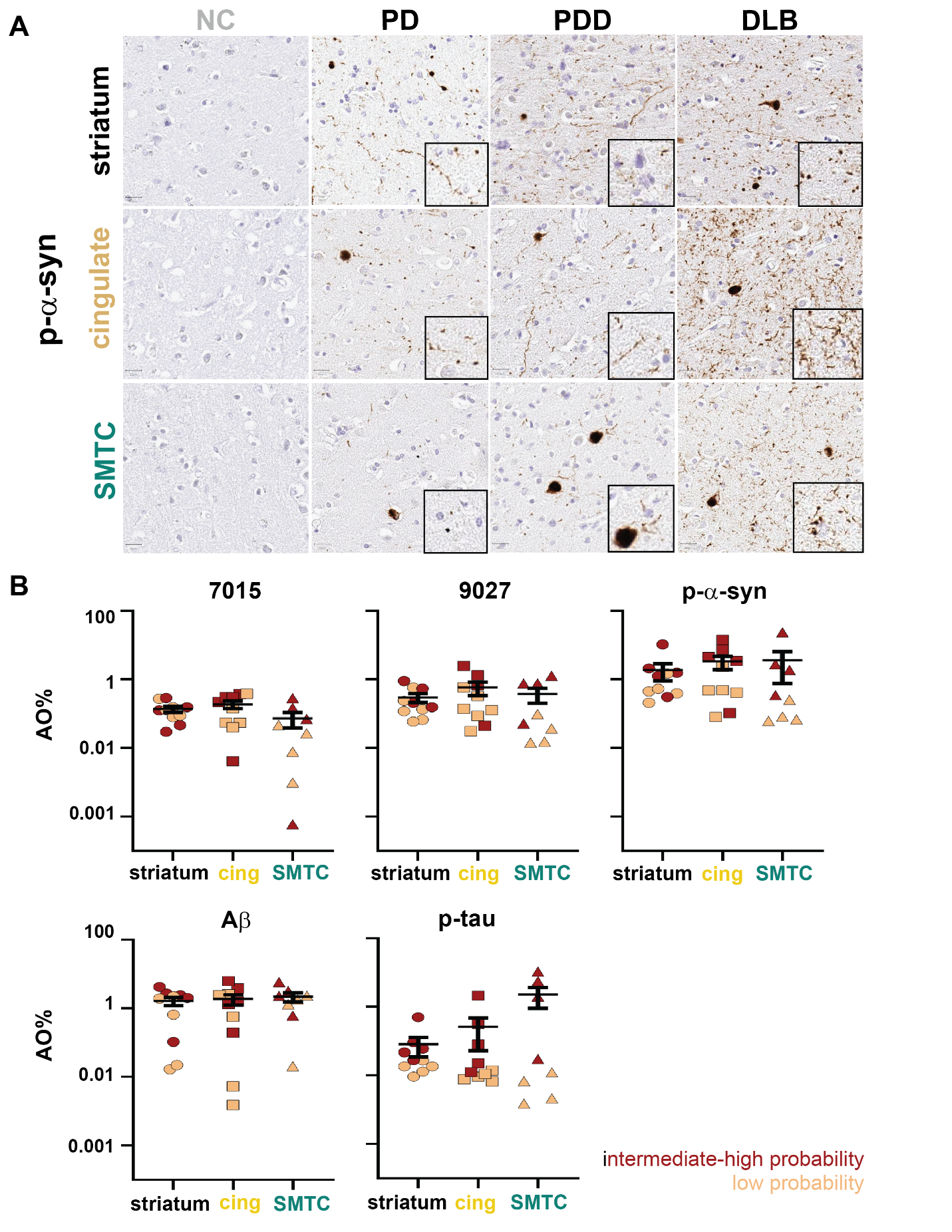

**Supplementary Figure 3. Strain B immunostaining is higher in individuals with higher** **levels of Alzheimer’s disease co-pathology.** Representative images of p-aSyn immunohistochemical staining are shown (**A**) from a set of three NC and ten individuals with LBD. The levels of strain A, strain B, p-aSyn, Aβ, and p-tau stratified by intermediate/high (red) or low (tan) probability of secondary AD neuropathologic diagnosis across striatum (circles), anterior cingulate cortex (squares), and SMTC (triangles) are shown (**B**), indicating higher AD co-pathology in areas with higher levels of Strain B antibody staining. Line represents mean and error bars represent SEM. AO = area occupied, cing = anterior cingulate cortex, SMTC = superior middle temporal cortex.

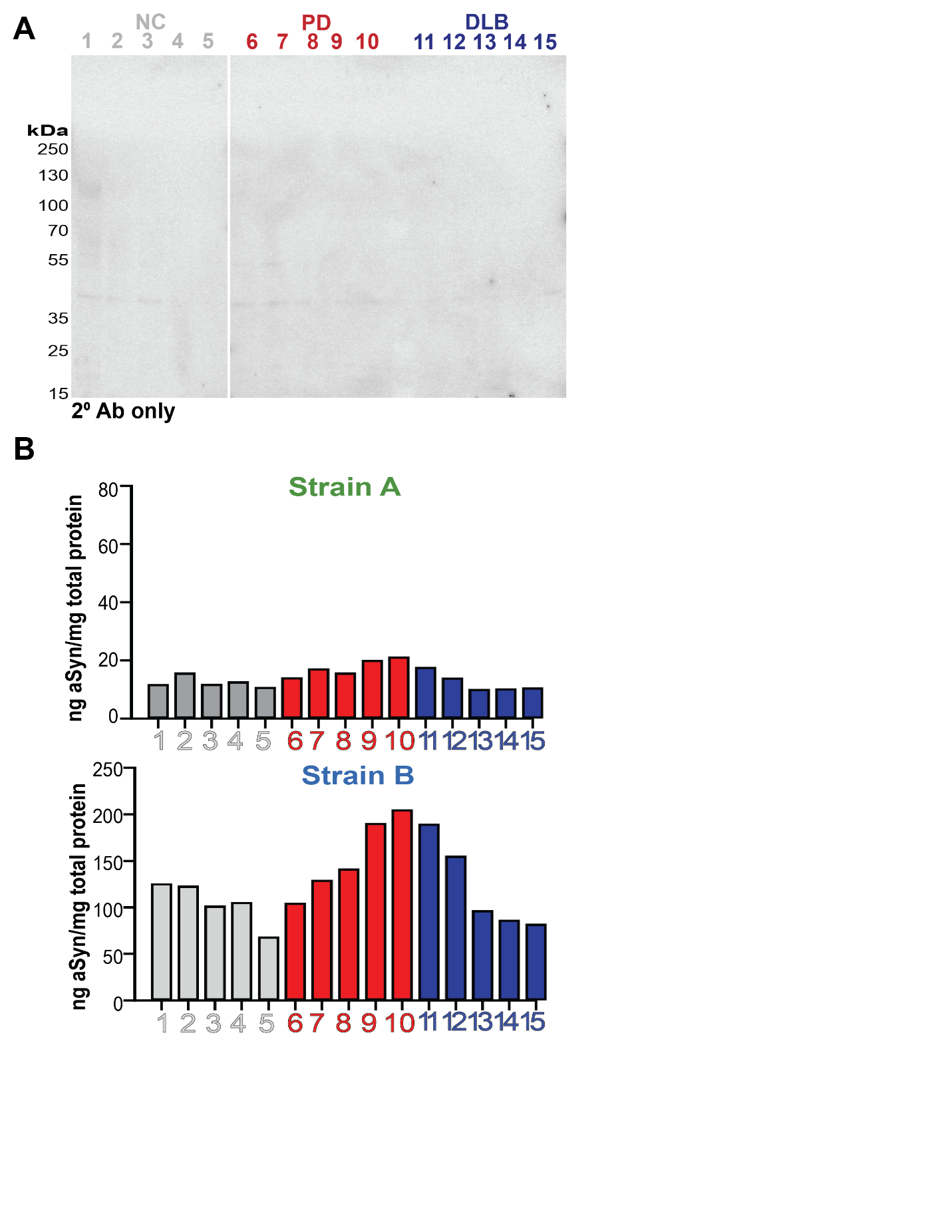

**Supplementary Figure 4. aSyn strain ELISAs detect pathologic forms of aSyn.** Western blot of caudate brain lysates from healthy controls (NC), individuals with Parkinson’s disease (PD), and individuals with dementia with Lewy bodies (DLB) with secondary antibody only does not show significant off-target binding (**A**). Levels of aSyn strain levels in cerebellum brain lysates in individuals with PD or DLB were lower than detected in caudate (Fig. 2E-F), a region relatively enriched for aSyn pathology.

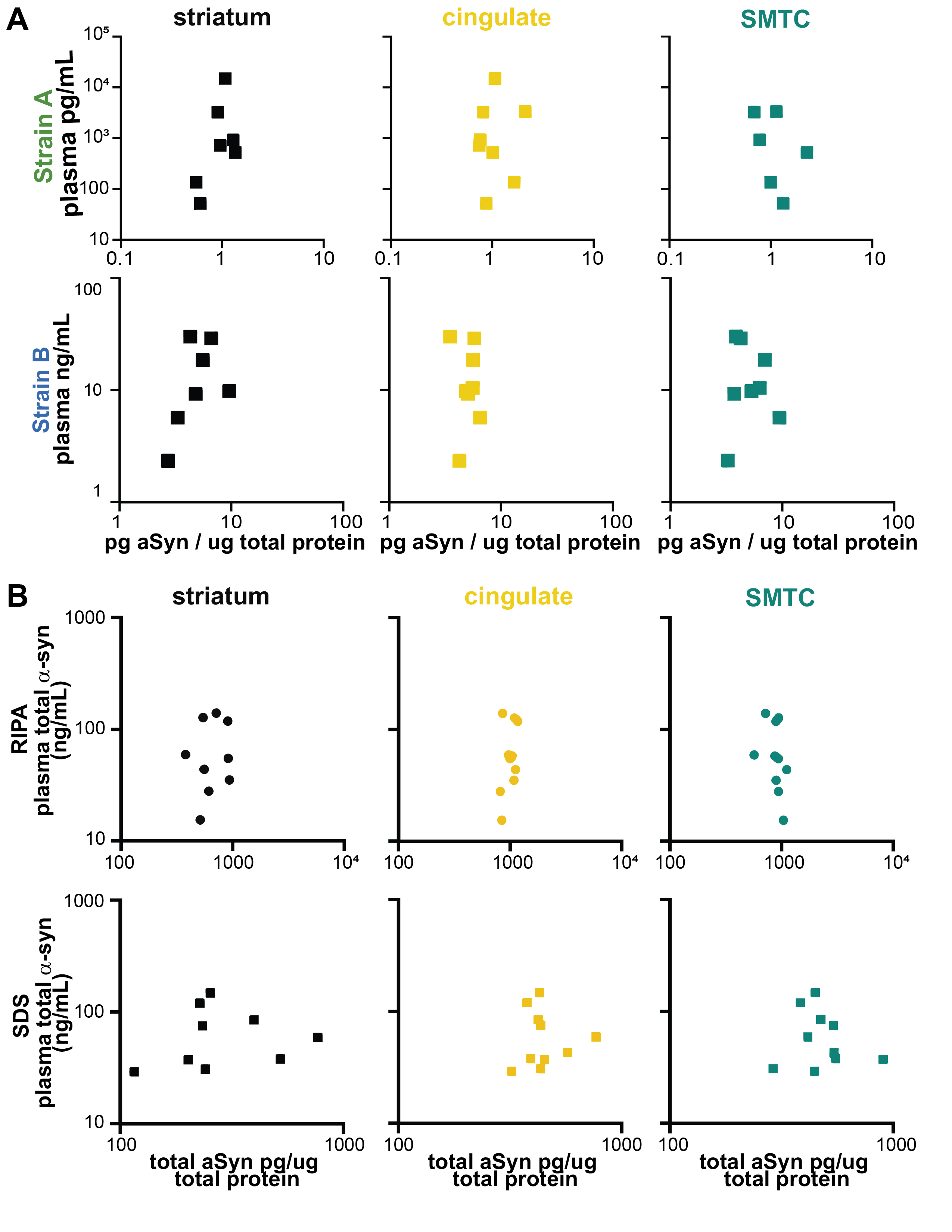
 **Supplementary Figure 5. Strain and total aSyn plasma levels do not correlate with brain levels.** For individuals with plasma collected within two years of autopsy (n = 10), plasma strain aSyn levels do not correlate with levels in brain caudate lysates extracted in 2% SDS buffer in regions associated with diffuse cortical spread of Lewy pathology (**A**). Total plasma aSyn levels also do not correlate with brain lysate levels in RIPA or SDS fractions (**B**). SDS = sodium dodecyl sulfate, SMTC = superior middle temporal cortex.

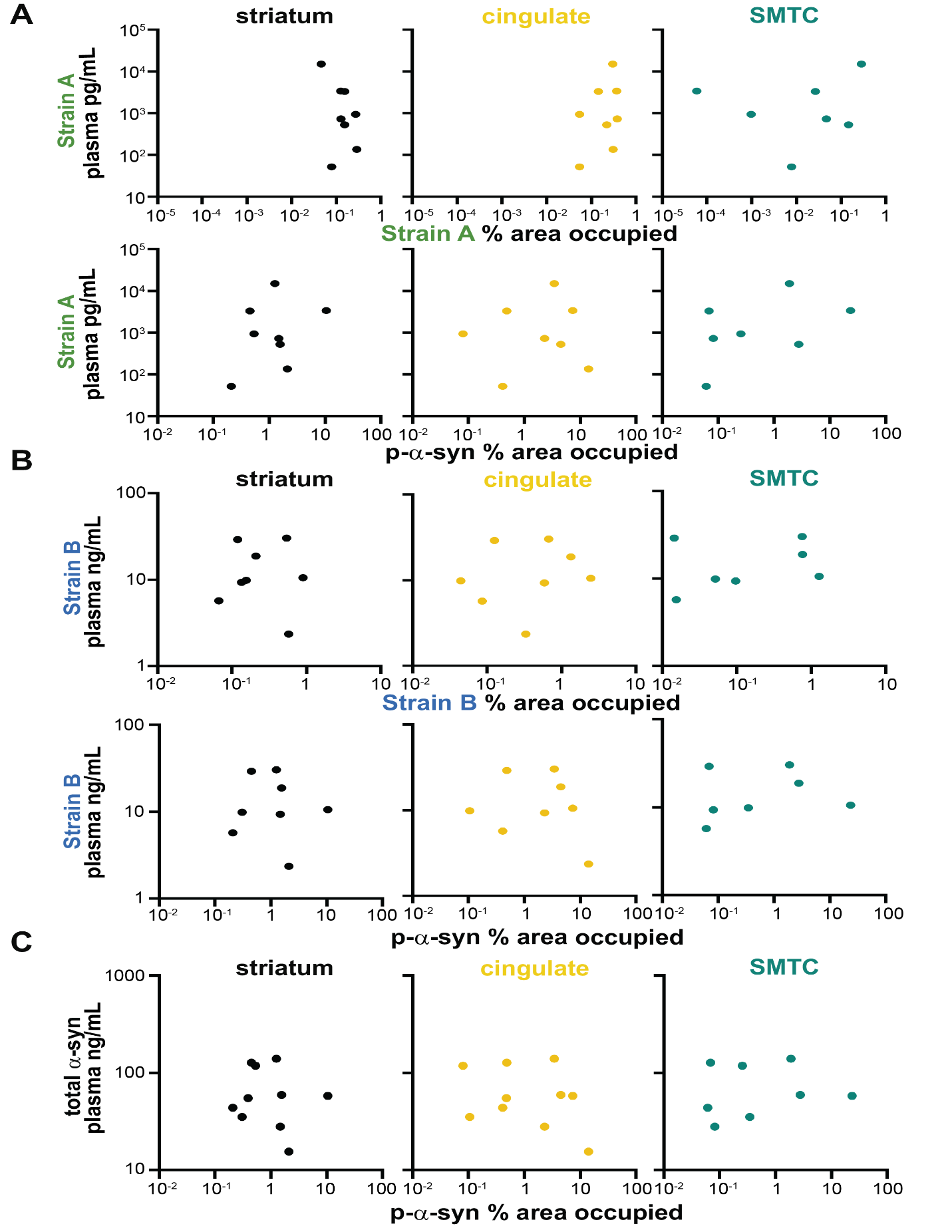
 **Supplementary Figure 6. Plasma levels of total and strain aSyn measured by ELISA do not correlate with brain immunohistochemistry.** For plasma samples collected within two years of autopsy (n = 10), plasma strain (**A, B**) or total (**C**) aSyn levels do not correlate with levels of immunohistochemical staining in regions associated with diffuse cortical spread of Lewy pathology. SMTC = superior middle temporal cortex.

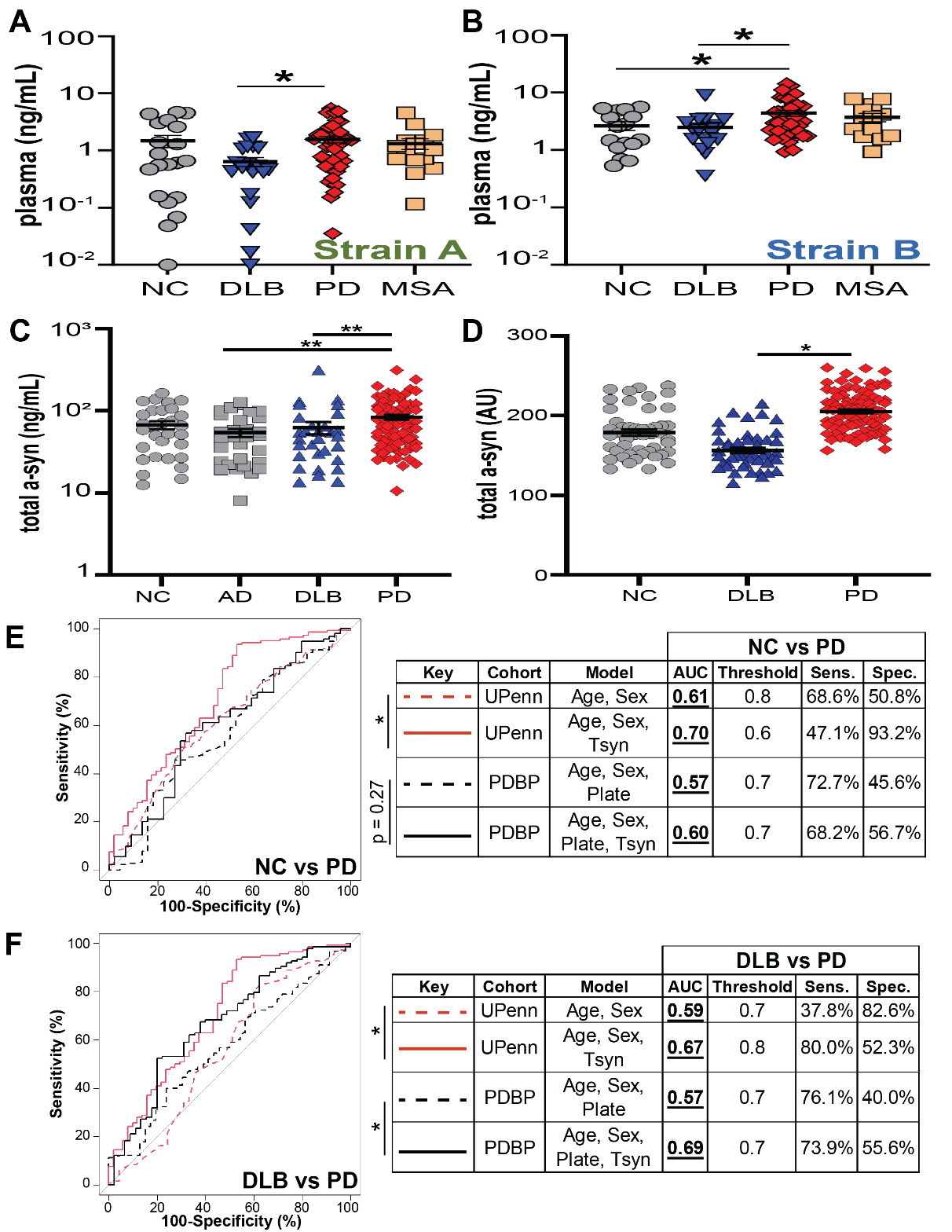

**Supplementary Figure 7. Plasma aSyn strain levels do not differ between individuals with Parkinson’s disease (PD) and multiple systems atrophy (MSA) and total plasma aSyn levels do not robustly differentiate PD and dementia with Lewy bodies (DLB).** Measurement of aSyn strain levels (n = 101) by ELISA reproduced the decrease previously observed in plasma from dementia with Lewy bodies relative to PD but levels were similar between PD and MSA plasma (**A, B**)**.** Total plasma aSyn levels were measured by a commercially available ELISA in the UPenn (**C**) and PDBP cohorts (**D**). ROC curves were used to differentiate PD vs. NC (**E**) or PD vs. DLB (**F**) incorporating age, sex, and plate with or without plasma total aSyn strain levels, as predictors. In all cases, incorporation of total aSyn plasma measures minimally improved discrimination. Thresholds generated by the Youden method. Error bars represent SEM. For differences among groups, p-values (corrected for multiple comparisons for Kruskal-Wallis post-hoc test) for one-way ANOVA are reported in **A** and **B.** Significance testing for differences between ROC curves was performed using the DeLong method. AU = arbitrary units, NC (grey circles) = normal controls, DLB (blue triangles) = dementia with Lewy bodies, PD (red diamonds) = Parkinson’s disease.

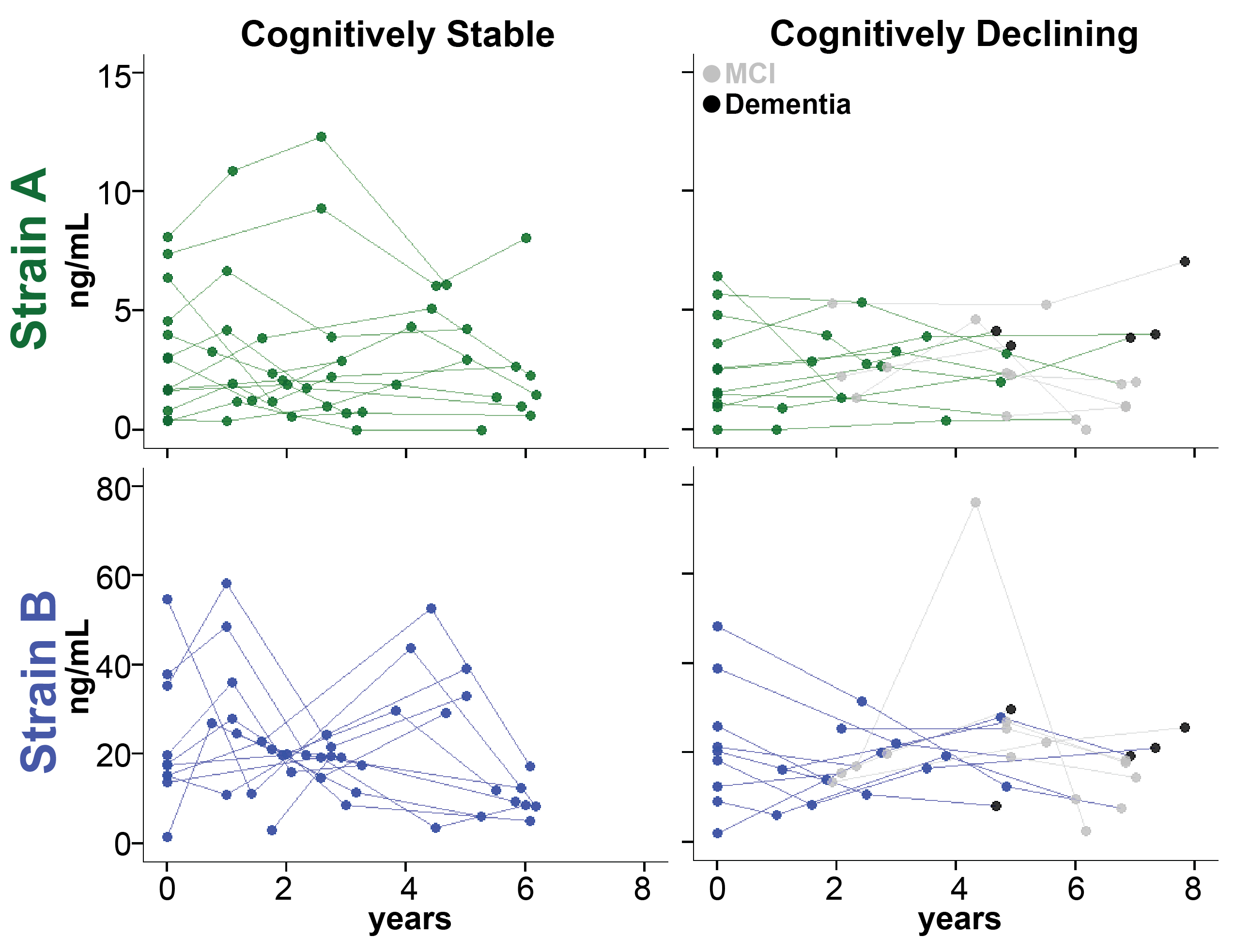
 **Supplementary Figure 8. aSyn strain plasma levels do not significantly change over time and do not mirror changes in cognition at the individual level.** Strain A and Strain B plasma levels were measured in 22 individuals with longitudinal plasma sampling. No differences were observed between individuals who did (n=11) vs. did not (n=11) develop cognitive decline. Groups were compared using a linear mixed-effects model adjusting for disease duration, baseline age, baseline cognition, and sex.

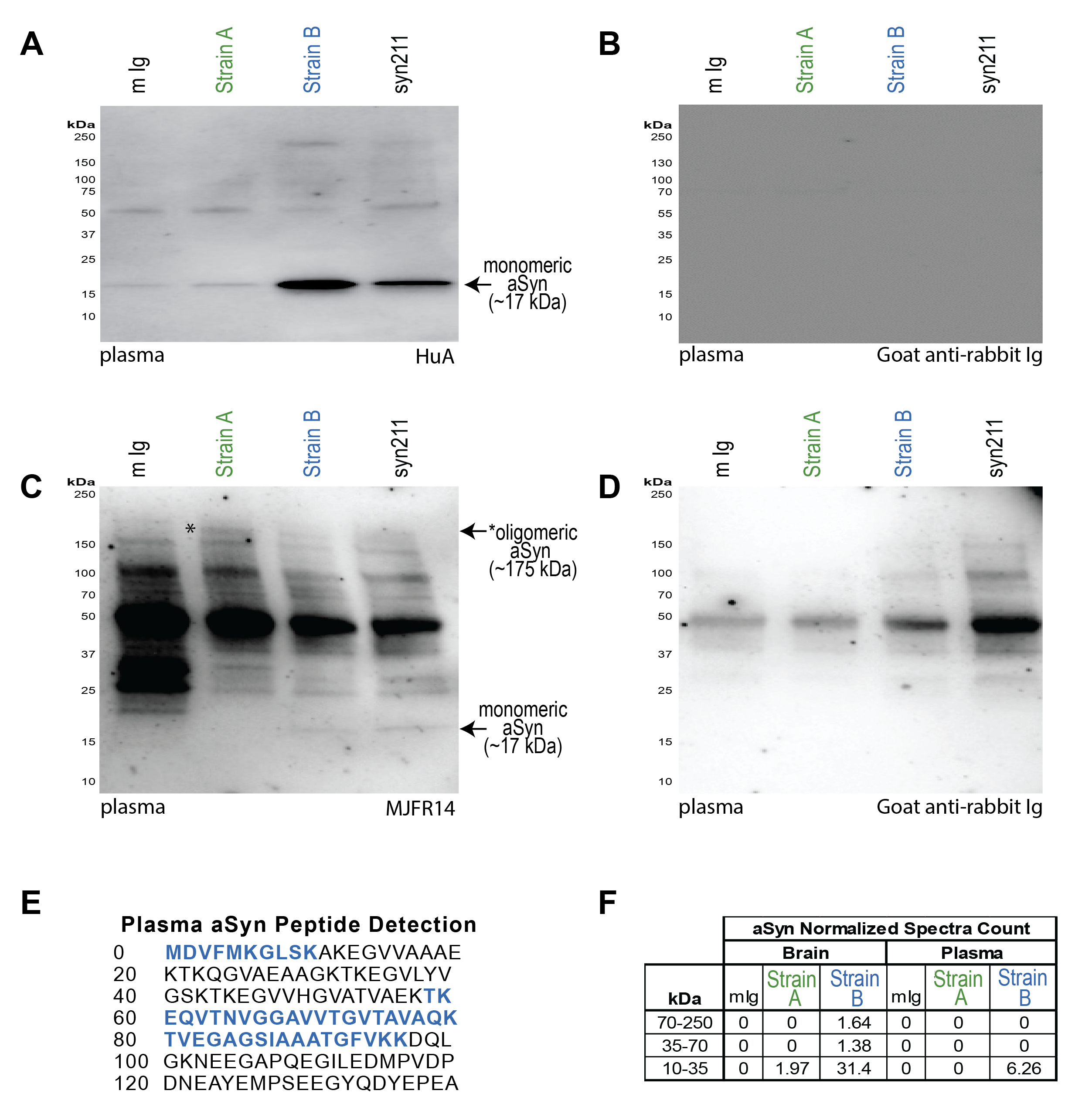

**Supplementary Figure 9. aSyn is detectabIe after immunoprecipitation with strain-selective antibodies.** Strain A, Strain B, or total aSyn (syn211) species were isolated by IP from pooled PD plasma or pooled PD brain caudate lysates. Western blot (WB) with HuA, a non-selective anti-human aSyn antibody, demonstrates robust signal at expected molecular weight of monomeric aSyn (~17 kDa) in Strain B and syn211 plasma IP with weak detection in Strain A plasma IP (A). WB with MJFR14, an anti-human aSyn antibody that prefers oligomeric species, demonstrates a ~175 kDa band (asterisk) that is enriched in the strain A IP (C) and not seen when probing with secondary antibody alone (B, D). These blots are representative of 5 and 3 independent plasma pools for HuA and MJFR14, respectively. aSyn was detected by mass spectrometry in both strain A and B IPs from brain but only in Strain B IP from plasma. Peptide sequences detected in Strain B plasma IP (E) and normalized spectra counts for 2D gel regions isolated and sent for proteomics identification are indicated (F).

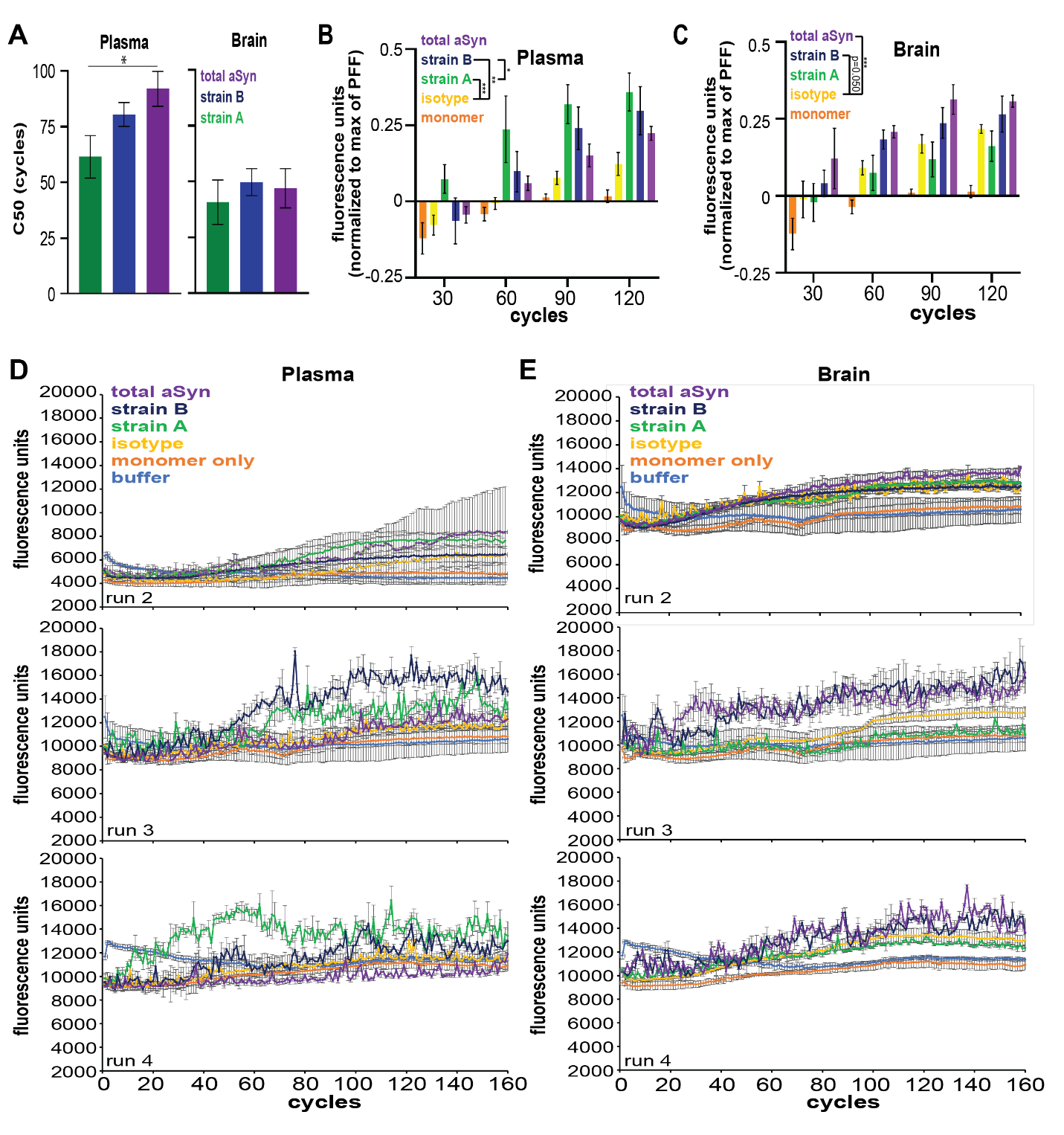

**Supplementary Figure 10. aSyn amplification curves from seed aggregation assays (SAA) using aSyn plasma and brain species in Parkinson’s disease from all performed runs.** A total of 4 SAA replicates with aSyn species immunoprecipitated with strain-selective and total aSyn (syn211) antibodies from plasma (n = 2 pools) and caudate brain lysates (n = 3 pools) with PFF, monomer, and buffer only controls were performed. C50 (cycle number at which half of maximum relative fluorescence is reached) was significantly lower with strain A seed compared to total aSyn only in plasma (**A**). In plasma, quantification of fluorescence by normalization to PFF maximum fluorescence demonstrated significantly increased fibrillization of aSyn with plasma aSyn species enriched by both Strain A and B antibodies (**B**). In brain, aSyn species enriched by syn211 (total) aSyn antibody significantly increased fibrillization and species enriched by Strain B antibody trended to significantly increased fibrillization (p = 0.052, **C**). SAA curves for additional replicates are also displayed (**D, E**). Background fluorescence based on buffer only condition was subtracted from all values. Error bars represent SEM. PFF = pre-formed fibrils. * p < 0.05, ** p < 0.01, *** p < 0.001.

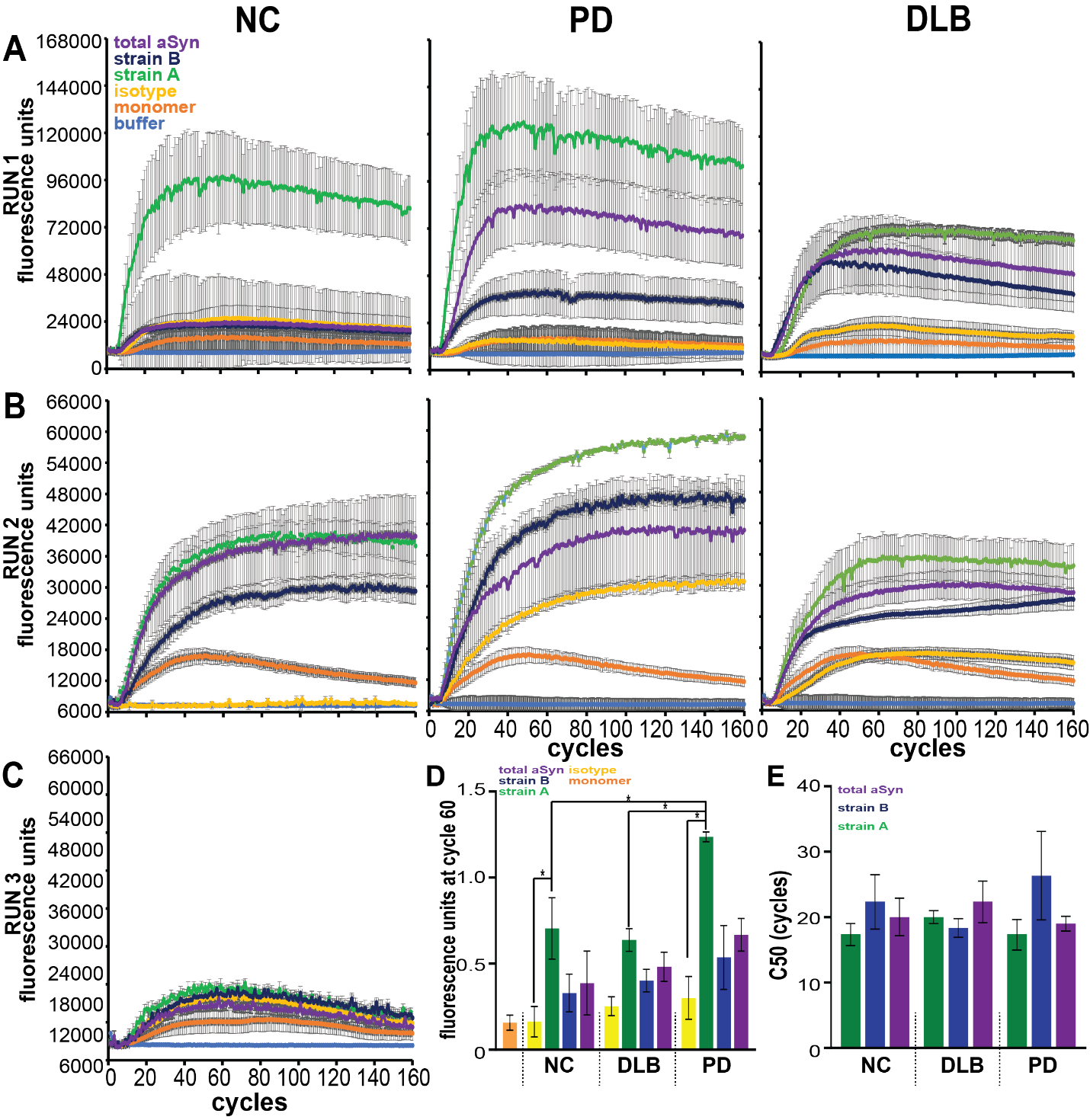
**Supplementary Figure 11. aSyn amplification curves from seed aggregation assays (SAA) using aSyn plasma species in normal controls (NC), dementia with Lewy bodies (DLB), and Parkinson’s disease (PD) from all performed runs.** A total of 3 SAA replicates with aSyn species immunoprecipitated with strain-selective and total aSyn (syn211) antibodies from plasma (n = 3 pools per disease group) with preformed fibril, monomer, and buffer only controls were performed. SAA curves for additional replicates are displayed (**A-C**). Strain A aSyn species significantly induced aggregation in SAA relative to Ig control when quantified by normalizing fluorescence at cycle 60 to maximum PFF fluorescence (**D**). Strain A aSyn species from PD plasma induced more significant aggregation relative to species from NC or DLB plasma. C50 (cycle number at which half of maximum relative fluorescence is reached) was unchanged across strain or disease group (**E**). Background fluorescence based on buffer only condition was subtracted from all values. Error bars represent SEM. * p < 0.05.

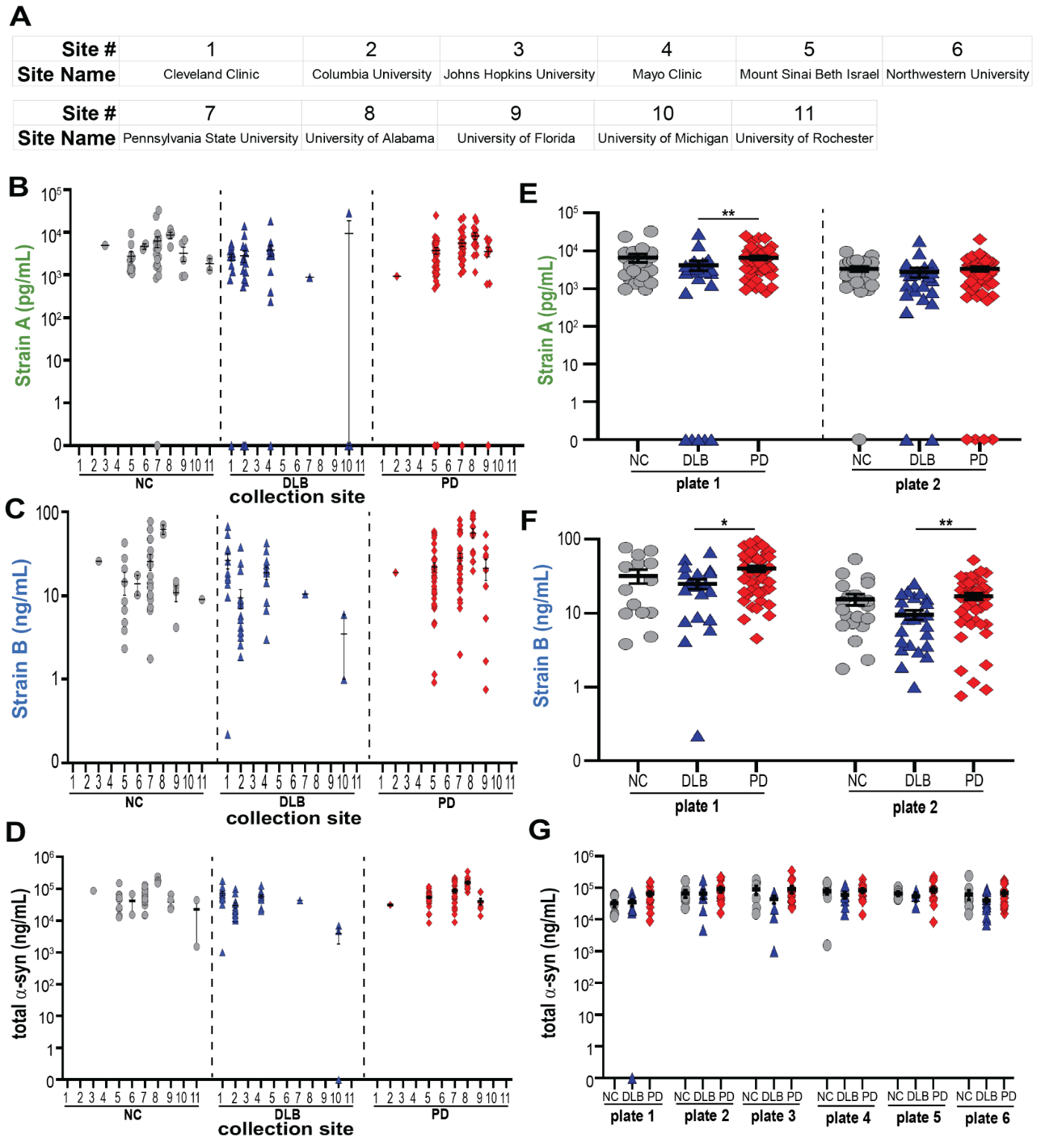
 **Supplementary Figure 12. PDBP cohort measures show site-to-site and plate-to-plate variability, demonstrating need for normalization.** The Parkinson’s Disease Biomarker Project (PDBP) cohort (n = 200) had plasma collected at 11 sites (**A**). Among sites, there was considerable site-to-site variability in plasma strain and total aSyn measurements (**B-D**). While each plate showed the same trends for PD vs. DLB, plate-to-plate variability was noted (**E-G**). For differences among groups, p-values (corrected for multiple comparisons for Kruskal-Wallis post-hoc test) for one-way ANOVA are reported in **E** and **F.** Error bars represent SEM. *p < 0.05, **p < 0.01. NC (grey circles) = normal controls, DLB (blue triangles) = dementia with Lewy bodies, PD (red diamonds) = Parkinson’s disease.

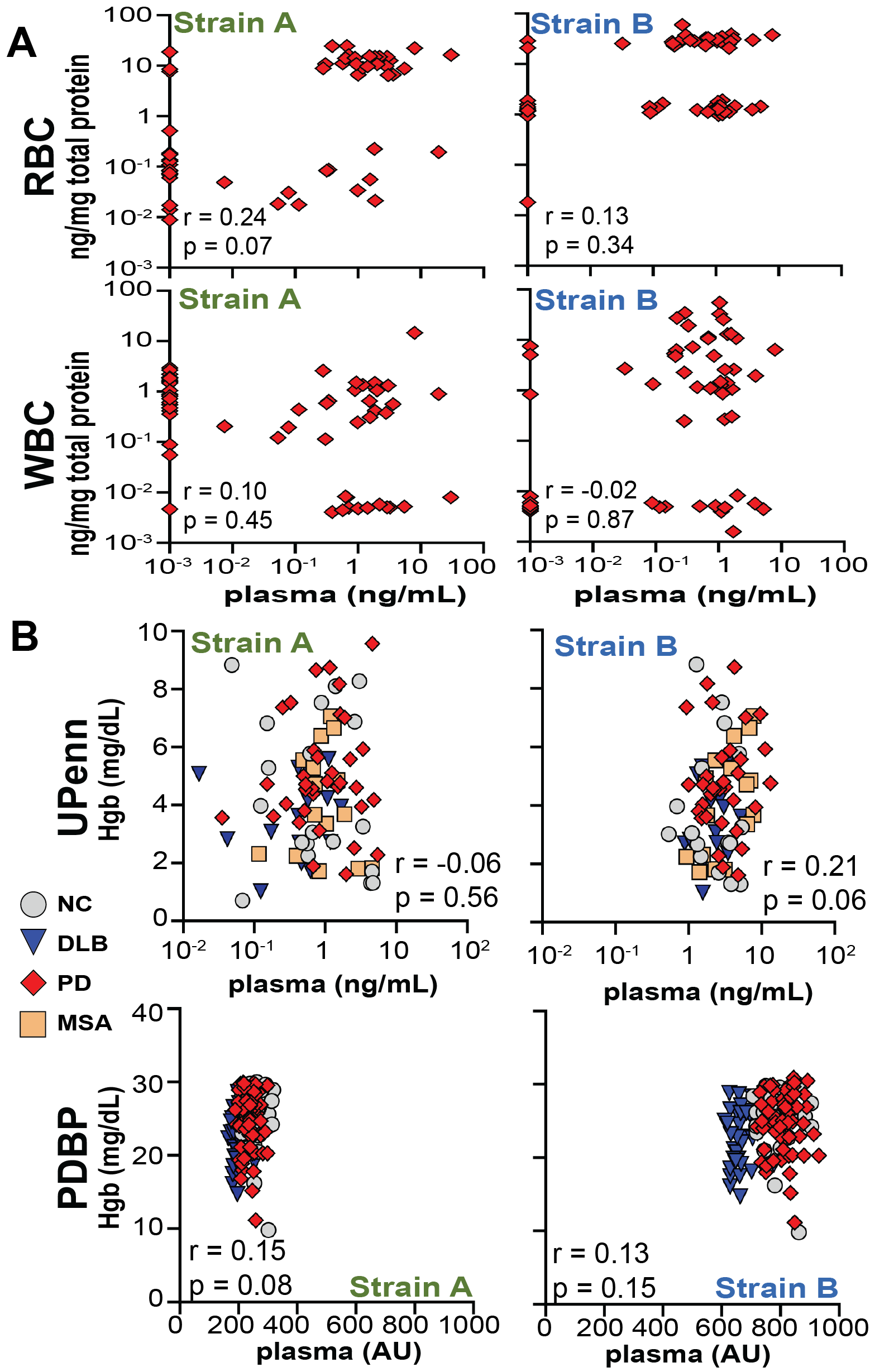

**Supplementary Figure 13. Plasma aSyn strain levels do not correlate with blood cell levels or hemolysis.** Levels of aSyn strains in plasma do not correlate (Pearson) with levels in matched white or red blood cell fractions (n = 65) from individuals with PD as measured by ELISA (**A**). In both the University of Pennsylvania (n = 235) and the PD Biomarker Project (n = 200) cohorts, plasma aSyn strain levels measured by ELISA did not correlate with hemoglobin levels (Hgb) measured by absorbance. Between group differences were assessed by one-way ANOVA with Kruskal-Wallis post-hoc testing (**B**). Pearson correlation r and p-values are displayed (**A-B**). * p < 0.05. AU = arbitrary units, NC = normal control, RBC = red blood cells, WBC = white blood cells.

| **#** | **Clinical Diagnosis** | **1º Path Dx** | **2º Path Dx** | **2º Path Dx Likelihood** | **Sex** | **Race** | **Ethnicity** | **Age at Death (yrs)** | **Disease Duration (yrs)** | **Age at Plasma Draw (yrs)** | **Interval Time (mo)** |
| --- | --- | --- | --- | --- | --- | --- | --- | --- | --- | --- | --- |
| 1 | DLB | LBD | AD | Int | M | White | Not Hispanic or Latino | 76 | 1 | 76 | 5 |
| 2 | DLB | LBD | AD | High | M | White | Not Hispanic or Latino | 81 | 2 | 81 | 2 |
| 3 | DLB | LBD | AD | Low | M | White | Not Hispanic or Latino | 74 | 1 | 73 | 14 |
| 4 | PD | LBD | AD | Low | F | White | Not Hispanic or Latino | 77 | 9 | 75 | 23 |
| 5 | PDD | LBD | AD | Low | M | White | Not Hispanic or Latino | 79 | 8 | 77 | 23 |
| 6 | PDD | LBD | AD | High | M | White | Not Hispanic or Latino | 74 | 2 | 73 | 9 |
| 7 | PDD | LBD | AD | Int | M | White | Not Hispanic or Latino | 85 | 11 | 84 | 5 |
| 8 | PDD | LBD | AD | Int | F | White | Not Hispanic or Latino | 91 | 11 | 89 | 21 |
| 9 | PDD | LBD | AD | Low | M | White | Not Hispanic or Latino | 70 | 8 | 68 | 16 |
| 10 | PDD | LBD | AD | Low | M | White | Not Hispanic or Latino | 81 | 18 | 80 | 15 |

**Supplementary Table 1. Samples for brain and matched plasma in Lewy boy disease (LBD).** Summary of characteristics of individuals selected for the brain immunohistochemistry and comparison of aSyn strain quantification in brain lysate and plasma by ELISA. Interval time refers to the number of months between the plasma sample collection date and the date of death. DLB = dementia with Lewy bodies, PD = Parkinson’s disease, PDD = PD dementia, Int = Intermediate, Path Dx = Pathological Diagnosis.

| # | **Clinical Diagnosis** | **Neuropathological Diagnosis** | **Sex** | **Race** | **Ethnicity** | **Age at Death (yrs)** | **Disease Duration (yrs)** |
| --- | --- | --- | --- | --- | --- | --- | --- |
| 1 | MSA-P | MSA | Female | White | Not Hispanic or Latino | 73 | 10 |
| 2 | MSA-P | MSA | Male | White | Unknown or Not Reported | 73 | 14 |
| 3 | MSA-C | MSA | Male | White | Not Hispanic or Latino | 75 | 6 |
| 4 | MSA-P | MSA | Male | White | Not Hispanic or Latino | 71 | 9 |
| 5 | MSA-P | MSA | Female | White | Not Hispanic or Latino | 74 | 4 |
| 6 | MSA-P | MSA | Male | White | Unknown or Not Reported | 76 | 5 |
| 7 | MSA-P | MSA | Male | White | Not Hispanic or Latino | 76 | 6 |
| 8 | MSA (unknown type) | MSA | Male | White | Not Hispanic or Latino | 79 | 5 |
| 9 | MSA-P | MSA | Male | White | Not Hispanic or Latino | 77 | 5 |
| 10 | MSA-C | MSA | Male | Unknown or Not Reported | Unknown or Not Reported | 72 | 7 |

**Supplementary Table 2. Samples for MSA brain immunohistochemistry.** Summary of characteristics of individuals with MSA selected for the brain immunohistochemistry. MSA = multiple systems atrophy, MSA-P = MSA parkinsonian type, MSA-C = MSA cerebellar type

| **#** | **Clinical Diagnosis** | **1º Path Dx** | **2º Path Dx** | **2º Path Dx Likelihood** | **Sex** | **Race** | **Ethnicity** | **Age at Death (yrs)** | **Disease Duration (yrs)** |
| --- | --- | --- | --- | --- | --- | --- | --- | --- | --- |
| 1 | NC | Pathological Aging | N/A | None | Male | White | Not Hispanic or Latino | 61 | N/A |
| 2 | NC | Normal | N/A | None | Male | White | Not Hispanic or Latino | 70 | N/A |
| 3 | NC | Pathological aging | N/A | None | Female | White | Not Hispanic or Latino | 70 | N/A |
| 4 | NC | Pathological Aging | N/A | None | Male | White | Unknown or Not Reported | 71 | N/A |
| 5 | NC | Normal | N/A | None | Male | Black or African American | Unknown or Not Reported | 66 | N/A |
| 6 | DLB | LBD | PART | N/A | Male | White | Not Hispanic or Latino | 73 | 10 |
| 7 | DLB | LBD | AD | Low | Male | White | Not Hispanic or Latino | 74 | 10 |
| 8 | DLB | LBD | AD | Intermediate | Male | White | Not Hispanic or Latino | 75 | 6 |
| 9 | DLB | LBD | AD | High | Male | White | Not Hispanic or Latino | 74 | 3 |
| 10 | DLB | LBD | AD | Low | Female | White | Not Hispanic or Latino | 85 | 8 |
| 11 | PD | LBD | AD | Low | Male | White | Not Hispanic or Latino | 75 | 21 |
| 12 | PD | LBD | None | N/A | Male | White | Unknown or Not Reported | 75 | 21 |
| 13 | PD | LBD | AD | Low | Male | White | Not Hispanic or Latino | 77 | 23 |
| 14 | PD | LBD | PART | N/A | Female | White | Not Hispanic or Latino | 80 | 10 |
| 15 | PD | LBD | AD | Low | Female | White | Not Hispanic or Latino | 77 | 10 |

**Supplementary Table 3. ELISA and Western Blot Comparison Samples.** Characteristics of individuals selected for the comparison of aSyn strains by ELISA and Western Blot. Path Dx = Pathological Diagnosis, NC = normal control, DLB = dementia with Lewy bodies, LBD = Lewy Body Disease, PD = Parkinson’s disease, AD = Alzheimer’s disease, N/A = not applicable.

| **N** | **Race** | **Sex** | **Clinical Diagnosis** | **UPDRS part III** | **MOCA** | **Age at**  **Plasma Collection (yrs)** | **Age at**  **CSF Collection (yrs)** | **Interval time (mo)** |
| --- | --- | --- | --- | --- | --- | --- | --- | --- |
| 44 | White 97.8%  Black 2.2 % | Male 60%  Female 40% | PD 97.7%  DLB 2.3% | 21.5 ± 1.6 | 26.0 ± 0.48 | 68.0 ± 1.1 | 68.0 ± 1.1 | 0.87 ± 0.25 |

**Supplemental Table 4. Matched CSF and Plasma Samples.** Summary of characteristics of individuals selected for the comparison of CSF and Plasma values on ELISA listed as mean ± standard error of the mean or frequency (%). Interval time refers to the number of months between the plasma sample date and the CSF sample date.

|  |  | **UPenn** | **PDBP** | **p-value** |
| --- | --- | --- | --- | --- |
| **NC** | N | 49 | 50 |  |
|  | Age (years) | 72.5 ± 0.18 | 68.2 ± 0.15 | 0.10 |
|  | Sex | Male 51% Female 49% | Male 76% Female 24% | ***0.0001*** |
|  | Race | White 71% Black 27% Multiracial 2% | White 90% Black 4% Native American 4% Not Reported 2% | ***0.0001*** |
|  | MOCA | 27.8 ± 0.07 | 27.8 ± 0.08 | 0.5 |
| **AD** | N | 46 |  |  |
|  | Age (years) | 74.6 ± 0.19 |  |  |
|  | Sex | Male 46% Female 54% |  |  |
|  | Race | White 81% Black 15% Multiracial 4% |  |  |
|  | Disease Duration (years) | 4.5 ± 0.06 |  |  |
|  | MOCA | 20 ± 0.26 |  |  |
| **DLB** | N | 25 | 50 |  |
|  | Age (years) | 69.2 ± 0.35 | 69.2 ± 0.12 | 0.74 |
|  | Sex | Male 51% Female 49% | Male 80% Female 20% | ***0.0001*** |
|  | Race | White 92% Black 4% Multiracial 4% | White 96% Black 2% Native American 2% | 0.12 |
|  | Disease Duration (years) | 4.1 ± 0.11 | 3.2 ± 0.10 | ***0.005*** |
|  | UPDRS part III (Motor) | 30.4 ± 1.2 | 31.8 ± 0.31 | 0.64 |
|  | MOCA | 14.3 ± 0.53 | 19.3 ± 0.14 | ***0.03*** |
| **PD** | N | 115 | 100 |  |
|  | Age | 66.8 ± 0.07 | 68.2 ± 0.06 | 0.09 |
|  | Sex | Male 69% Female 31% | Male 78% Female 22% | ***0.03*** |
|  | Race | White 96% Black 4% | White 98% Native American 1% Other 1% | 0.15 |
|  | Disease Duration | 5.0 ± 0.04 | 7.3 ± 0.11 | 0.60 |
|  | UPDRS part III (Motor) | 22.1 ± 0.09 | 25.9 ± 0.19 | 0.05 |
|  | MOCA | 24.9 ± 0.05 | 25.3 ± 0.06 | 0.87 |

**Supplementary Table 5. UPenn and PDBP Plasma Cohorts.** Characteristics of UPenn and PDBP cohorts used for plasma and clinical analyses listed as mean ± standard error of the mean or frequency (%). P-values represent Mann-Whitney tests and chi-square tests for continuous variables or proportions, respectively.

|  | **NC** | **DLB** | **PD** | **MSA** | **p-value** |
| --- | --- | --- | --- | --- | --- |
| **N** | 20 | 20 | 43 | 18 |  |
| **Sex** | Male (40%), Female (60%) | Male (45%), Female (55%) | Male (58.1%), Female (41.9%) | Male (38.9%), Female (61.1%) | 0.40 |
| **Race** | White (85%), Black (15%) | White (95%), Multiracial (5%) | White (95%), Black (5%) | White (89%), Unknown (11%) | ***0.03*** |
| **Ethnicity** | Not Hispanic or Latino (100%) | Not Hispanic or Latino (100%) | Not Hispanic or Latino (100%) | Hispanic (5.6%), Not Hispanic or Latino (94.4%) | 0.18 |
| **Age** | 65.6 ± 1.7 | 67.2 ± 1.7 | 67.7 ± 0.9 | 64.9 ± 2.2 | 0.48 |
| **Disease Duration** | N/A | 3.8 ± 0.7 | 9.3 ± 0.8 | 5.1 ± 1.1 | ***<0.001*** |
| **UPDRS part III** | N/A | 27.5 ± 4.9 | 22.0 ± 1.8 | N.D. | 0.37 |
| **MOCA** | N.D. | 17.2 ± 3.6 | 25.6 ± 0.7 | 22.3 ± 1.2 | 0.003 |

**Supplementary Table 6. Characteristics of Plasma Samples for comparison of aSyn strains in plasma from normal controls (NC), dementia with Lewy bodies (DLB), Parkinson’s disease, and Multiple Systems Atrophy (MSA).** Characteristics of plasma samples listed as mean ± standard error of the mean or frequency (%). P-values represent Kruskal-Wallis tests and Fisher exact tests for continuous variables or proportions, respectively. N/A = not applicable, N.D. = no data.

| **Clinical Diagnosis** | **N** | **Race** | **Sex** | **Age (yrs)** | **Disease Duration (yrs)** |
| --- | --- | --- | --- | --- | --- |
| Healthy Control | 30 | 80.0% White  20.0% Black or African American | 66.7% Male  33.3% Female | 68.8 ± 1.3 | - |
| Alzheimer's Disease | 30 | 93.3% White  3.3% Black or African American  3.3% Asian | 63.3% Male  36.7% Female | 69.2 ± 1.3 | 4.3 ± 0.7 |
| Dementia with Lewy bodies | 30 | 93.3% White  3.3% Black or African American  3.3% Multiracial | 66.7% Male  33.3% Female | 68 ± 1.4 | 3.7 ± 0.7 |
| Parkinson’s Disease | 88 | 94.3% White  2.3% Black or African American  2.3% Asian  1.1% Not Reported | 67.0% Male  33.0% Female | 68.9 ± 0.8 | 7.2 ± 0.5 |

**Supplementary Table 7. Total α-synuclein ELISA Samples.** Characteristics of individuals selected for analysis on total α-synuclein ELISA, listed as mean ± standard error of the mean or frequency (%).

| **N** | **Clinical Diagnosis at Baseline** | **Race** | **Sex** | **Age at Baseline (yrs)** | **Disease Duration at Baseline (yrs)** | **# of Visits** | **Follow-up Duration (years)** | **Baseline UPDRS part III** | **Baseline Age-Adjusted DRS** |
| --- | --- | --- | --- | --- | --- | --- | --- | --- | --- |
| 95 | PD 75.7% PD-MCI 24.2% | White 96.8% Black 3.1% | Male 68% Female 32% | 66.7 ± 0.8 | 5.1 ± 0.4 | 5.1 ± 0.2 | 5.2 ± 0.3 | 20.9 ± 1.0 | 10.8 ± 0.3 |

**Supplementary Table 8. Samples for Motor and Cognitive Linear Mixed-Effects Model.** Characteristics of individuals selected for motor and cognitive linear mixed-effects model listed as mean ± standard error of the mean or frequency (%). Cognitive status established by consensus diagnosis.

| **#** | **Sex** | **Age at BL (yrs)** | **Disease Duration Visit 1** | **Cognitive Status** | **Disease Duration Visit 2** | **Cognitive Status** | **Disease Duration Visit 3** | **Cognitive Status** | **Disease Duration Visit 4** | **Cognitive Status** |
| --- | --- | --- | --- | --- | --- | --- | --- | --- | --- | --- |
| 1 | M | 68 | 5 | Normal | 7 | Normal | 9 | Normal | 13 | Dementia |
| 2 | M | 73 | 8 | Normal | 11 | Normal | 13 | MCI | 15 | Dementia |
| 3 | F | 75 | 7 | Normal | 9 | Normal | 11 | Normal | 13 | MCI |
| 4 | F | 69 | 21 | Normal | 23 | Normal | 24 | Normal | 26 | Dementia |
| 5 | M | 64 | 4 | Normal | 7 | Normal | 9 | Normal | 10 | Normal |
| 6 | M | 64 | 4 | Normal | 5 | Normal | 9 | MCI | 11 | MCI |
| 7 | M | 63 | 4 | Normal | 6 | Normal | 8 | Normal | 10 | Normal |
| 8 | F | 60 | 5 | Normal | 6 | Normal | 8 | Normal | 10 | Normal |
| 9 | F | 59 | 3 | Normal | 4 | Normal | 5 | Normal | 7 | Normal |
| 10 | M | 54 | 5 | Normal | 8 | MCI | 10 | MCI | 12 | MCI |
| 11 | M | 70 | 10 | Normal | 12 | MCI | 16 | MCI | 18 | Dementia |
| 12 | F | 70 | 1 | Normal | 3 | Normal | 4 | Normal | 7 | Normal |
| 13 | F | 73 | 7 | Normal | 9 | Normal | 10 | Normal | 13 | Normal |
| 14 | F | 72 | 12 | Normal | 14 | Normal | 16 | Normal | 18 | Normal |
| 15 | M | 62 | 1 | Normal | 2 | Normal | 5 | Normal | 7 | MCI |
| 16 | M | 72 | 2 | Normal | 3 | Normal | 6 | Normal | 8 | Normal |
| 17 | M | 64 | 4 | Normal | 5 | Normal | 7 | Normal | 9 | Normal |
| 18 | F | 63 | 12 | Normal | 14 | MCI | 15 | MCI | 17 | Dementia |
| 19 | F | 65 | 3 | Normal | 7 | Normal | 8 | MCI | 10 | MCI |
| 20 | M | 69 | 6 | Normal | 8 | Normal | 9 | Normal | 12 | Normal |
| 21 | M | 71 | 9 | Normal | 12 | Normal | 14 | MCI | 16 | MCI |
| 22 | M | 75 | 5 | Normal | 6 | Normal | 8 | Normal | 10 | Normal |

**Supplemental Table 9. Longitudinal Plasma ELISA Samples.** Characteristics of individuals selected for the longitudinal plasma cohort whose aSyn strain levels were measured on the ELISA. All individuals were of white race. Disease duration reported in years. Cognitive status established by consensus diagnosis.

| **Pool** | **N** | **Clinical Diagnosis** | **Sex** | **Race** | **Age at Sample (yrs)** | **Disease Duration (yrs)** |
| --- | --- | --- | --- | --- | --- | --- |
| 1 | 43 | PD (93%), PD-MCI (7%) | Female (41%),  Male (59%) | White (87.5%), Black (5%),  Unknown (5%),  Asian (2.5%) | 67.2 ± 1.2 | 9.4 ± 1.1 |
| 2 | 119 | PD (100%) | Female (42%),  Male (58%) | White (89%),  Black (5%),  Asian (3%),  Other (5%) | 68.9 ± 0.9 | 5.9 ± 0.7 |
| 3 | 51 | PD (100%) | Female (32%), Male (68%) | White (97.9%), Black (2.1%), | 68.1 ± 1.6 | 11.4 ± 10.9 |
| 4 | 52 | PD (100%) | Female (33%), Male (67%) | White (93.9%), Black (4.1%),  Other (2.0) | 68.6 ± 1.3 | 11.7 ± 1.4 |
| 5 | 52 | PD (100%) | Female (35%), Male (65%) | White (95.8%), Black (2.1%),  Asian (2.1%) | 68.4 ± 1.6 | 12.8 ± 1.3 |
| 6 | 50 | DLB (100%) | Female (28%),  Male (72%) | White (91.5%), Black (6.4%), Other (2.1%) | 69.6 ± 1.2 | 4.2 ± 0.5 |
| 7 | 60 | DLB (100%) | Female (20%),  Male (80%) | White (88.3%), Black (5%), Other (6.7%) | 67.1 ± 0.9 | 5.0 ± 0.5 |
| 8 | 60 | DLB (100%) | Female (15%),  Male (85%) | White (85%), Black (8.3%), Other (6.7%) | 67.2 ± 0.9 | 5.7 ± 0.4 |
| 9 | 54 | NC (100%) | Female (32%), Male (68%) | White (66.7%), Black (24.0%),  Asian (1.9%), Other (7.4%) | 68.4 ± 1.0 | N/A |
| 10 | 64 | NC (100%) | Female (61%),  Male (39%) | White (67.2%), Black (28.1%),  Asian (1.7%), Other (3.0%) | 72.5 ± 1.0 | N/A |
| 11 | 60 | NC (100%) | Female (75%),  Male (25%) | White (61.7%), Black (31.7%),  Asian (1.7%), Other (4.9%) | 73.9 ± 1.0 | N/A |

**Supplementary Table 10. Pooled Plasma Immunoprecipitation Samples.** Characteristics of individuals selected for pooled plasma for immunoprecipitation listed as mean ± standard error of the mean or frequency (%).

| **Pool** | **N** | **Clinical Diagnosis** | **Sex** | **Race** | **Age at Death (yrs)** | **Disease Duration (yrs)** |
| --- | --- | --- | --- | --- | --- | --- |
| 1 | 10 | DLB (30%), PD (10%), PDD (60%) | Female (20%), Male (80%) | White (100%) | 78.8 ± 1.9 | 7.1 ± 1.8 |
| 2 | 8 | PDD (50%),  PD (37.5%),  PD-MCI (12.5%) | Female (37.5%), Male (67.5%) | White (100%) | 79.8 ± 0.8 | 12.1 ± 2.2 |
| 3 | 17 | DLB (18%),  PD (18%),  PDD (53%),  PD-MCI (11%) | Female (71%), Male (29%) | White (100%) | 79.9 ± 1.3 | 13.9 ± 1.7 |

**Supplementary Table 11. Pooled Brain Lysate Immunoprecipitation Samples.** Characteristics of individuals selected for pooled brain caudate lysate for immunoprecipitation listed as mean ± standard error of the mean or frequency (%).

| **plasma** | **Strain A (ng/mL)** | **Fold. Enrich.** | **Strain B (ng/mL)** | **Fold. Enrich.** | **Total aSyn (ng/mL)** | **Fold. Enrich.** |
| --- | --- | --- | --- | --- | --- | --- |
| **input** | 1.29 ± 0.37 |  | 10 ± 2.9 |  | 52 ± 22 |  |
| **Neg control IP** | 0.18 ± 0.03 |  | 0.0 ± 0.0 |  | 0.44 ± 0.16 |  |
| **Strain A IP** | 0.55 ± 0.07 | 3.4 ± 1.1 | 0.3 ± 0.2 | 11.7 ± 9.3 | 1.4 ± 0.9 | 20 ± 18 |
| **Strain B IP** | 0.61 ± 0.03 | 3.7 ± 1.0 | 7.3 ± 3.4 | 221 ± 128 | 11.3 ± 5.1 | 241 ± 219 |
| **Total aSyn IP** | 0.47 ± 0.25 | 2.2 ± 1.0 | 4.8 ± 3.1 | 227 ± 208 | 9.4 ± 2.5 | 302 ± 283 |
| **brain** | **Strain A (ng/mL)** | **Fold. Enrich.** | **Strain B (ng/mL)** | **Fold. Enrich.** | **Total aSyn (ng/mL)** | **Fold. Enrich.** |
| **input** | 885 ± 23 |  | 3196 ± 579 |  | 5413 ± 165 |  |
| **Neg control IP** | 3.8 ± 3.8 |  | 0.0 ± 0.0 |  | 27 ± 25 |  |
| **Strain A IP** | 5.3 ± 5 | 3.6 ± 2.3 | 0.9 ± 0.7 | 54 ± 40 | 44 ± 20 | 125 ± 90 |
| **Strain B IP** | 33 ± 5.7 | 274 ± 270 | 163 ± 41 | 9803 ± 2433 | 670 ± 8.0 | 1520 ± 790 |
| **Total aSyn IP** | 13 ± 5.7 | 46 ± 44 | 49 ± 23 | 2942 ± 1377 | 630 ± 22 | 1460 ± 780 |

**Supplementary Table 12. Concentration of aSyn Isolated by IP from Plasma and Brain with Strain-selective Antibodies.** Concentration of aSyn in eluates isolated by strain-selective or total aSyn antibody IP are represented by the mean ± standard error of the mean of three brain IPs and six plasma IPs. The negative control IP was performed with isotype-matched IgG antibody. Fold enrichment calculated based on half of lower limit of detection if negative (neg) control IP ELISA concentration was below limit of detection. IP = immunoprecipitation.

| **N** | **Clinical Diagnosis** | **Race** | **Sex** | **Age at Sample Collection (years)** | **Disease Duration (years)** |
| --- | --- | --- | --- | --- | --- |
| 65 | PD (100 %) | White (81.5 %), Black (16.9 %), Asian (1.6%) | Male (50.8 %), Female (49.2%) | 68.9 ± 1.1 | 6.9 ± 1.0 |

**Supplementary Table 13. Parkinson’s disease (PD) cohort for matched blood cell and plasma samples.** Characteristics of PD cohorts used for measurement of aSyn strain levels in matched blood cell and plasma samples listed as mean ± standard error of the mean or frequency (%).

| **Reagent** | **Source** |
| --- | --- |
| 7015 (Strain A) mAb | Center for Neurodegeneration Research, University of Pennsylvania, Philadelphia, PA, USA |
| 9027 (Strain B) mAb | Center for Neurodegeneration Research, University of Pennsylvania, Philadelphia, PA, USA |
| Anti-total-aSyn (MJFR1) mAb | Abcam ab138501, Cambridge, UK |
| Anti-total-aSyn (syn211) mAb | Santa Cruz Biotechnology sc-12767, Dallas, TX, USA |
| Anti-tau (AT8) mAb | Thermofisher **MN1020, Waltham, MA, USA** |
| Anti-Aβ (NAB228) mAb | Center for Neurodegeneration Research, University of Pennsylvania, Philadelphia, PA, USA |
| Anti-total-aSyn (HuA) mAb | Center for Neurodegeneration Research, University of Pennsylvania, Philadelphia, PA, USA |
| Anti-total-aSyn (MJFR14) mAb | Abcam ab227047, Cambridge, UK |
| Anti-phosphorylated aSyn (MJF-R13) mAb | Abcam ab168381, Cambridge, UK |
| EDTA Vacutainer | BD 367863, Franklin Lakes, NJ, USA |
| Block ACE | Bio-Rad BUF029, Hercules, CA, USA |
| Chemiluminescence-based LEGEND MAX ™️ Human α-Synuclein ELISA Kit | Biolegend 844101, San Diego, CA, USA |
| Colorimetric LEGEND MAX ™️ Human α-Synuclein ELISA Kit | Biolegend 448607, San Diego, CA, USA |
| TMB Substrate Solution | Thermofisher N301, Waltham, MA, USA |
| Goat anti-mouse-HRP | Jackson Immunoresearch Laboratories 115-035-062, West Grove, PA, USA |
| Anti-GAPDH | Advanced ImmunoChemical Inc, Sigma, MAB6C5, Long Beach, CA, USA |
| Diagenode Protein Extraction Beads | Diagenode C20000021, Denville, NJ, USA |
| Streptavidin-HRP | Jackson Immunoresearch Laboratories 016-030-084, West Grove, PA, USA |
| PMSF Protease Inhibitor | Thermofisher 36978, Waltham, MA, USA |
| Diagenode Bioruptor Plus | Diagenode B01020001, Denville, NJ, USA |
| 4-20% Criterion TGX Midi Protean gels | Bio-Rad 5671094, Hercules, CA, USA |
| nitrocellulose 0.45 µm membrane | Bio-Rad 1620113, Hercules, CA, USA |
| Semi-dry TransBlot Turbo transfer apparatus | Bio-Rad 1703848, Hercules, CA, USA |
| Westernbright ECL reagent | Advansta K-12045, San Jose, CA, USA |
| Sirius ECL reagent | Advansta K-12043, San Jose, CA, USA |
| Chemidoc system | Bio-Rad 12003154, Hercules, CA, USA |
| Aminolink kit | Thermofisher #44890, Waltham, MA, USA |
| 96-well clear bottom black-well plate | Thermofisher #3165305, Waltham, MA, USA |
| Silica beads | OPS Diagnostics PFMB 800-100, Lebanon, NJ, USA |
| Pepstatin | Sigma P4265, St. Louis, MO, USA |
| Leupeptin | Sigma L2023, St. Louis, MO, USA |
| TPCK | Sigma T4376, St. Louis, MO, USA |
| TLCK | Sigma T7254, St. Louis, MO, USA |
| Trypsin Inhibitor | Sigma T9003, St. Louis, MO, USA |
| Polymorphoprep | CosmoBioUSA Catalog No:AXS-1114683, Carlbad, CA 90210 |
| Red blood cell lysis buffer | Sigma 11814389001, St. Louis, MO, USA |

**Supplementary Table 14.** Antibodies and reagents used in this manuscript.
